# The CIViC knowledge model and standard operating procedures for curation and clinical interpretation of variants in cancer

**DOI:** 10.1101/700179

**Authors:** Arpad M. Danos, Kilannin Krysiak, Erica K. Barnell, Adam C. Coffman, Joshua F. McMichael, Susanna Kiwala, Nicholas C. Spies, Lana M. Sheta, Shahil P. Pema, Lynzey Kujan, Kaitlin A. Clark, Amber Z. Wollam, Shruti Rao, Deborah I. Ritter, Dmitriy Sonkin, Gordana Raca, Raymond H. Kim, Alex H. Wagner, Subha Madhavan, Malachi Griffith, Obi L. Griffith

**Author notes:** These authors contributed equally to this work (Arpad M. Danos, Kilannin Krysiak, Erica K. Barnell). Corresponding authors: Malachi Griffith; Obi Griffith.

## Abstract

Manually curated variant knowledgebases and their associated knowledge models are serving an increasingly important role in distributing and interpreting variants in cancer. These knowledgebases vary in their level of public accessibility, and the complexity of the models used to capture clinical knowledge. CIViC (Clinical Interpretations of Variants in Cancer - www.civicdb.org) is a fully open, free-to-use cancer variant interpretation knowledgebase that incorporates highly detailed curation of evidence obtained from peer-reviewed publications. Currently, the CIViC knowledge model consists of four main components: Genes, Variants, Evidence Items, and Assertions. Each component has an associated knowledge model and methods for curation. Gene and Variant data contextualize the genomic region(s) involved in the clinical statement. Evidence Items provide structured associations between variants and their clinically predictive/therapeutic, prognostic, diagnostic, predisposing, and functional implications. Finally, CIViC Assertions summarize collections of CIViC Evidence Items for a specific Disease, Variant, and Clinical Significance with incorporation of clinical and technical guidelines. Here we present the CIViC knowledge model, curation standard operating procedures, and detailed examples to support community-driven curation of cancer variants.

## Introduction

Expansion of pan-cancer sequencing efforts in research and clinical settings has led to a rapid increase in the number of variants that require clinical annotation [1–5]. Computational and manual requirements for variant identification and interpretation has been shown to hinder the development of optimal treatment protocols for patients [6,7]. These issues highlight the need for normalized clinical classification and representation of relevant variants, as well as open distribution of a standardized cancer variant knowledgebase [8,9]. The Clinical Interpretations of Variants in Cancer (CIViC) knowledgebase (www.civicdb.org) was developed to address the challenges outlined above [10].

CIViC is a fully open, free-to-use knowledgebase, which incorporates clinical evidence associated with a biomedical publication. Evidence supporting specific clinical interpretations is gathered via crowdsourced curation followed by expert review and moderation. All submissions, revisions, moderations, and comments on CIViC entries are tracked and displayed through the CIViC web interface, providing transparency and clear provenance of all content in the knowledgebase.

Here we describe the CIViC knowledge model, and provide a standard operating procedure for clinical interpretation of variants in cancer using the CIViC platform. We first provide details on how the knowledge model permits consumption as well as curation of clinical information. We then describe each of the four components (Genes, Variants, Evidence Items, and Assertions) by outlining associated features, detailing methods for moderation, and providing curation examples. Further details on the CIViC knowledge model, standards and guidelines for curation and moderation, and details on the CIViC project are available in the CIViC help documents (https://civicdb.org/help/introduction).

## The CIViC knowledge model and key components

### The CIViC knowledge model for clinical variants

The CIViC knowledgebase was built to permit both consumption (i.e., searching, browsing, and downloading) of existing entries as well as curation of new content. The knowledgebase has been organized into a four-level hierarchy: Genes, Variants, Evidence Items, and Assertions (**Figure 1A**). Each level has its own knowledge model. All data created using these knowledge models are available through a web interface (www.civicdb.org) and an application programming interface (API, https://griffithlab.github.io/civic-api-docs).

**Figure 1.**
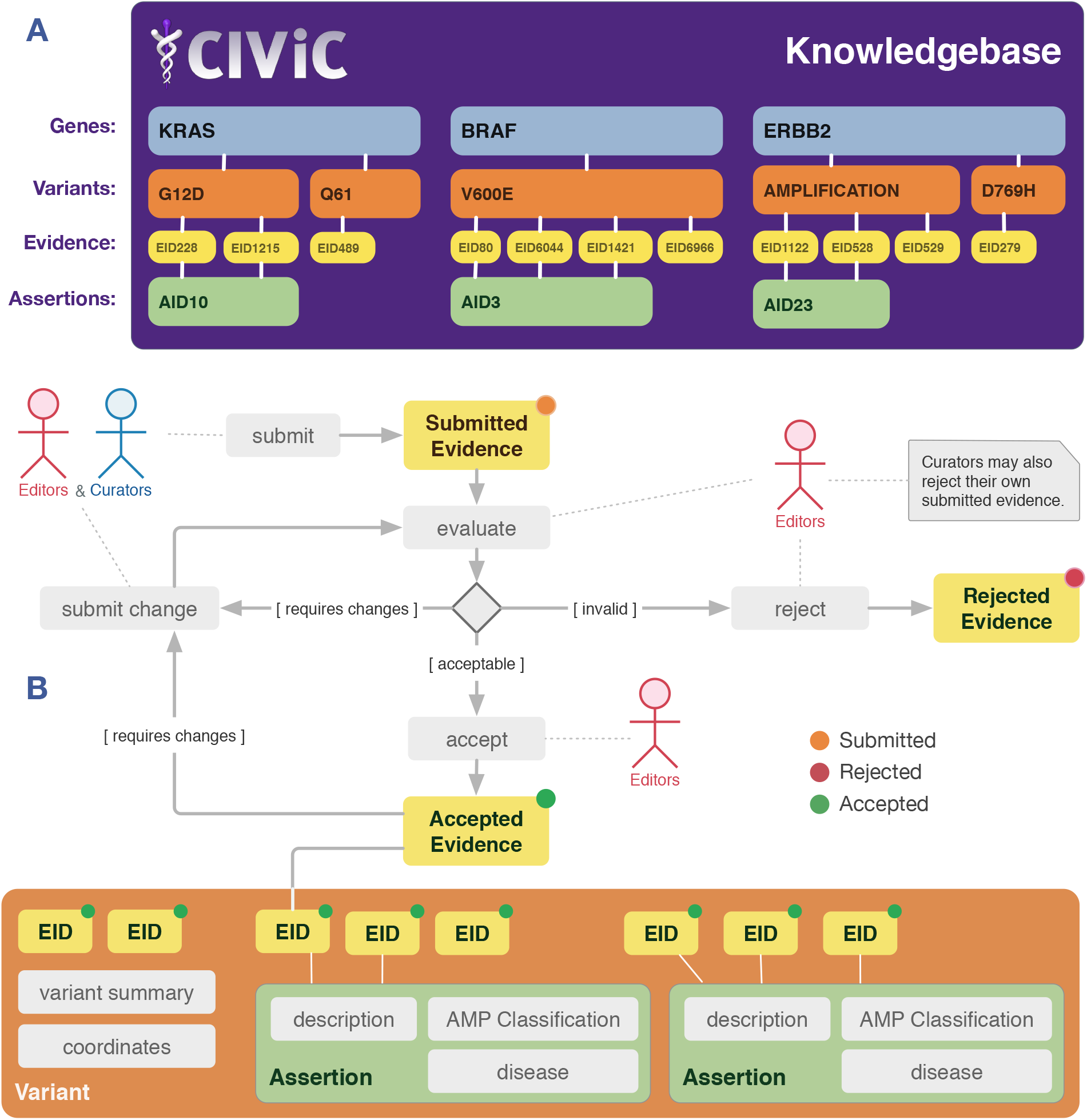
Overview of the CIViC knowledge model for the exploration of existing data (i.e., searching and browsing) and content curation. **A)** The CIViC knowledge model consists of four interconnected levels that contribute to the content within CIViC: Genes (blue), Variants (orange), Evidence (yellow), and Assertions (green). Each broadly defined variant is associated with a single gene but can have many lines of evidence linking it to clinical relevance. **B)** CIViC curation typically begins with the submission of an Evidence Item. Creation of an Evidence Item will automatically generate Gene and Variant records in the knowledgebase if they do not already exist. Once submitted, the Evidence Item undergoes evaluation by a panel of expert editors and (if necessary) revision with ultimate rejection or acceptance. Accepted Evidence Items can be used to build Assertions, which are visualized at the Variant-level. Similar cycles of curation and moderation are employed for all curatable entities in CIViC (e.g. Variant Summaries, Coordinates, Assertions).

For data generation, curators can add or edit curated content at each level (**Figure 1B**). Adding content involves submitting new Evidence Items or Assertions that subsequently undergo revision and review. Editing content involves adding or revising the clinical summary and/or its associated features. Once changes are made within the CIViC database, the additions/revisions become visible (depending on user display preferences). However, the change will be listed as a “submitted” (i.e., pending) until it is accepted by an editor. Users may reject (but not accept) their own submissions/revisions.

CIViC curators should avoid directly copying phrases from original sources (including abstracts) for summaries, statements, and comments. This practice prevents plagiarism and copyright infringement for articles with limited public access. Suggested revisions should always include a comment, providing rationale for the change. This allows editors to better understand the changes being proposed and facilitates acceptance or further modification. Additionally, if a curator finds inaccuracies or inconsistencies in the database, they should flag such entities to assist editors in rectifying curation issues. Other useful curation features that are found throughout CIViC are described in **Supplementary Table 1**.

### Structure and curation of the Gene knowledge model

#### Structure of the Gene knowledge model

The **Gene** knowledge model provides summarized genomic context for all CIViC variants contained by the gene. Gene features include: Gene Name, Gene Summary, external link to The Drug Gene Interaction Database [11–13], useful citations on the overall clinical relevance of the gene, and link-out details from MyGene.info [14] (**Figure 2A**). For a gene record to become visible, it must be associated with at least one Evidence Item that describes a variant contained by the gene.

**Figure 2.**
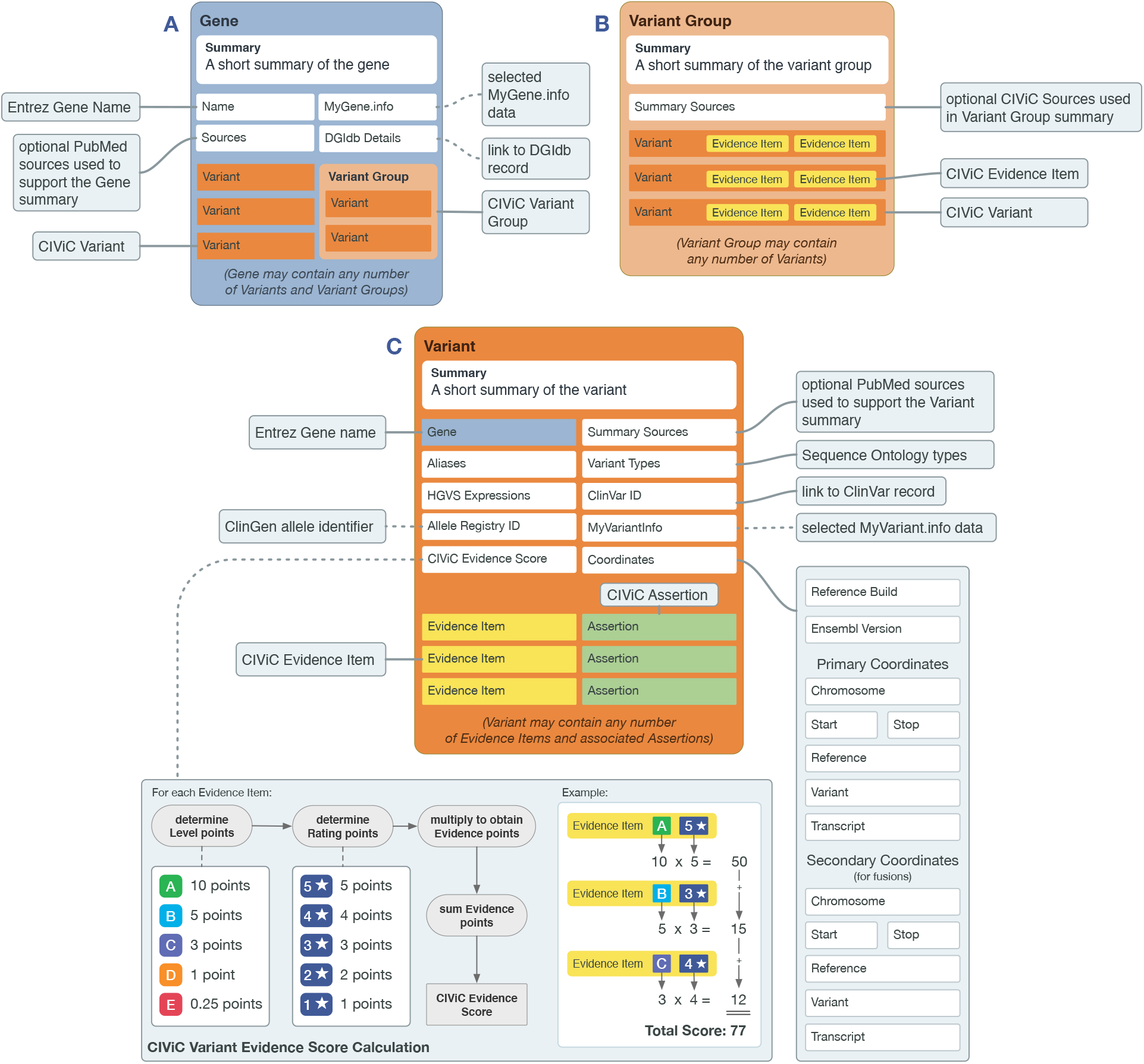
Overview of the Gene and Variant knowledge models and the structure of Variant Groups. The Gene and Variant knowledge models shown below display their associated features (including the Variant Groups feature of Variants) and their origins. Features that are linked to their notes with dotted lines are automatically generated, whenever possible. **A)** Gene data (blue box) consists of curated features (Gene Name, Summary, Sources) and auto-generated links to external entities (MyGene.info and DGIdb). Each Gene can be associated with any number of Variants (dark orange box) and Variants can be grouped (light orange box) based on any unifying feature type (e.g., fusions, activating mutations). **B)** Variant Group features are outlined by the light orange box. These features include a Summary with Sources and associated Variants. **C)** Variant data (dark orange box) includes the Gene Name, Aliases, HGVS Expressions, Variant Evidence Score, Allele Registry ID, Summary Sources, Variant Types, ClinVar IDs, MyVariant.info, and Coordinates. Variants can be associated with CIViC Assertions (green) and Evidence Items (yellow).

#### Curating within the Gene knowledge model

The CIViC **Gene Name** utilizes official Entrez Gene Names from the Entrez Gene database, which are approved by the HUGO Gene Nomenclature Committee (HGNC). Curators must enter a valid Entrez Gene Name (e.g., *TP53*) and should verify the correct entry against the Entrez Gene ID automatically displayed by the CIViC interface. Alternative Gene Names (Aliases/Synonyms) are imported from Entrez and are searchable throughout the database.

A CIViC **Gene Summary** should be created to provide a high-level overview of clinical relevance of cancer variants for the gene. Gene Summaries should focus on emphasizing the clinical relevance from a molecular perspective and should not describe the biological function of the gene unless necessary to contextualize its clinical relevance in cancer. Gene Summaries should include relevant cancer subtypes, specific treatments for the gene’s associated variants, pathway interactions, functional alterations caused by the variants in the gene, and normal/abnormal functions of the gene with associated roles in oncogenesis (**Supplementary Figure 1**). A CIViC Gene Summary should generally be limited to one or two paragraphs and cite relevant reviews to further support the gene’s clinical relevance in cancer.

### Structure and curation of the Variant knowledge model

#### Structure of the Variant knowledge model

A CIViC **Variant** represents any molecular alteration with evidence for clinical relevance in cancer. A new variant is added to the CIViC database when a new Evidence Item for that variant is submitted. The CIViC definition of a variant is intentionally broad to encompass not only simple variation (e.g., SNVs and indels), but also regional variation (e.g., exon mutation), or other types of variation (e.g., expression, fusions, etc.) (**Supplemental Table 2**). Features within the Variant knowledge model include: Variant Summary, Variant Type, HGVS expressions, ClinVar IDs, Variant Evidence Score, representative Variant Coordinates and Transcript, associated Assertions, and external data from MyVariant.info [14] (**Figure 2**). Methods for editing Variant information are shown in **Supplementary Figure 2** and an exemplary Variant entry is shown in **Supplementary Figure 3**.

#### Curating within the Variant knowledge model

The **Variant Name** describes the specific variant being interpreted for clinical utility. When curating this field, the most specific Variant Name described by the source should be used (e.g., *KRAS* G12/G13 rather than *KRAS* Exon 2 Mutation). The Variant Name can be very specific [e.g., *VHL* R176fs (c.528delG)], or can refer to a collection of variants fitting a named category (i.e., *categorical variants [15]*). Examples of categorical variants include *KRAS* G12/G13, *EGFR* Exon 20 Insertion, and *PIK3CA* Mutation (**Supplementary Figure 4**). Other Variant Names, including star-allele nomenclature adopted by pharmacogenetics field (e.g., DPYD*2A; **Supplementary Figure 5**) [16] are also supported. A list of common variant types supported by CIViC are described in **Supplementary Table 2**.

**Variant Aliases** are alternative names, descriptions, or identifiers that differ from the primary CIViC Variant Name. These terms are manually curated and are incorporated into the search fields within the CIViC interface. Aliases may include protein changes on alternative transcripts (e.g., D754Y for *ERBB2* D769Y), dbSNP IDs [17], COSMIC IDs [18] or other identifiers used in the literature.

The **Variant Summary** is a user-defined summary of the clinical relevance of the specific variant. The Variant Summary should be a synthesis of the existing Evidence Statements for the variant. Basic information on recurrence rates and biological/functional impact of the variant may be included, but the focus should be on the clinical impact. Additionally, for Predisposing variants, any appropriate American College of Medical Genetics (ACMG) evidence codes can be recorded (**Supplementary Figure 6**) [19]. Associated sources (PubMed IDs), including valuable review articles that might not be appropriate for the development of Evidence Items, may be used as references for the Variant Summary.

**Variant Type(s)** are used to classify variants by Sequence Ontology terms [18,20]. These terms permit advanced searching for categories of variants in the CIViC interface and downstream semantic analyses of CIViC variants. The most specific term(s) that can be applied to a given variant should be utilized. Use of the Sequence Ontology browser (http://www.sequenceontology.org/browser/obob.cgi) is recommended to identify appropriate terms. When choosing variant types, selection of multiple terms is supported in order to capture both functional and structural effects of the variant (**Supplementary Table 3**). However, these terms should not be ancestors or descendents of one another, and all selected terms should be descendents of the ‘sequence_variant’ term whenever possible.

The **Variant Evidence Score** sums the Evidence Scores for all Evidence Items associated with the Variant. Evidence Item Scores are calculated by multiplying a weighted Trust Rating (i.e., one point for each star) by the values assigned to Evidence Level (i.e., A=10, B=5, C=3, D=1, E=0.5). The Variant Evidence Score is a relative measure of the total amount of curation in the database for a specific variant and does not take into account conflicting evidence.

**Primary and Secondary Coordinates** for each variant are manually curated and verified. Each Variant is assigned representative genomic coordinates (Chromosome, Start, Stop, Reference base, and Variant base) for the assigned reference assembly (e.g., GRCh37). Primary Coordinates are generated for all Variants. Secondary Coordinates are utilized for structural variants involving two loci (e.g., fusion variants). Specific guidelines for choosing representative coordinates and transcripts are described below.

##### Choosing representative coordinates

Although multiple genomic changes can often lead to functionally equivalent alterations (e.g., same amino acid change), CIViC uses representative coordinates to provide user-friendly variant context rather than enumerate all possible alterations that could cause the variant. When choosing a representative variant, it is preferable to use the most common or highly recurrent alteration observed (**Supplementary Figure 7-8**). Genomic coordinates are 1-based with left-shifted normalization and include a specified reference assembly (GRCh37 preferred). Based on manually curated representative coordinates, an automated linkout to the ClinGen **Allele Registry** [21] is created. This link provides additional information such as unique and referenceable identifiers for every registered variant with links to additional resources (e.g., gnomAD, ClinVar). If the required ClinGen Allele does not yet exist, the curator should create a ClinGen account and register it.

##### Choosing a representative transcript

Multiple transcripts can often be expressed for a single gene. For this reason, a specific protein coding alteration, resulting from a genomic change, should always be expressed relative to a specific/individual transcript sequence. CIViC representative transcripts use the Ensembl archived version 75 (GRCh37), and should always include the transcript version number (i.e., ENST00000078429.1 instead of ENST00000078429). There is rarely only one correct transcript. Representative transcripts must contain the variant but are otherwise chosen based on priority criteria such as: wide use in the literature, having the longest open reading frame or most exons, containing the most common exons between transcripts, or having the widest genomic coordinates (**Supplementary Figure 9**). These are consistent with Ensembl’s glossary definition of canonical.

The CIViC Variant knowledge model supports The **Human Genome Variation Society (HGVS) Expression** to describe sequence variation in genomic, RNA, coding DNA, and protein coordinates [22] as well as curated **ClinVar IDs** for each variant. ClinVar IDs and HGVS expressions must be entered individually in the Variant editing interface and may capture ClinVar IDs and HGVS entries not described by the representative coordinates. Manual entry is required (e.g., not automatically linked based on representative coordinates) to permit entries for complex or categorical variants and to support alternate transcripts and reference build versions (**Supplementary Figure 4**).

### Structure and curation of the evidence knowledge model

#### Structure of the Evidence knowledge model

At the core of the CIViC knowledge model lies the CIViC **Evidence Item (EIDs)**. EIDs follow a structured knowledge model with 12 required fields (Gene name, Variant Name, Source Type, Variant Origin, Disease, Evidence Statement, Evidence Type, Evidence Level, Evidence Direction, Clinical Significance, and Trust Rating) with additional optional fields (Associated Phenotypes). Based on the Evidence Type, additional required or optional fields become available (e.g., Predictive Evidence Types require a Drug Name). **Figure 3** describes each field with associated requirements for successful curation and **Supplementary Figure 10-11** show the Evidence Item submission form and display in the Evidence grid.

**Figure 3.**
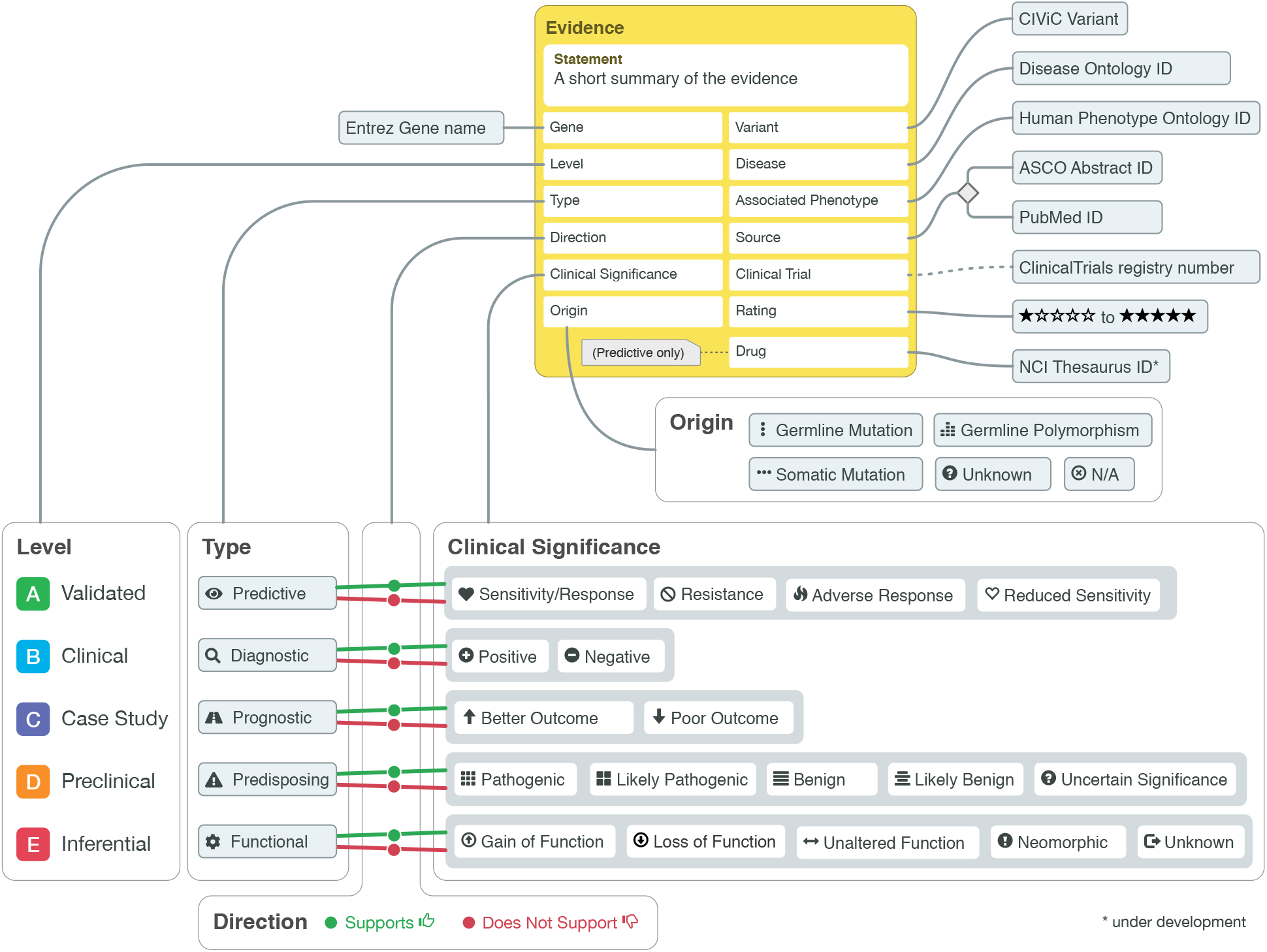
Diagram of the Evidence Item knowledge model. Evidence Items provide a summarized statement about a variant’s implication in clinical oncology in the context of structured data. The knowledge model consists of features (yellow box) that are user-generated and human-readable while leveraging outside ontologies and CIViC-defined fields. Features that are linked to their notes with dotted lines are automatically generated, whenever possible. The Variant Type, Direction, and Clinical Significance features allow curators to develop complex Evidence Item with nuanced meaning while maintaining queryable structure.

#### Curating within the Evidence knowledge model

A **Gene Name** and **Variant Name** are required for EID submission. The Gene Name field will auto-fill using type-ahead search for genes in the Entrez database or their associated Aliases. The Variant Name will also auto-fill based on existing variants within the CIViC database. User-defined variants are also permitted if the desired Variant record does not already exist. To prevent redundancy, it is recommended to browse existing Variant Names for the gene of interest before creating a new term.

Each Evidence Item must be associated with a **Source Type** and **Source ID**, which link the EID to the original publication supporting clinical claims. Currently, CIViC accepts publications indexed on PubMed or abstracts published through the American Society of Clinical Oncology (ASCO). If a PubMed Source Type is selected, the curator can then enter the PubMed ID, which can be verified by comparing the desired source to the abbreviated citation that is automatically generated below the PubMed ID field. If an ASCO Source Type is selected, the ASCO Web ID should be entered into the source ID field. Additionally, **ClinicalTrials Registry Number(s)** are automatically linked via the PubMed database, when available.

The **Variant Origin** categorizes the variant based on method of acquisition. Options for this field include: Somatic Mutation, Germline Mutation, Germline Polymorphism, Unknown, or N/A. The Variant Origin should be somatic if the variant is only found in tumor cells (i.e., arose in a specific, non-germ cell / tissue) and is not expected to be inherited or passed to offspring. The Variant Origin is not applicable (N/A) in some circumstances, particularly in variants that involve differences in expression, methylation, or other post-translational modifications (**Supplementary Figure 12**).

The **Disease** field requires a term that is known to the Disease Ontology (DO) database [23]. The field will auto-fill based on existing diseases (in the cancer subset of DO) and the most specific disease subtype available should be selected. Only a single Disease term can be associated with an EID. If the variant and clinical evidence is implicated in multiple diseases, then multiple Evidence Items should be created. If the Disease cannot be identified in the Disease Ontology, the “Could not find disease” box can be selected and a new field will appear that permits free text entry. In this case, it is recommended to submit a request to the Disease Ontology Term Tracker (http://disease-ontology.org/faq/).

The **Evidence Level** describes the robustness of the study supporting the Evidence Item. Five Evidence Levels are currently available: Validated association (A), Clinical evidence (B), Case study (C), Preclinical evidence (D), Inferential evidence (E) (**Supplementary Figure 13-17**). Validated EIDs (A) have a proven or clinical consensus on the variant association in clinical practice. Typically, these Evidence Items describe Phase III clinical trials, regulatory approvals, or have associated companion diagnostics. Clinical EIDs (B) are typically clinical trials or other primary patient data supporting the clinical association. These EIDs usually include more than 5 patients supporting the claim made in the Evidence Statement. Case studies (C) are individual case reports or small case series. Preclinical evidence (D) is derived from *in vivo* or *in vitro* experiments (e.g., mouse models or cell lines) that support clinical claims. Finally, inferential associations (E) indirectly associate the variant to the provided clinical evidence. These can involve hypotheses generated from previous experiments but not yet supported by experimental results.

The **Evidence Type** refers to the type of clinical (or biological) association described by the Evidence Item’s clinical summary. Five Evidence Types are currently supported: Predictive (i.e. Therapeutic), Diagnostic, Prognostic, Predisposing, and Functional. Each Evidence Type describes the clinical or biological effect a variant has on the following: therapeutic response (Predictive), determining a patient’s diagnosis or disease subtype (Diagnostic), predicting disease progression or patient survival (Prognostic), disease susceptibility (Predisposing), or biological alterations relevant to a cancer phenotype (Functional) (**Supplemental Figures 18-22**). Selecting an Evidence Type has implications on available selections for Clinical Significance, as outlined in **Figure 2** and **Supplementary Table 5**.

The **Evidence Direction** indicates if the Evidence Statement supports or refutes the clinical significance of an event. The available options include: “Supports” or “Does not support”. Nuanced examples for how to correctly use the Evidence Direction for Predictive Evidence Types are shown in **Supplementary Table 4** and **Supplementary Figure 23**.

**Clinical Significance** describes how the variant is related to a specific, clinically-relevant property as described in the Evidence Statement. The available options for Clinical Significance depend on the Evidence Type selected for the Evidence Statement. These options are shown in **Figure 3** with details in **Supplementary Table 5**. In brief, they describe the severity or type of treatment response (Predictive), inclusivity or exclusivity of a cancer type or subtype (Diagnostic), the type of outcome (Prognostic), the germline variant classification according to ACMG guidelines [19] (Predisposing), or the type of biological change (Functional). Note that predisposing Evidence Items may include ACMG evidence codes in the Evidence Statement; however, very few publications single-handedly warrant classification beyond a Variant of Unknown Significance.

The **Trust Rating** is scored on a scale from 1-5 stars reflecting the curator’s confidence in the quality of the summarized evidence (**Supplementary Figure 24-28**). This rating depends on a number of factors, including journal impact, study size, study design, orthogonal validation, and reproducibility. Although the overall publication/study might be high quality and in a high impact publication, the Trust Rating may be low for an Evidence Item referring to a single conclusion in the study that is not well supported. This is a largely subjective measure, however, general guidelines for trust ratings are provided in **Supplementary Table 6**.

The **Evidence Statement** is a brief summary of the clinical implications of the Variant in the context of a specific Disease, Evidence Type and Clinical Significance as described in the cited literature source. An Evidence Statement should synthesize the information from a published study relevant to the clinical association of the variant. Evidence Statements should be as brief as possible (typically 1 to 3 sentences), but include sufficient experimental detail to interpret and evaluate the evidence without repeating the original text or using domain-specific acronyms or colloquialisms. Such details include the type of study (e.g., phase, design), controls used, outcomes measured, the number of individuals involved and relevant statistical values (e.g., p-values, R^2^, confidence intervals). Data constituting protected health information (PHI) should not be entered in the Evidence Statement field.

For Predictive evidence items, a **Drug Names** field will become available. Multiple drugs can be added to a single Evidence Item, requiring a **Drug Interaction Type** (Combination, Sequential, or Substitutes) that describes the relationship of these drugs in the study. The Drugs and Drug Interaction Types should be explicitly stated in the source supporting the Evidence Item and not inferred by the curator. Trade names should not be used for Drugs. Older drug names/aliases should be referred to by their newer name in the Drug field while mentioning the old and new name in the Evidence Statement to minimize confusion (**Supplementary Figure 16**).

When additional phenotypes not captured by the Disease field alone are indicated, **Associated Phenotypes** available in the Human Phenotype Ontology (HPO) database [24] can be added to any Evidence Item. Associated Phenotypes should provide additional information beyond what is implied by the Disease field. Phenotypes should be particularly considered for Predisposing Evidence Items whereby the variant is associated with a non-binary phenotype or syndrome for a particular genotype.

The last field in the Evidence Item submission form permits free-form text for additional comments about the Evidence Item. For example, curators can call an editor’s attention to a particular comment using macro notation (**Supplementary Table 7**). These comments will appear first in the item’s comment thread and will be visible to editors during review.

### Structure and curation of the assertion knowledge model

#### Structure of the Assertion knowledge model

The CIViC **Assertion** summarizes the clinical relevance of a variant in a specific disease context using a collection of Evidence Items (**Figure 4**). Consistent with Evidence Items, Assertions include a Gene, Variant, Variant Origin, Disease, Assertion Type, Assertion Direction, Clinical Significance, Drug (Predictive), Drug Interaction Type (Predictive), and Associated Phenotypes (optional). Fields unique to Assertions include annotation with clinical guidelines such as Association for Molecular Pathology (AMP) Tier and Level, ACMG codes, National Comprehensive Cancer Network (NCCN) guideline/version, FDA approvals/diagnostics. A short, one sentence Summary and a longer Description of the Assertion are also required for submission. If available, existing Evidence Items should be associated with the Assertion to support the Summary/Description. The Assertion curation form can be found in **Supplementary Figure 29**.

**Figure 4.**
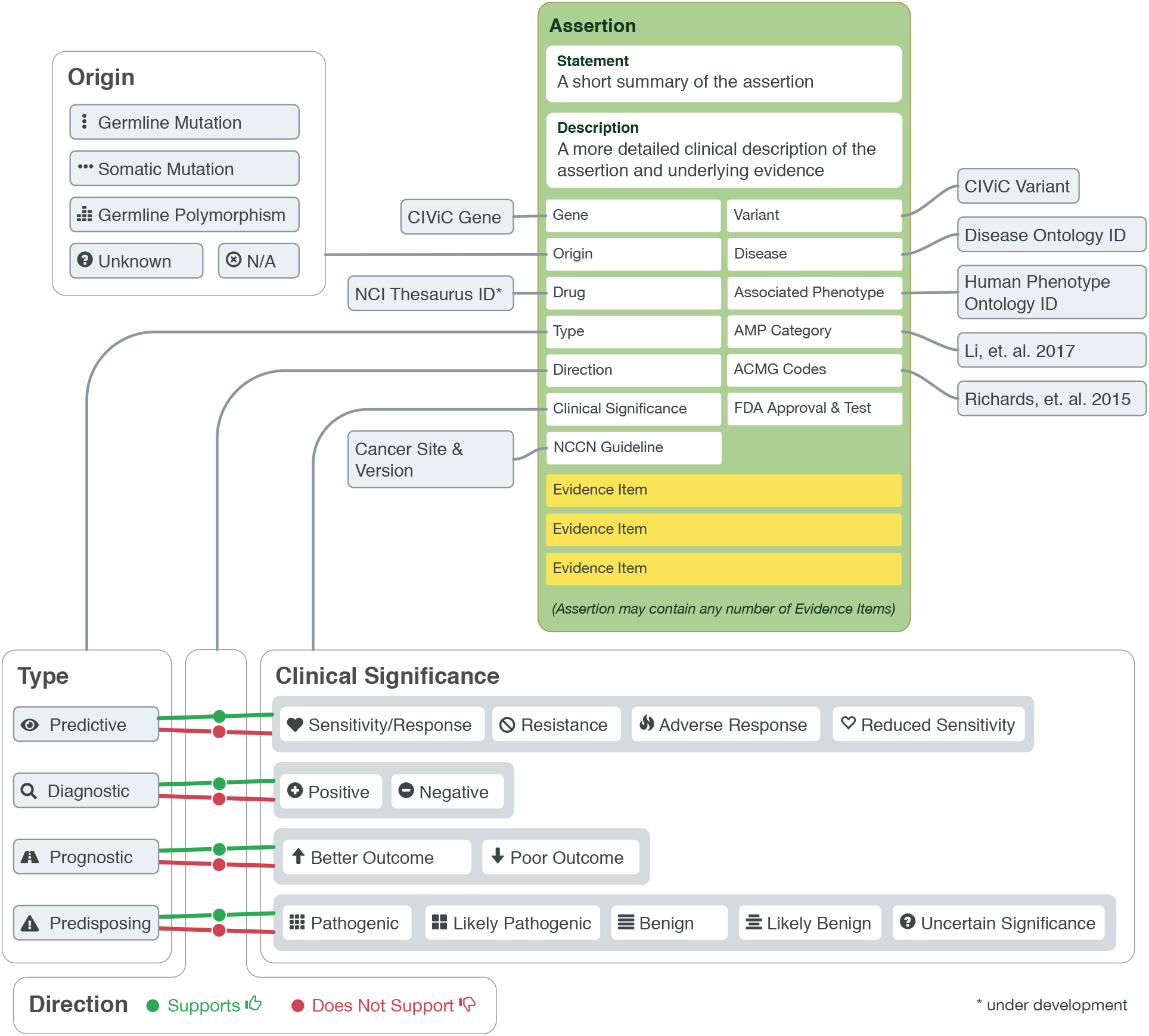
Diagram of knowledge model for CIViC Assertions. Assertions summarize a collection of Evidence Items to make a definitive clinical statement about the Variant in a specific Disease context which incorporates all known data within the knowledgebase. Assertions features (green box) build on the Evidence Item knowledge model to bring together clinical guidelines, public resources, and regulatory approvals relevant to a final variant interpretation. Assertions can be associated with any number of Evidence Items. Like Evidence Items, Assertion Type, Direction, and Clinical Significance can be used to create a specific meaning for the Assertion.

#### Curating within the Assertion knowledge model

The **Gene Name** and **Variant Name** for an Assertion have curation constraints. Assertions can only be created for Genes and Variants associated with at least one Evidence Item and are selected from an auto-populated list using type-ahead search. Variant Names are restricted to those associated with the selected Gene. The **Variant Origin** follows the same guidelines as described for Evidence Items (**Supplementary Figure 30**).

The **Disease** associated with the Assertion must already exist within the CIViC database. Only one Disease is permitted for each Assertion. It is recommended that the Disease be as specific as possible while still holding true for all Evidence Items associated with the Assertion (e.g., an Assertion for “non-small cell lung cancer” can be supported by Evidence Items associated with “lung adenocarcinoma” and “non-small cell lung cancer” as well as general disease categories such as “cancer”) (**Supplementary Figure 31)**.

CIViC currently supports the following **Assertion Types**: Predictive, Diagnostic, Prognostic, and Predisposing. As with the Evidence Item submission form, selecting an Assertion Type will alter available choices for **Clinical Significance**, which is outlined in **Figure 4**. Options for **Assertion Direction** include “Supports” and “Does not Support”. Predictive, Prognostic or Diagnostic Assertions (**Figure 5A, Supplementary Figure 32-34**), utilize the somatic variant interpretation guidelines, providing an AMP Tier (I-IV) and Level (A-D) [25]. Predisposing Assertions (**Figure 5B, Supplementary Figure 35**) utilize the ACMG guideline classifications (Pathogenic, Likely Pathogenic, Likely Benign, Benign and Variant of Unknown Significance), their predicate ACMG evidence codes (i.e., PVS1, PP2, etc), and rules for combining criteria [19], as well as recommended updates [26–28]. Assertions are classified based on the combination of evidence [EIDs and public sources (e.g., gnomAD, CADD)], associated with the Assertion. The Assertion Description should specify the guidelines or classification system used.

**Figure 5.**
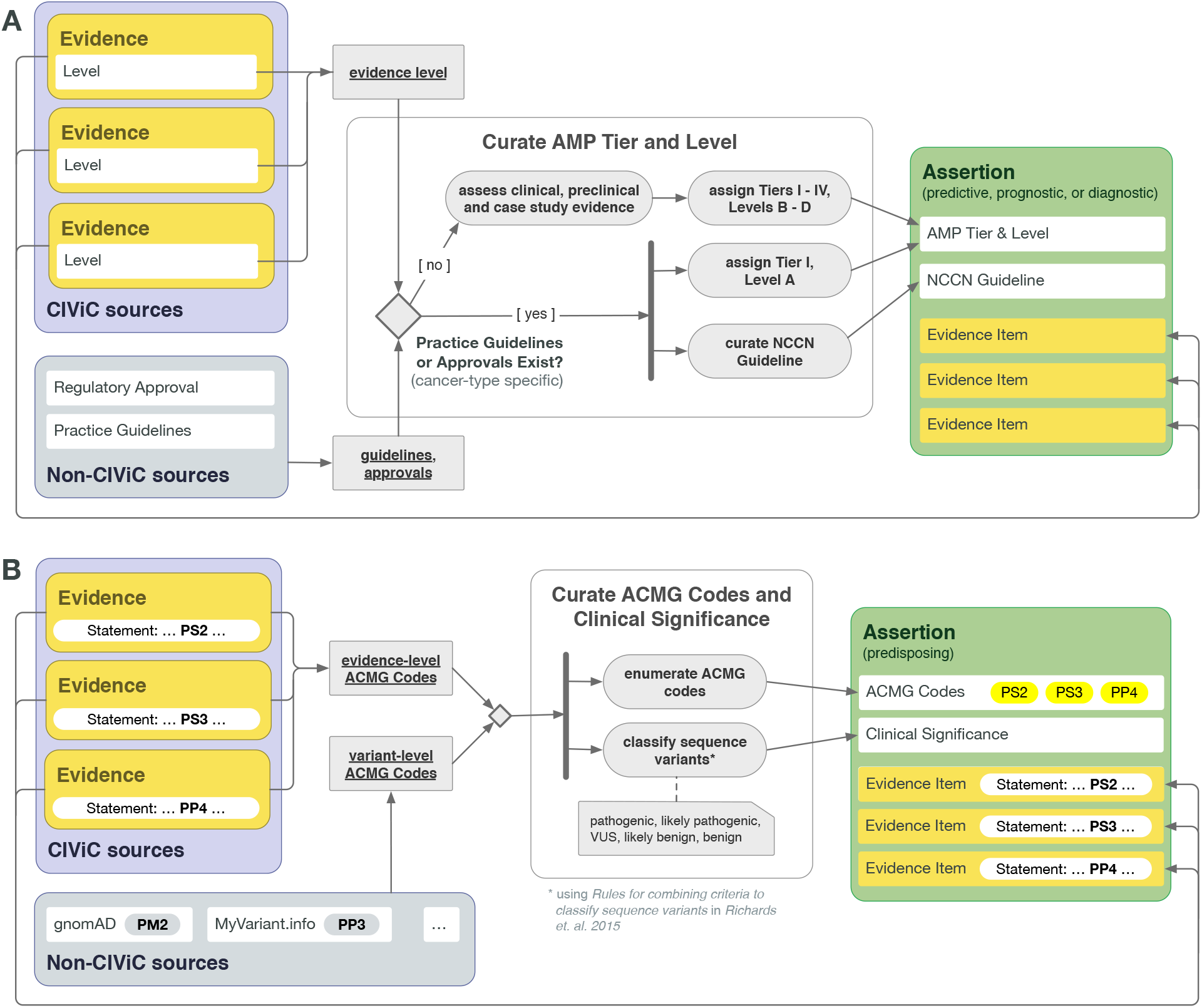
CIViC Assertion development by Assertion Type. CIViC Assertions summarize a collection of Evidence Items which reflect the state of literature for the given variant and disease. **A)** For Assertion Types typically associated with somatic variants (Predictive, Prognostic, or Diagnostic), AMP guidelines are followed to associate the Assertion with an AMP Tier and Level, which involves consideration of practice guidelines as well as regulatory approvals associated with specific drugs, as well as consideration of available clinical evidence in the absence of explicit regulatory or practice guidelines. **B)** CIViC Predisposing Assertions utilize ACMG guidelines to generate a 5-tier pathogenicity valuation for a variant in a given disease context, which is supported by a collection of CIViC Evidence Items, along with other data. ACMG evidence codes for an Assertion are supplied by a collection of supporting CIViC Evidence Items (e.g., PP1 from co-segregation data available in a specific publication), and additionally are derived from Variant data (e.g., PM2 from population databases such as gnomAD). ACMG evidence codes are then combined at the Assertion level to generate a disease-specific pathogenicity classification for the Assertion.

Optional descriptive fields for Assertions include **Associated Phenotypes** and **NCCN Guideline(s)/Version(s)** (**Supplementary Figure 29**). If the Variant-Disease association described by the Assertion has a cleared/approved FDA companion diagnostic or a drug with **FDA Regulatory Approval**, then the appropriate box should be checked.

Each Assertion requires a one-sentence **Summary** and a longer, more complete **Description** of the Assertion. The Description is designed to capture special considerations or additional data (e.g., specific treatment regimens, source of ACMG codes) used by the curator to assemble the Assertion. Important specific details from practice guidelines (e.g. NCCN) should be included in the Summary, including disease stage, and in the case of predictive Assertions, treatment line which practice guidelines recommend.

The **Supporting Evidence** grid allows users to associate Evidence Items with Assertions. This collection of Evidence Items should cover the important clinically relevant findings for the variant in the context of a specific cancer. For Predictive assertions, the collection of Evidence Items should also consider the Drug(s) and their Drug Interaction Type. Assertions do not require Evidence Items for development; however, complete (revised and accepted) Evidence Items must be added to the Assertion before it can be accepted by editors (**Supplementary Figure 36**). While there is no enforced minimum Evidence Item count, a sufficient amount of Evidence should be added to Assertions to independently support the chosen ACMG classification or AMP Tier/Level.

## Conclusions

The CIViC knowledge model has been developed to permit all individuals within the oncology field to effectively incorporate variant interpretations into a structured knowledgebase. The presented framework supports a wide variety of variants with complex clinical annotations (**Supplementary Table 2**) and permits inclusion of data from multiple variant interpretation sources. Additionally, the presented knowledge models provide a framework that allows users to both explore and curate clinical interpretations of variants in cancer. The curation standard operating procedures presented here can be utilized to improve clarity and ease for our community of curators. This will hopefully increase the overall quality of data entered into the knowledgebase.

It is our hope that future advancements and updates to the presented framework will further expand CIViC’s ability to capture all variants and associated clinical annotations. Advancements include development of supplemental features that increase the breadth of variants and annotations that are supported by the interface. Specifically, we hope to support annotations for variant combinations and large-scale or genomewide alterations (e.g., mismatch repair deficiency). We also plan to continuously update the evolving variant annotation standards/guidelines and integrate curation from additional external knowledgebases [29].

The field is currently facing exponential growth and complexity of variant interpretation, which underscores the need for a standardized and comprehensive schematic for variant curation and interpretation. The presented CIViC knowledge models provide such a framework for optimized curation of variants. The use of integrated and hierarchical knowledge models with standardized methods for curation creates the ability to generate a powerful knowledgebase that can improve clinical annotation of variants in cancer.

## Supporting information

Supplementary Figures and Lables

## List of Abbreviations

API: Application Programming Interface
ACMG: American College of Medical Genetics
AID: CIViC Assertion identifier
AMP: Association for Molecular Pathology
ASCO: American Society of Clinical Oncology
CIViC: Clinical Interpretations of Variants In Cancer
DGIdb: Drug Gene Interaction database
DOID: Disease Ontology identifiers
EID: CIViC Evidence Item identifier
FDA: Food and Drug Administration
HGNC: Human Gene Nomenclature Committee
HPO: Human Phenotype Ontology
NCI: National Cancer Institute
NCCN: National Comprehensive Cancer Network
SOP: Standard Operating Procedure
SOID: Sequence Ontology identifier

## Declarations

### Ethics approval and consent to participate

Not applicable

### Consent for publication

Not applicable

### Availability of data and material

Not applicable

### Authors’ contributions

AMD, KK, EKB, ACC, JFM, SK, NCS, AHW, MG, OLG developed the knowledgebase and knowledge models. AMD, KK, EKB, NCS, LMS, SPP, LK, KAC, AW, SR, DIR, DS, GR, RHK, AHW, SM, MG, OLG developed curation practices. ACC, JFM, and SK developed and implemented the CIViC application and curation interface. AMD, KK, DS, GR, RHK, SM, AHW contributed to the integration of CIViC with external guidelines, standards and ontologies. AMD, KK, EKB wrote the manuscript. AMD, KK, EKB, ACC, JFM, SK, AHW, MG, OLG revised and edited the manuscript. JFM designed the figures with conceptual input from AMD, KK, EKB, AHW. MG and OG supervised the study. All authors reviewed and approved the manuscript.

## Acknowledgements

We are grateful to the community of curators, editors, domain experts and users who make CIViC possible. We would also like to thank cancer patients and their families who consent to participate in research studies. Without their generous contributions, much of the knowledge represented in CIViC would not exist.

## References

1. Kamps R, Brandão RD, van den Bosch BJ, Paulussen ADC, Xanthoulea S, Blok MJ, et al. Next-Generation Sequencing in Oncology: Genetic Diagnosis, Risk Prediction and Cancer Classification. Int J Mol Sci [Internet]. 2017;18. Available from: http://dx.doi.org/10.3390/ijms18020308

2. Good BM, Ainscough BJ, McMichael JF, Su AI, Griffith OL. Organizing knowledge to enable personalization of medicine in cancer. Genome Biol. 2014;15:438.

3. The AACR Project GENIE Consortium. AACR Project GENIE: Powering Precision Medicine through an International Consortium. Cancer Discov. American Association for Cancer Research; 2017;7:818–31.

4. Consortium TICG, The International Cancer Genome Consortium. Erratum: International network of cancer genome projects. Nature. 2010;465:966–966.

5. Cancer Genome Atlas Research Network, Weinstein JN, Collisson EA, Mills GB, Shaw KRM, Ozenberger BA, et al. The Cancer Genome Atlas Pan-Cancer analysis project. Nat Genet. 2013;45:1113–20.

6. Hoskinson DC, Dubuc AM, Mason-Suares H. The current state of clinical interpretation of sequence variants. Curr Opin Genet Dev. 2017;42:33–9.

7. Yorczyk A, Robinson LS, Ross TS. Use of panel tests in place of single gene tests in the cancer genetics clinic. Clin Genet. 2015;88:278–82.

8. Amendola LM, Dorschner MO, Robertson PD, Salama JS, Hart R, Shirts BH, et al. Actionable exomic incidental findings in 6503 participants: challenges of variant classification. Genome Res. 2015;25:305–15.

9. Shah PD, Nathanson KL. Application of Panel-Based Tests for Inherited Risk of Cancer. Annu Rev Genomics Hum Genet. 2017;18:201–27.

10. Griffith M, Spies NC, Krysiak K, McMichael JF, Coffman AC, Danos AM, et al. CIViC is a community knowledgebase for expert crowdsourcing the clinical interpretation of variants in cancer. Nat Genet. 2017;49:170–4.

11. Griffith M, Griffith OL, Coffman AC, Weible JV, McMichael JF, Spies NC, et al. DGIdb: mining the druggable genome. Nat Methods. 2013;10:1209–10.

12. Wagner AH, Coffman AC, Ainscough BJ, Spies NC, Skidmore ZL, Campbell KM, et al. DGIdb 2.0: mining clinically relevant drug–gene interactions. Nucleic Acids Res. Narnia; 2016;44:D1036–44.

13. Cotto KC, Wagner AH, Feng Y-Y, Kiwala S, Coffman AC, Spies G, et al. DGIdb 3.0: a redesign and expansion of the drug–gene interaction database. Nucleic Acids Res. Narnia; 2018;46:D1068–73.

14. Xin J, Mark A, Afrasiabi C, Tsueng G, Juchler M, Gopal N, et al. High-performance web services for querying gene and variant annotation. Genome Biol. 2016;17:91.

15. Patterson SE, Statz CM, Yin T, Mockus SM. Utility of the JAX Clinical Knowledgebase in capture and assessment of complex genomic cancer data. NPJ Precis Oncol. 2019;3:2.

16. Robarge JD, Li L, Desta Z, Nguyen A, Flockhart DA. The star-allele nomenclature: retooling for translational genomics. Clin Pharmacol Ther. 2007;82:244–8.

17. Kitts A, Phan L, Ward M, Holmes JB. The Database of Short Genetic Variation (dbSNP). National Center for Biotechnology Information (US); 2014.

18. Tate JG, Bamford S, Jubb HC, Sondka Z, Beare DM, Bindal N, et al. COSMIC: the Catalogue Of Somatic Mutations In Cancer. Nucleic Acids Res. Narnia; 2019;47:D941–7.

19. Richards S, Aziz N, Bale S, Bick D, Das S, Gastier-Foster J, et al. Standards and guidelines for the interpretation of sequence variants: a joint consensus recommendation of the American College of Medical Genetics and Genomics and the Association for Molecular Pathology. Genet Med. 2015;17:405–24.

20. Eilbeck K, Lewis SE, Mungall CJ, Yandell M, Stein L, Durbin R, et al. The Sequence Ontology: a tool for the unification of genome annotations. Genome Biol. 2005;6:R44.

21. Pawliczek P, Patel RY, Ashmore LR, Jackson AR, Bizon C, Nelson T, et al. ClinGen Allele Registry links information about genetic variants. Hum Mutat. 2018;39:1690–701.

22. HGVS Expressions at NCBI [Internet]. [cited 2019 May 17]. Available from: https://www.ncbi.nlm.nih.gov/variation/hgvs/

23. Kibbe WA, Arze C, Felix V, Mitraka E, Bolton E, Fu G, et al. Disease Ontology 2015 update: an expanded and updated database of human diseases for linking biomedical knowledge through disease data [Internet]. Nucleic Acids Research. 2015. p. D1071–8. Available from: http://dx.doi.org/10.1093/nar/gku1011

24. Köhler S, Doelken SC, Mungall CJ, Bauer S, Firth HV, Bailleul-Forestier I, et al. The Human Phenotype Ontology project: linking molecular biology and disease through phenotype data. Nucleic Acids Res. 2014;42:D966–74.

25. Li MM, Datto M, Duncavage EJ, Kulkarni S, Lindeman NI, Roy S, et al. Standards and Guidelines for the Interpretation and Reporting of Sequence Variants in Cancer: A Joint Consensus Recommendation of the Association for Molecular Pathology, American Society of Clinical Oncology, and College of American Pathologists. J Mol Diagn. 2017;19:4–23.

26. Mester JL, Ghosh R, Pesaran T, Huether R, Karam R, Hruska KS, et al. Gene-specific criteria for PTEN variant curation: Recommendations from the ClinGen PTEN Expert Panel. Hum Mutat. Wiley Online Library; 2018;39:1581–92.

27. Ghosh R, Harrison SM, Rehm HL, Plon SE, Biesecker LG, ClinGen Sequence Variant Interpretation Working Group. Updated recommendation for the benign stand-alone ACMG/AMP criterion. Hum Mutat. 2018;39:1525–30.

28. Abou Tayoun AN, Pesaran T, DiStefano MT, Oza A, Rehm HL, Biesecker LG, et al. Recommendations for interpreting the loss of function PVS1 ACMG/AMP variant criterion. Hum Mutat. 2018;39:1517–24.

29. Wagner AH, Walsh B, Mayfield G, Tamborero D, Sonkin D, Krysiak K, et al. A harmonized meta-knowledgebase of clinical interpretations of cancer genomic variants [Internet]. bioRxiv. 2018 [cited 2019 May 31]. p. 366856. Available from: https://www.biorxiv.org/content/10.1101/366856v2

