## Supplementary Figures and Lables for "The CIViC knowledge model and standard operating procedures for curation and clinical interpretation of variants in cancer"

### Supplementary Materials

#### Table of Contents


Curating Gene-level entities

- Supplementary Figure 1. Visualizing and curating the Gene-level knowledge model.

Curating Variant-level entities

- Supplementary Figure 2. Visualizing curated variants within the Variant-level knowledge model
- Supplementary Figure 3. Exemplary Variant that has been curated in CIViC.
- Supplementary Figure 4. Example of a categorical variant.
- Supplementary Figure 5. Example of a CIViC variant that has clinical implications in pharmacogenetics.
- Supplementary Figure 6. Example of a CIViC variant summary with inline inclusion of ACMG evidence codes.
- Supplementary Figure 7. Defining variant coordinates for SNVs and small indels
- Supplementary Figure 8. Defining variant coordinates for categorical and large-scale variants
- Supplementary Figure 9. Choosing a Representative Transcript

The Evidence Item data model

- Supplementary Figure 10. Overview of CIViC evidence entry form
- Supplementary Figure 10.5 View of the Evidence Grid for a given Variant

Examples of Variant Origin

- Supplementary Figure 11. Example of Evidence Item where the Variant Origin is not applicable (N/A).

Examples of Evidence Levels

- Supplementary Figure 12. Example of a A-level (Validated) Evidence Item.
- Supplementary Figure 13. Example of a B-level (Clinical) Evidence Item.
- Supplementary Figure 14. Example of a C-level (Case Study) Evidence Item.
- Supplementary Figure 15. Example of a D-level (Preclinical) Evidence Item.
- Supplementary Figure 16. Example of an E-level (Inferential) Evidence Item.

Examples of Evidence Types

- Supplementary Figure 17. Example of a Predictive Evidence Type
- Supplementary Figure 18. Example of a Diagnostic Evidence Type
- Supplementary Figure 19. Example of a Prognostic Evidence Type
- Supplementary Figure 20. Example of a Predisposing Evidence Type
- Supplementary Figure 21. Example of a Functional Evidence Type
- Supplementary Figure 22. Interpreting clinical trial data to curate Predictive and Prognostic Evidence Items

Examples of Trust Ratings

- Supplementary Figure 23. Evidence Item with 5-star Trust Rating
- Supplementary Figure 24. Evidence Item with 4-star Trust Rating
- Supplementary Figure 25. Evidence Item with 3-star Trust Rating
- Supplementary Figure 26. Evidence Item with 2-star Trust Rating
- Supplementary Figure 27. Evidence Item with 1-star Trust Rating

The Assertion data model
Examples of Assertion curation
Supplementary Figure 28. Overview of CIViC assertion entry form.
Examples of Variant Origin
Supplementary Figure 29. Example of curation of Variant Origin.
Examples of Disease
Supplementary Figure 30. Selection of Disease Type for Assertions
Examples of Assertions
Supplementary Figure 31. Exemplary Predictive Assertions
Supplementary Figure 32. Exemplary Prognostic Assertions
Supplementary Figure 33. Diagnostic Assertions
Supplementary Figure 34. Example of a Predisposing Assertion
Examples of Assertion Supporting Evidence
Supplementary Figure 35. Requirements for an Assertion to be accepted.

Supplementary Tables
Supplementary Table 1. General curation notation for all items within CIViC
Supplementary Table 2. Examples of Variants supported by the CIViC interface
Supplementary Table 3. Examples of classification Sequence Ontology Term classifications.
Supplementary Table 4. Use cases for all combinations of Evidence Types, Direction, and Clinical Significance for Predictive Evidence.
Supplementary Table 5. Definitions of Clinical Significance for all Evidence Types.
Supplementary Table 6. General guidelines and examples for trust ratings.
Supplementary Table 7. Markdown and Macros.

References

### Curating Gene-level entities

#### Supplementary Figure 1. Visualizing and curating the Gene knowledge model

This example shows the Gene record for *BRAF* (top). The Gene knowledge model displays the Gene summaries with associated sources, a link to a DGIdb [1–3] search for the associated gene, and information pulled from MyGene.info [4] (blue box) with a link to additional details and resources. Selection of the purple pencil in the upper left corner activates the Suggested Revision form (bottom). This form allows curators to edit the Gene Summary and associated sources and also requires a Revision Description.

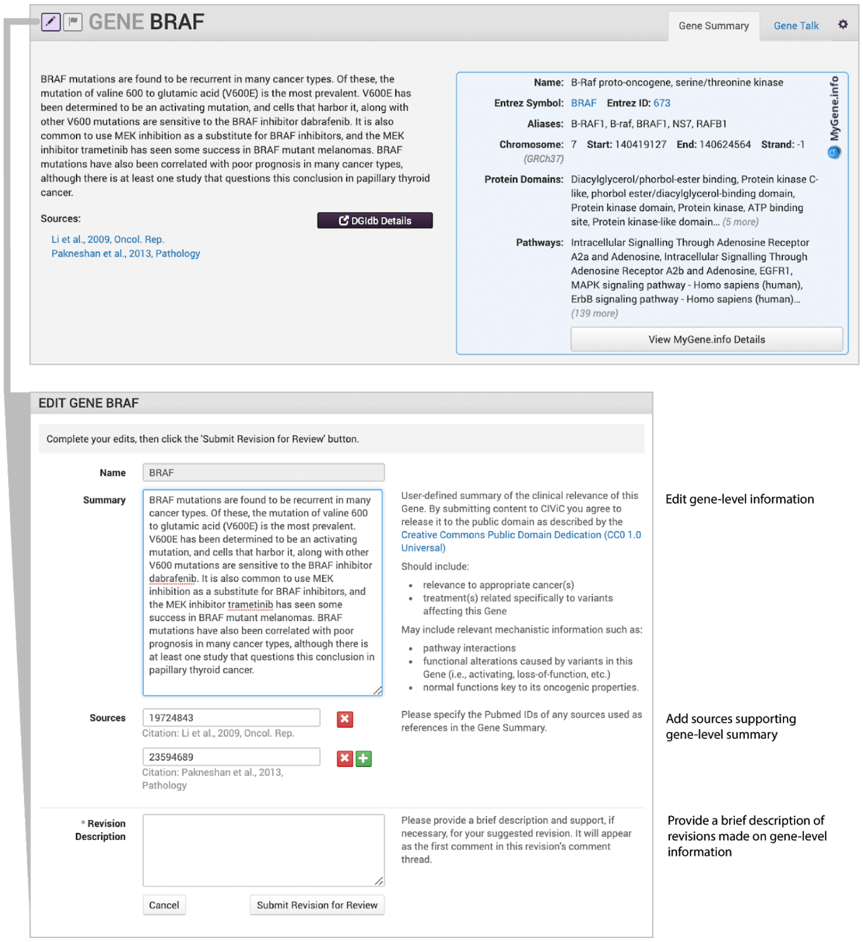


**Curation Practices:**

- Gene Summaries should include relevant cancer subtypes, specific treatments for the gene's associated variants, pathway interactions, functional alterations caused by the variants in the gene, and normal/abnormal functions of the gene with associated roles in oncogenesis.
- A CIViC Gene Summary should generally be limited to one or two paragraphs and cite relevant reviews for a more extensive discussion of clinical relevance of the gene in cancer.
- The sources used for Gene Summaries should be derived from Pubmed and, unlike typical CIViC Evidence Items, may include review articles.
- Instructions for curation are provided in the right column of the Suggested Revision form.

### Curating Variant-level entities

**Supplementary Figure 2. Visualizing and curating the Variant knowledge model**

Variants within a gene are displayed in a dynamic list (top left panel) that can be quickly filtered by name. Variant Groups provide user-defined grouping of Variants within and between genes that can be quickly created. The Variant knowledge model (bottom left panel) includes the Variant Name, Aliases, Variant Summary and associated Sources, HGVS Expressions, ClinVar IDs, Variant Types, and Primary Coordinates. Secondary Coordinates (not shown) can also be added to describe genomic context for fusion variants. Selecting any Variant from the list will show the Variant data and associated Evidence Items. Selecting the pencil in the top left corner allows individuals to enter or edit Variant data using a Suggested Revision form (right panel).

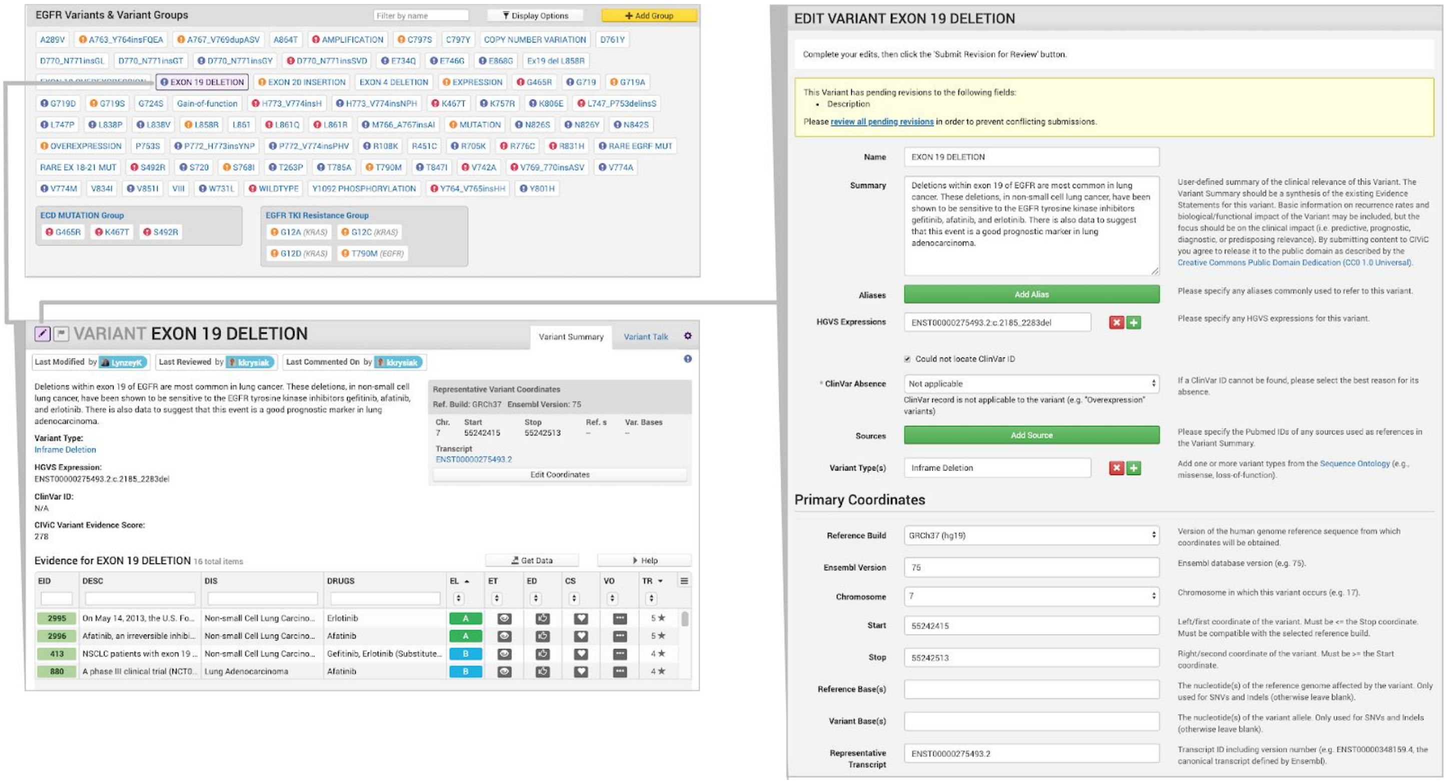


**Curation Practices:**

- Display options allow users to tailor their variant viewing experience by displaying some or all Variants by their status (by default, only variants with at least one accepted Evidence Item are shown).
- Status icons containing the exclamation point indicate variants with active curation involving submitted evidence (orange), pending revisions (purple), or both (red).
- The Evidence Grid for each Variant provides quick filtering and sorting options with status filtering provided by the 3 line icon in the top right.
- Users can download the Evidence Grid by clicking “Get Data”.
- When adding ASCO Abstract Sources, it should be noted that the ASCO Web ID is not the ASCO abstract number. The ASCO Web ID is the identification number found in the URL of the abstract.

**Supplementary Figure 3. Exemplary Variant that has been curated in CIViC**

This example shows the BCR-ABL T315I variant, which can be used to guide treatment decisions for patients with chronic myeloid leukemia. The Variant Name describes the specific protein change (T315I) that can be observed in conjunction with the pathognomonic fusion (BCR-ABL) for CML. The Variant Summary describes how this variant combination confers resistance to imatinib but sensitivity/response to newer tyrosine kinase inhibitors such as dasatinib and ponatinib. Variant Aliases, HGVS Expression, Variant Types, and ClinVar IDs have all been manually curated with links to external databases, if applicable. The Representative Variant Coordinates describe the most common single base pair change that could result in a T315I variant and these coordinates were used to link to MyVariant.info. The CIViC Variant record currently has 40 submitted Evidence Items that support the listed Summary of the Variant, many of which are B-level (Clinical) Evidence Items. The status of an Evidence item is indicated by color with submitted in orange or accepted in green.

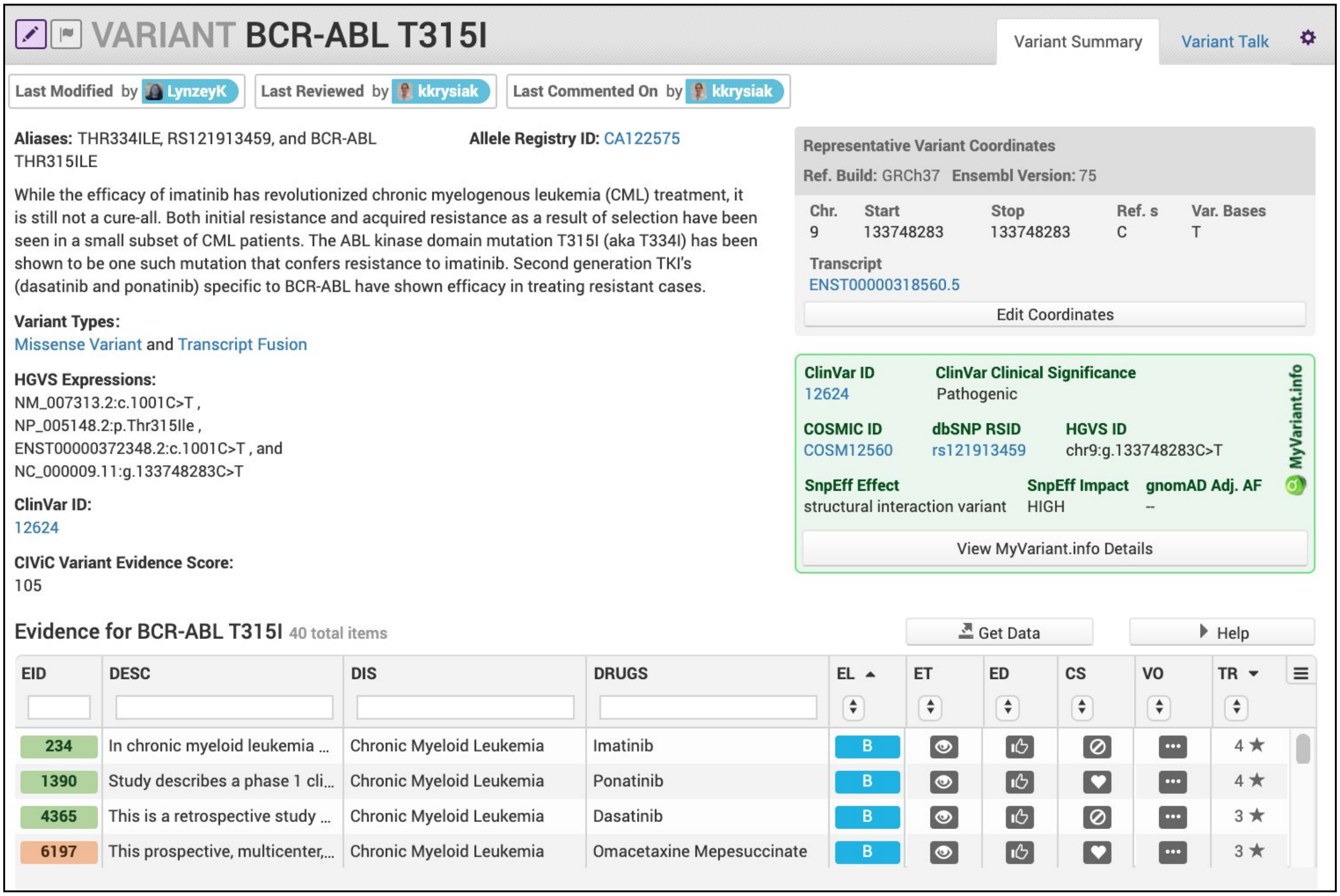


**Curation Practices:**

- The Variant Summary should attempt to describe all associated and accepted Evidence Items for that Variant. If possible, be specific about the Diseases being referenced or the implicated Drugs.
- Variants can be associated with multiple Variant Types; however, the most specific term should be used and multiple terms related to one another as ancestors or descendants should not be used.
- Note that moderated and accepted Evidence Items (EIDs) are labeled green (EIDs 234, 1390, and 4365) while EIDs that have not yet completed moderation are labeled orange (EID 6197).

**Supplementary Figure 4. Example of a categorical variant**

The image below depicts the KRAS - G12/13 categorical variant. The term *categorical variant* (sometimes called *bucket variant*) is given to any collection of variants that include some level of ambiguity that prevents assignment of an Evidence Item to a specific variant. Due to factors such as the assay used, sample sizes, or experimental design, such ambiguous groups of variants are prevalent throughout the literature. For example, studies often look at survival differences between patients with or without a mutation in a given gene to show a significant difference in clinical outcome between populations. However, the clinical outcome is either not significant or not enumerated for the specific variants observed for each individual within the study. Categorical variants are associated with multiple ClinVar entries, if applicable. In the example below, each of the ClinVar IDs links to a specific missense variant that can change *KRAS* at amino acid positions G12 and/or G13.

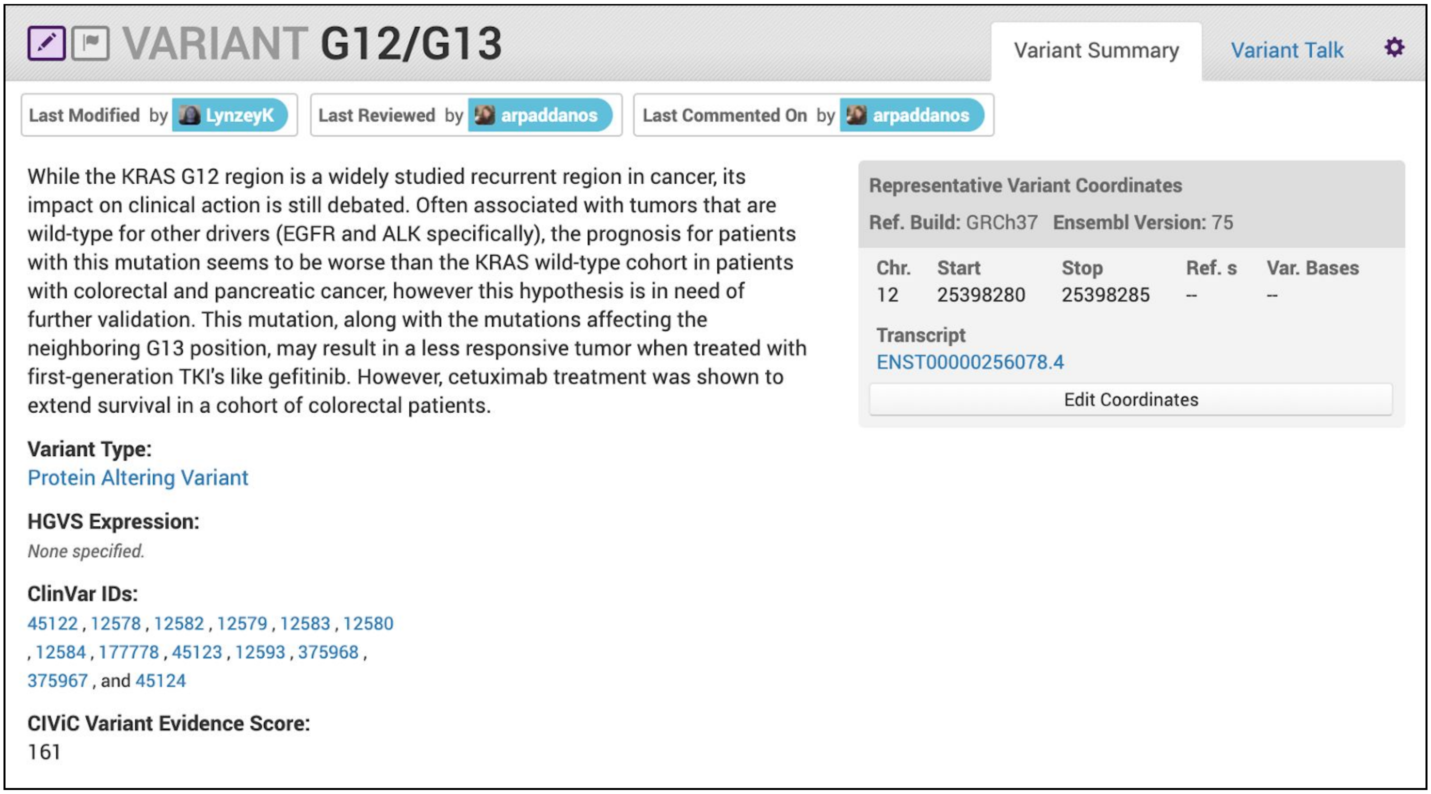


#### Curation Practices

- Curators can develop categorical variants for a variety of reasons. For example, categorical variants can be used to enhance statistical power of the variant being analyzed. Alternatively, the categorical Variant Name may reflect the assay used (e.g., Sanger sequencing of a small region).
- Often, data used to generate Evidence Items that are associated with categorical variants can also be used to develop additional Evidence Items for individual case studies. For example, a group of individuals with a single categorical mutation (e.g., *TP53* - Mutation) can be used to generate a B-level Evidence Item that supports response to a therapeutic. Additionally, the specific variants from this cohort (e.g., *TP53* - R282L and *TP53* - R213P) can be used to generate C-level Evidence Items to describe therapeutic response for individual cases.

**Supplementary Figure 5. Example of a CIViC variant that has clinical implications in pharmacogenetics**

The image below depicts the DPYD*2A Variant, which is a germline polymorphism that has pharmacogenetic implications. Specifically, the Clinical Pharmacogenomics Implementation Consortium Guidelines does not recommend the use of 5-fluorouracil or capecitabine in patients with homozygous DPYD*2A, *13 or rs67376798 [5].

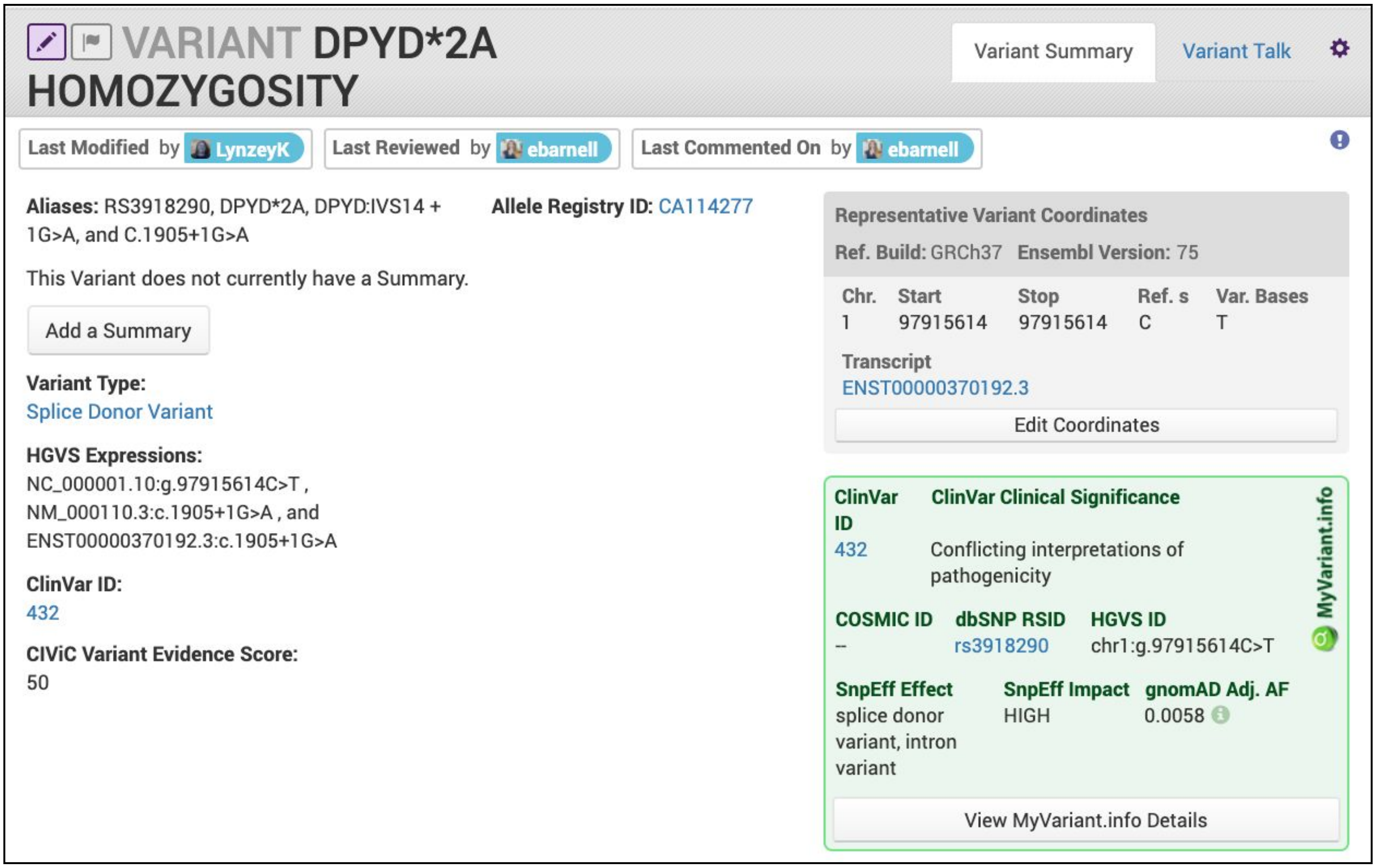


**Curation Practices:**

- Variant Names should be representative of the name described by the publication unless more current descriptions are applicable. In this case, the older name should be added to the list of Variant Aliases.
- Curators should follow naming guidelines provided by applicable associations. This includes recommendations by the Human Genome Variation Society (HGVS), the Human Variome Project (HVP), and the Human Genome Organization (HUGO).

**Supplementary Figure 6. Example of a CIViC variant summary with inclusion of ACMG evidence codes**

The image below depicts the [*TP53* - R175H Variant] which includes germline ACMG evidence codes. In this example, the PM2 code and the specific source of the information used to apply that code are included. This reduces the time needed to evaluate the source of this evidence and whether it warrants updating. Certain evidence codes relating to a variant's population frequency, *in silico* predicted effect, domain location, etc. rely on tools or databases and not specific publications. Such codes relate to the clinical importance of a variant and can appropriately be incorporated at the Variant-level.

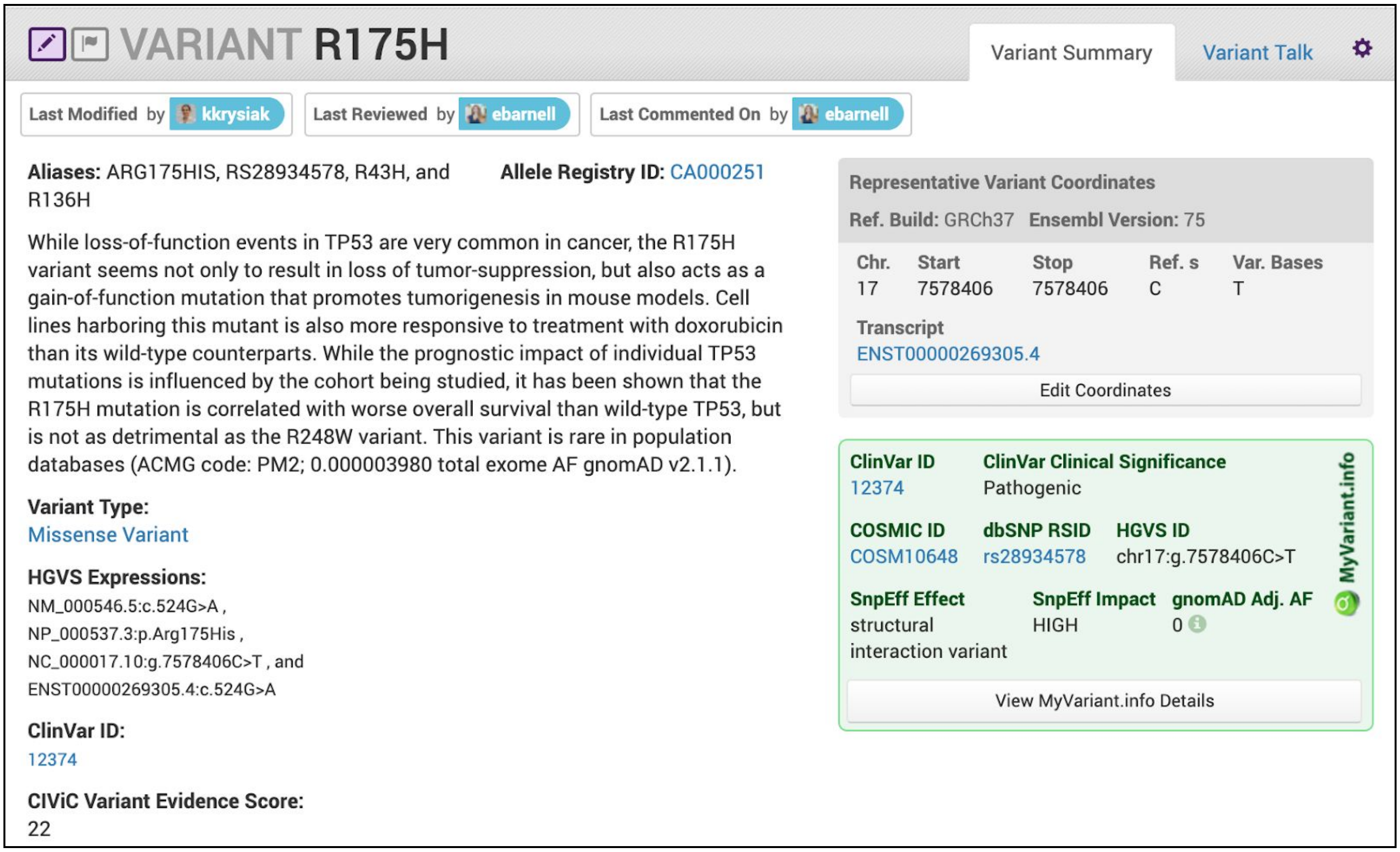


**Curation Practices**

- ACMG codes, which apply independent of disease type (e.g. population allele frequency PM2), may be listed in the Variant Summary.
- Addition of context-specific ACMG evidence codes may also be captured but should be explicit for circumstances in which the code should, or should not, be used (e.g., PM3 for use only in a recessive disorder).
- Variant-level cautions or warnings of inappropriate codes can be incorporated here (e.g., a warning that PVS1 should not be used for a variant due to an alternative start site after the variant's amino acid).

**Supplementary Figure 7. Defining variant coordinates for SNVs and small indels**

Variant annotation for single nucleotide variants (SNV) and small insertion/deletions (indels) follow a 1-based coordinate system and utilize left-shifted normalization. Reference positions are indicated in green and variant positions in purple. Below, each representation is the text that would be entered into the CIViC Reference Base and Variant Base fields in the Variant Suggested Revision form.

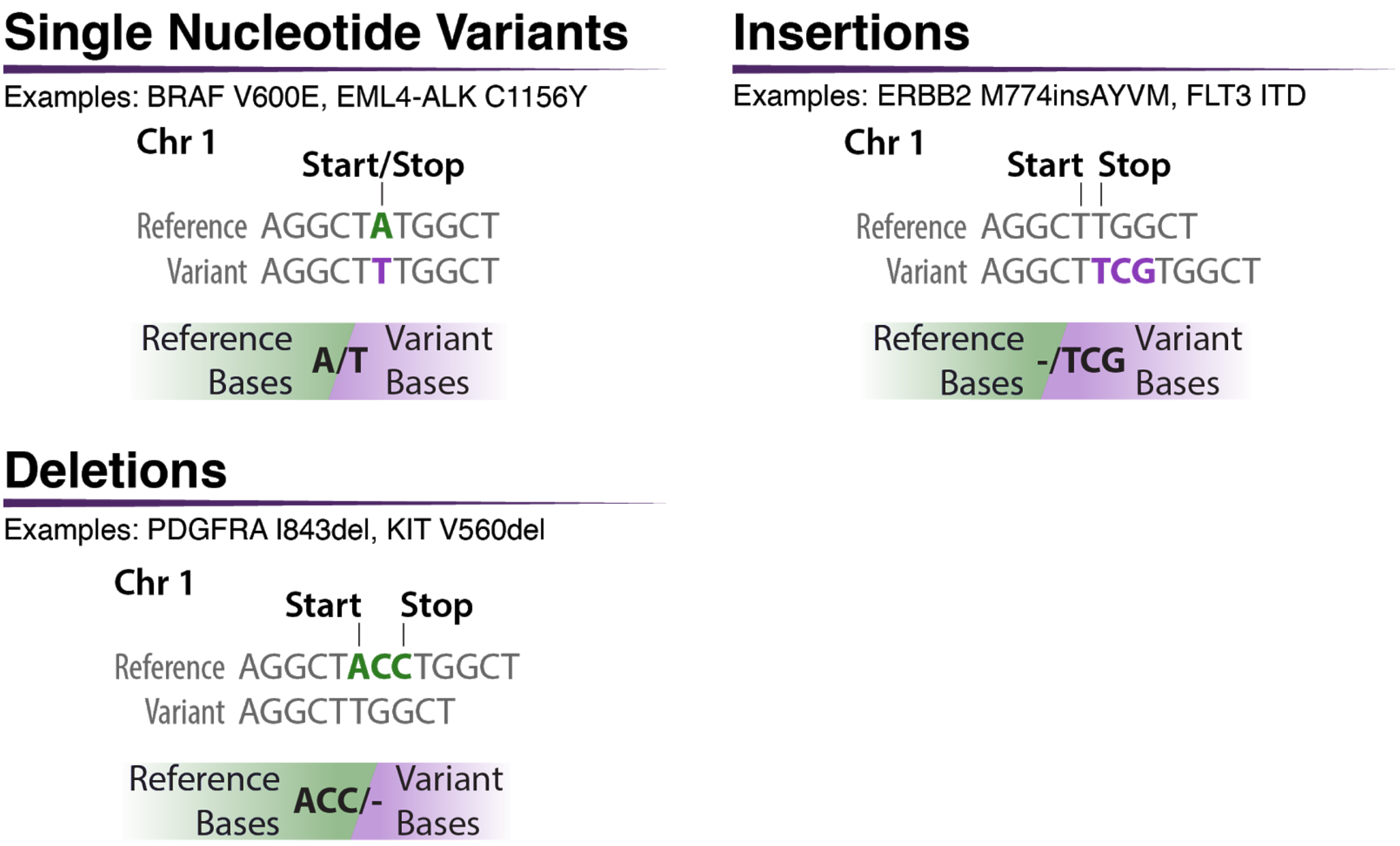


**Curation Practices:**

- When selecting a representative variant, utilize the most specific and recurrent variant, whenever possible. For example, there are more than 70 insertions listed in COSMIC [6] that lead to the highly recurrent *NPM1* W288fs mutation; however, one 4bp insertion (also known as NPM1-A) accounts for more than 90% of all variant entries. Therefore, the coordinates associated with the NPM1-A variant were chosen as the representative coordinates for the *NPM1* W288fs variant in CIViC.
- For complex variants such as SNVs in genes involved in fusions (e.g., EML4-ALK C1156Y), enter the genomic position of the SNV.
- Categorical variants involving a single amino acid can be indicated by using the three or two base pairs of the corresponding triplet codon that could result in a SNV at that site (see BRAF V600).

**Supplementary Figure 8. Defining variant coordinates for categorical and large-scale variants**

Variant annotations for large-scale rearrangements or categorical variants aim to utilize the minimum genomic space that encompasses the range of variants observed for that gene. For example, although *PIK3CA* amplification can encompass much larger genomic coordinates, the outermost coordinates of the gene *PIK3CA* are used to define the start and stop coordinates. Similarly, categorical variants that contain mutations within the same domain or exon use the outermost coordinates of that domain or exon. Fusion coordinates are based on the closest exon boundary included in the fusion. Although multiple breakpoints can occur, the most common breakpoint is prefered for the representative coordinate.

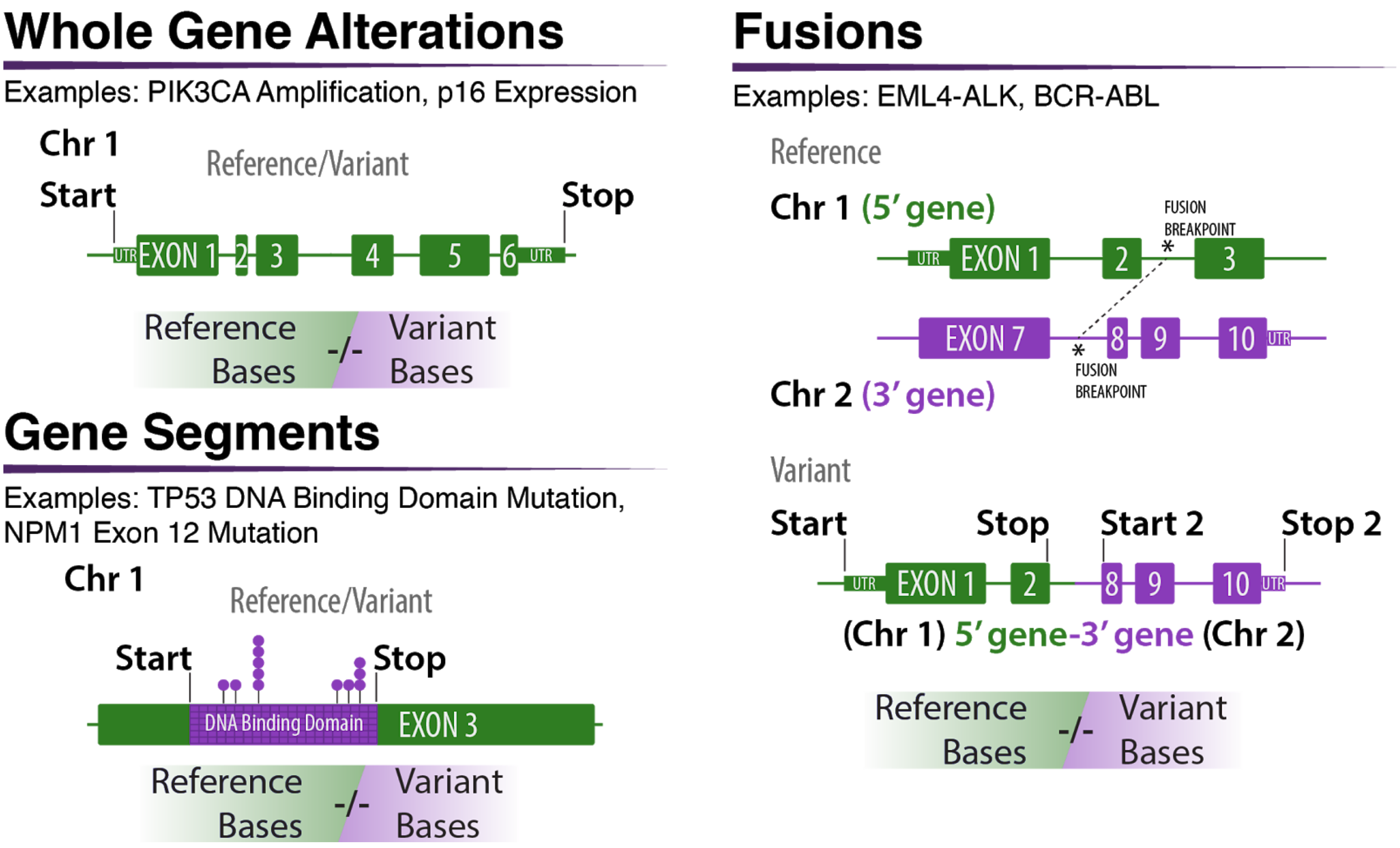


**Curation Practices:**

- Use coordinates that would encompass the most variants that fit the variant name / description to aid others using coordinates to find relevant and similar variants.
- For fusions:
  - Names are always written in a 5’ to 3’ order (e.g. 5’ *BCR-ABL* 3’);
  - The Variant is placed under the most ‘important’ gene - e.g. kinase domain - (not repeated under both) which is often the 3’ gene;
  - Coordinates represent the entire putative fusion transcript including start to end of 5’ transcript fusion partner (primary coordinates) and start to end of 3’ transcript fusion partner (secondary coordinates).

**Supplementary Figure 9. Choosing a representative transcript**

Genes often have multiple transcript representations. CIViC utilizes Ensembl v75 for transcript annotations. The representative transcript for *WT1* depicted in purple below was chosen because it has the widest outer coordinates with the most common exons compared to the other transcripts depicted in green. This transcript is further highlighted by *** because it is also designated as the "canonical transcript" by Ensembl using select criteria defined in their glossary of terms.

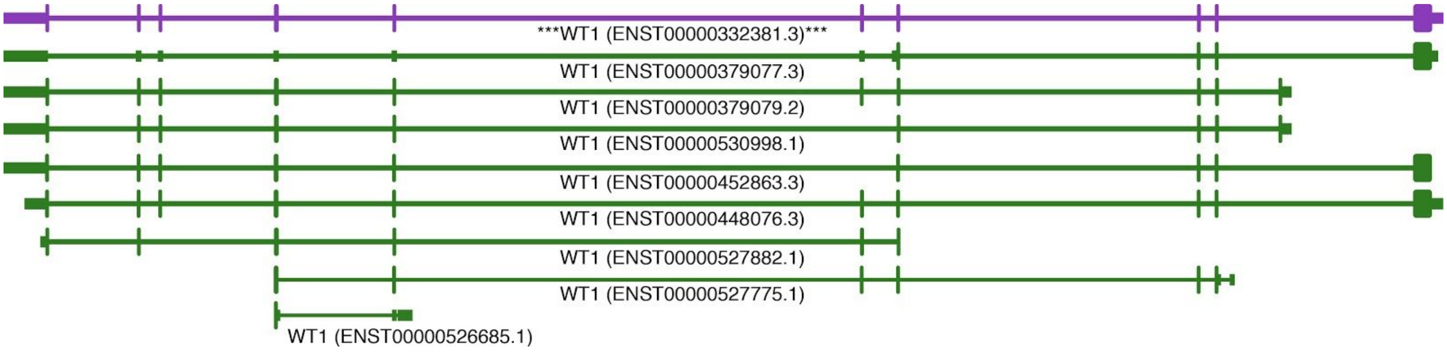


**Curation Practices:**

- There is no one 'right' answer for representative transcript.
- It must:
  - contain the variant (except in rare cases like promoter mutations);
  - be based on an Ensembl transcript and include the transcript version.
- It may:
  - be the transcript with the longest ORF or most exons;
  - be the transcript that contains the 'canonical exons' that are used in many transcripts;
  - be the variant that has the greatest outer coordinates;
  - be the transcript that is widely used in literature;
  - be a transcript that is compatible with interpretation/visualization in the primary literature source.
- An IGV reference transcript file containing Ensembl (v75) transcripts can be obtained here: https://civicdb.org/downloads/Ensembl-v75_build37-hg19_UcscGenePred_CIViC-Genes.ensGene
  - Ensembl canonical transcripts are designated by ***.
- Selection of Representative Transcripts for intronic or regulatory variants follow a similar pattern as protein coding variants.

### The Evidence Item data model

**Supplementary Figure 10. Overview of CIViC Evidence Item entry form**

The CIViC evidence entry form reacts to user input with type-ahead search, controlled vocabularies, reactive help text and field layouts updated based on user choices.

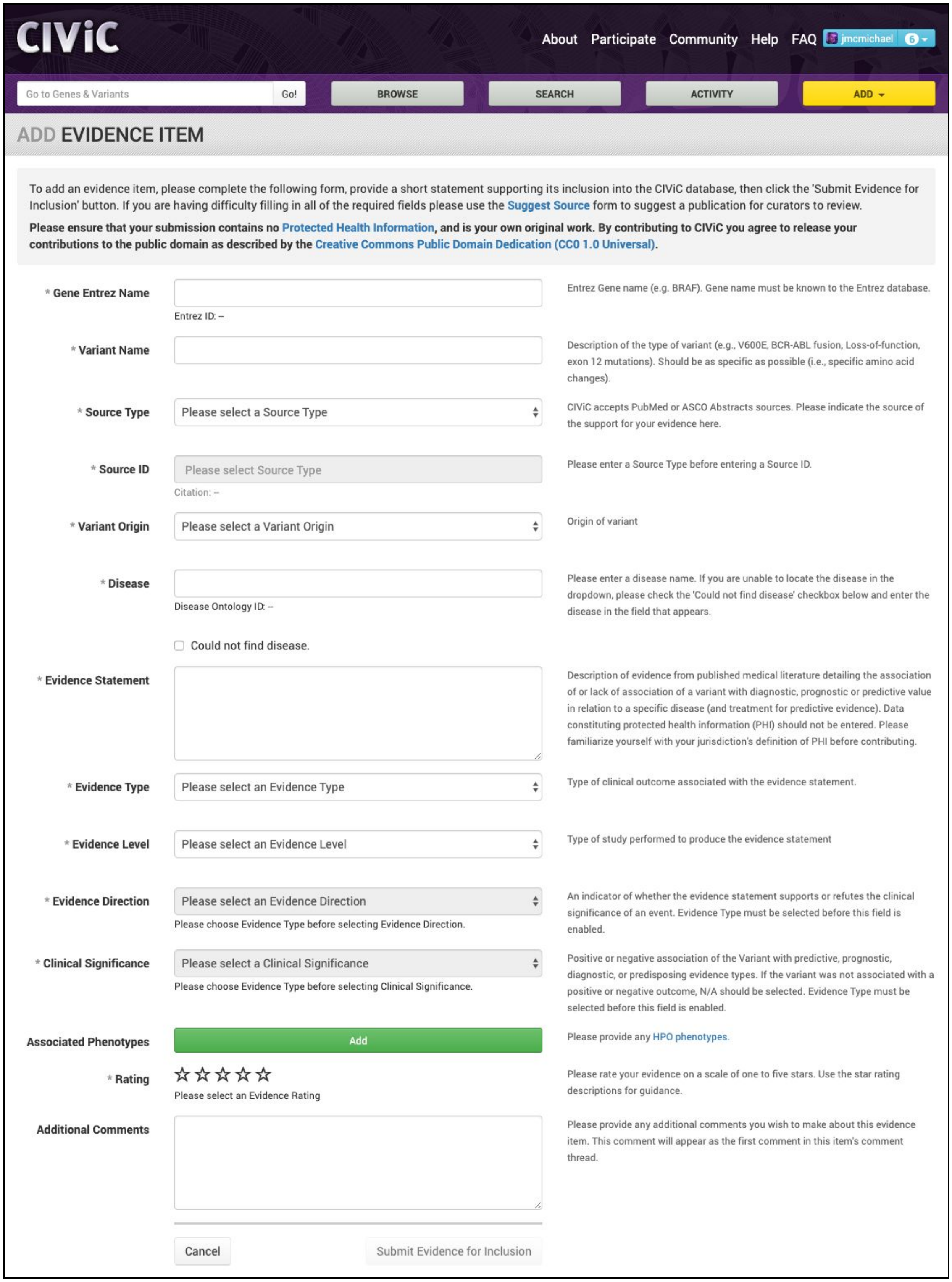

**Supplementary Figure 11. View of the Evidence Grid for a given Variant**

Evidence for a given Variant is displayed in an Evidence Grid. This grid is highly customizable with each field allowing for text-based searching (e.g., EID, Description), entity filtering (e.g., Evidence Level), and sorting. The variety of evidence for a Variant can be quickly consumed using the combination of colors and icons (see “Help” for full legend). The current curation state of the Evidence Item including status and any pending revisions are displayed by the color of the EID and the presence of the Pending Revisions icon. This curation state should be considered when consuming data for any Variant in CIViC. By default the grid shows Accepted and Submitted evidence but all evidence can be viewed by using the “Grid Options Menu” (top right). Downloading the contents of the grid can be quickly achieved through the “Get Data” or “Grid Options Menu.”

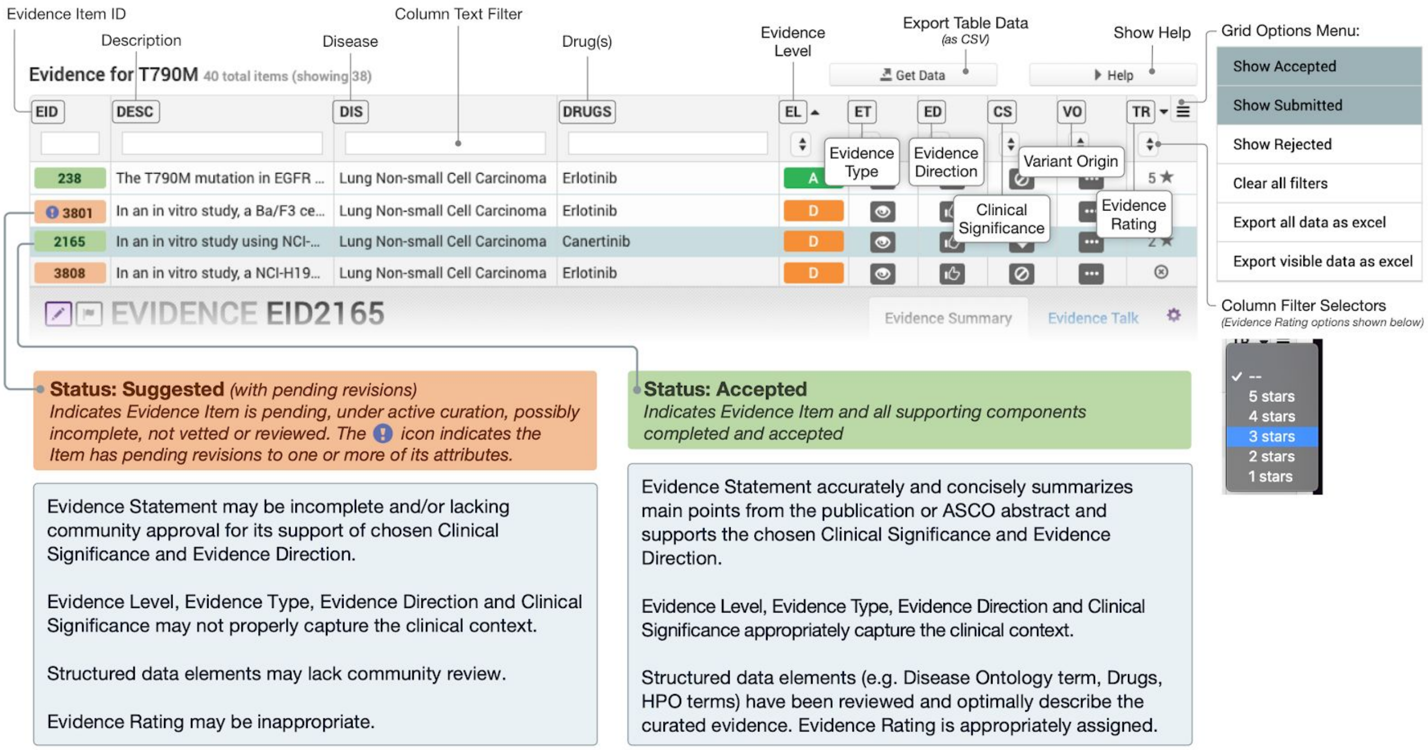


**Curation Practices:**

- Evidence Items should generally be prepared from primary literature rather than from review articles. It is recommended that curators use reviews to identify primary literature referenced in the review and curate individual Evidence Items based on these cited articles. Reference articles can also be used to develop Gene and Variant summaries.
- When curating new evidence, the curator should keep in mind the existing evidence for that Variant and Evidence Type.
    - For clinical trials and case reports (Levels A, B, and C), overlapping patient populations should be avoided, if possible, or carefully noted to alert users of this nuance and avoid conclusions that mistake these studies as independent.
- Disease stage, prior treatments, and other experimental details influencing evidence interpretation should be captured within an Evidence Item to maximize user comprehension of the underlying study and the appropriate context in which it is relevant. Such details are critical parts of clinical guidelines and can impact which clinical guidelines should be used as well as drug sensitivities (see EID1008 and EID1009).

### Examples of Variant Origin

**Supplementary Figure 12. Example of Evidence Item where the Variant Origin is not applicable (N/A)**

Below is an Evidence Item that support sensitivity/response to Trastuzumab for a patient with breast cancer and an *ERBB2* - Amplification variant. In this example, the clinical trial describing this Evidence Item refers to patients with either Amplification or Overexpression of ERBB2. In the case of over-expression (measured by IHC) or Amplification (measured by FISH), typically only tumor samples are assayed. Thus it is not possible to definitely state that the variant is of somatic origin. Furthermore, in some cases over-expression may be driven by epigenetic causes where a variant does not apply. For these reasons, the variant origin has been entered as N/A.

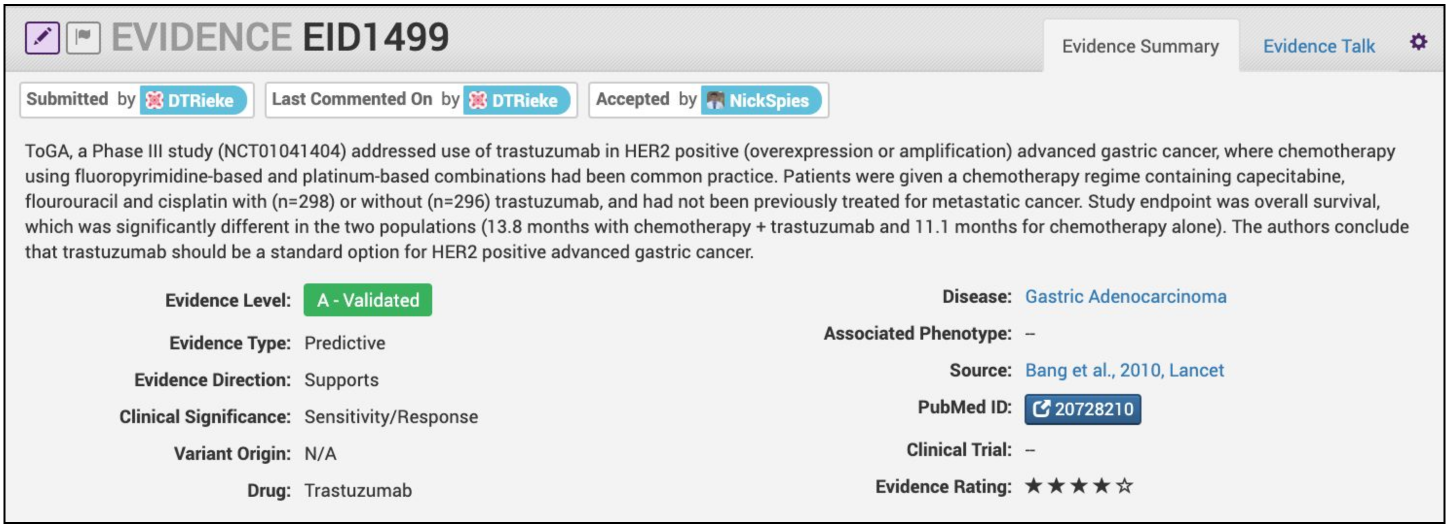


**Curation Practices:**

- N/A Variant Origin is particularly used in cases that involve differences in expression, methylation, or other post-translational modifications.
- Variants characterized in functional Evidence Items with *in vitro* data will usually have N/A Variant Origin.

### Examples of Evidence Levels

**Supplementary Figure 13. Example of a A-level (Validated) Evidence Item**

Below is an example of an A-level (Validated) Evidence Item for the *BRAF* - V600E variant. In this example, the Evidence Item is describing the Phase 3 randomized clinical trial that was submitted to the FDA for therapeutic approval of Vemurafenib with Dacarbazine for treatment of untreated, metastatic melanoma.

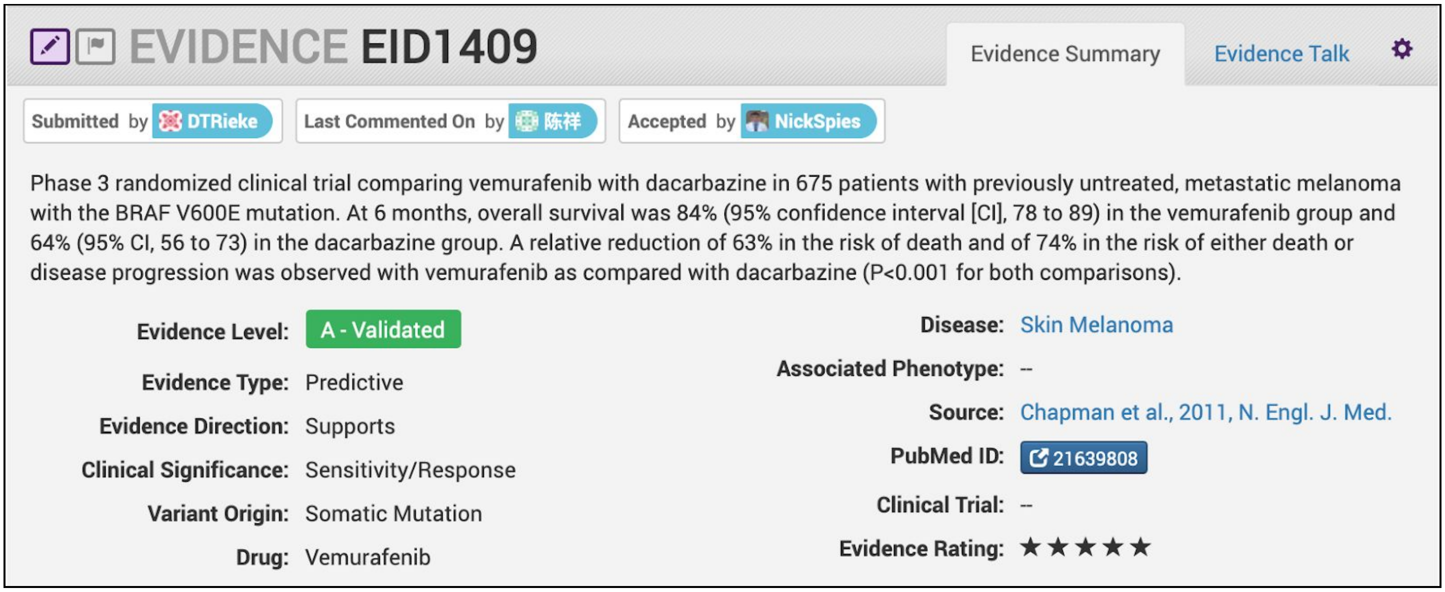


**Curation Practices:**

- Typically, A-level Validated Evidence Items describe Phase III Clinical Trials (for therapeutics or companion diagnostics), which are subsequently submitted to the FDA for pre-market approval.
- In general, Evidence Items derived from any study cited in approvals, established practice guidelines, or considered the definitive practice-changing study, may be labeled Level A. In some cases Phase I trials can meet this requirement (see EID1187).
- Evidence Statements should include the gene/variant being evaluated, the study population, disease state, study size, statistical significance (e.g., p-value, confidence interval), duration of the study, and other relevant information that is required to assess the evidence for variant interpretation.
- Evidence Items derived from publications describing practice guidelines (e.g. WHO diagnostic criteria) are labeled A-Validated Evidence Level.

**Supplementary Figure 14. Example of a B-level (Clinical) Evidence Item**

Below is an example of a B-level (Clinical) Evidence Item for the *BRAF* - V600E variant. In this example, the Evidence Item is describing a Phase 2 randomized clinical trial that was used to assess preliminary efficacy of the use of Vemurafenib for treatment of patients with previously treated skin melanoma.

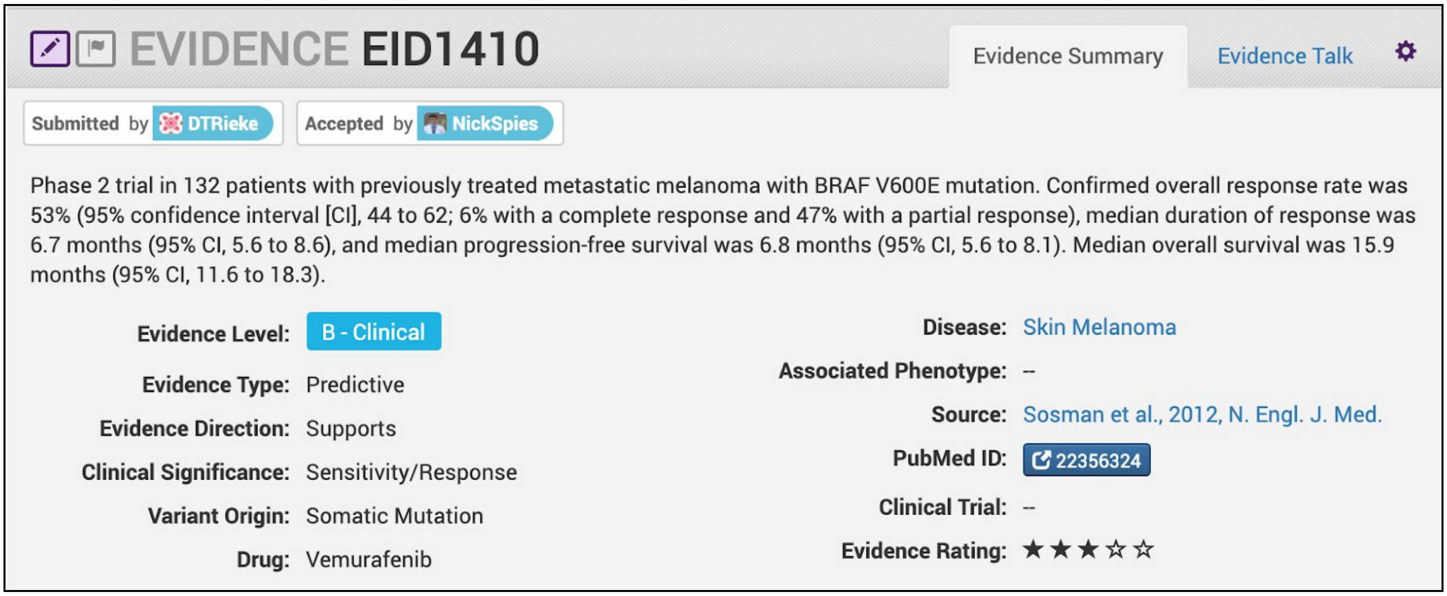


**Curation Practices:**

- B-level Evidence Items can describe trials submitted to the FDA during the approval process; however, relative to A-level Evidence Items, B-level Evidence Items typically have a smaller sample size or assess less significant outcomes (e.g., response rate instead of overall survival).
- Phase I, II, and III clinical trials make up a significant percentage of Level B Evidence Items.
- For curation of Phase I evidence, notes on treatment related adverse events may be added to the main evidence statement describing the variant-positive patient subgroup response to treatment, as dosing and adverse events are among the main focuses of Phase I studies.
- B-level Evidence Items do not have to be derived from clinical trials but also can describe studies which attain a sufficient sample size to be considered more informative than a series of case studies, and ideally have some component of statistical conclusions in their results.
- Greater than five patients are typically required for an Evidence Item to be considered a B-level Evidence Item.
- Evidence Statements should include the gene/variant being evaluated, the study population, disease state, study size, statistical significance (e.g., p-value, confidence interval), duration of the study, and other relevant information that is required to assess the evidence for variant interpretation.
- Categorical variants (sometimes called bucket variants colloquially) often appear in B-level Evidence Items describing clinical trials, which pool together patient populations with mutations of a certain class (e.g. “PIK3CA mutation”), in order to attain a disease specific, statistically significant, clinical result across the patient population (e.g. Trastuzumab resistance in HER2 positive breast cancer).

**Supplementary Figure 15. Example of a C-level (Case Study) Evidence Item**

Below is an example of a C-level (Case Study) Evidence Item for the *BRAF* - V600E variant. In this example, the Evidence Item is describing a single patient with the *BRAF* - V600E variant who demonstrated sensitivity/response to Pictilisib in the disease context of melanoma. This Evidence Item was classified as a Case Study because it described results for a single patient with advanced melanoma who had been enrolled in a larger Phase I clinical trial that evaluated 60 patients with advanced solid tumors and any *BRAF* variant for sensitivity to Pictilisib.

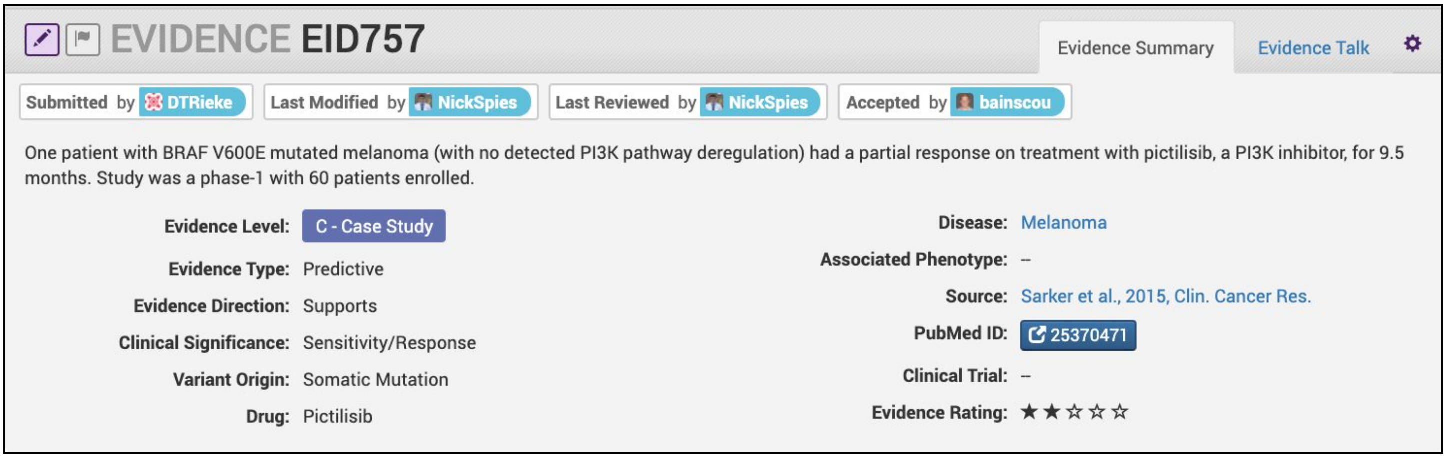


**Curation Practices:**

- C-level Evidence Items should describe a specific variant and likely will not apply to a categorical variant.
- In some cases a clinical trial employing a categorical or bucket variant (e.g. EGFR mutation) will contain additional supplementary information on individual patient mutations and outcomes (e.g. CR, PR, SD or PD as best response). In such cases, along with the B-level Evidence Item based on the categorical variant, individual C-level case study Evidence Items can be curated for each listed variant.
- Evidence Items involving fewer than five patients are typically considered to be C-level Evidence Items.
- Evidence Statements should include the gene/variant being evaluated, the study population, disease state, study size, statistical significance (e.g., p-value, confidence interval, if applicable), duration of the study, and other relevant information that is required to assess the evidence for variant interpretation.

**Supplementary Figure 16. Example of a D-level (Preclinical) Evidence Item**

Below is an example of a D-level (Preclinical) Evidence Item for the *BRAF* - V600E variant. In this example, 49 BRAF-mutant melanoma cell lines exhibited resistance to a combination of dactolisib and selumetinib treatment. Note that older drug names were used in this study, BEZ238 and AZD6244, but since then, the drug names have been updated to dactolisib and selumetinib. To reduce confusion, the more current names are used in the drug field and the curator has included both the old and new names in the Evidence Statement.

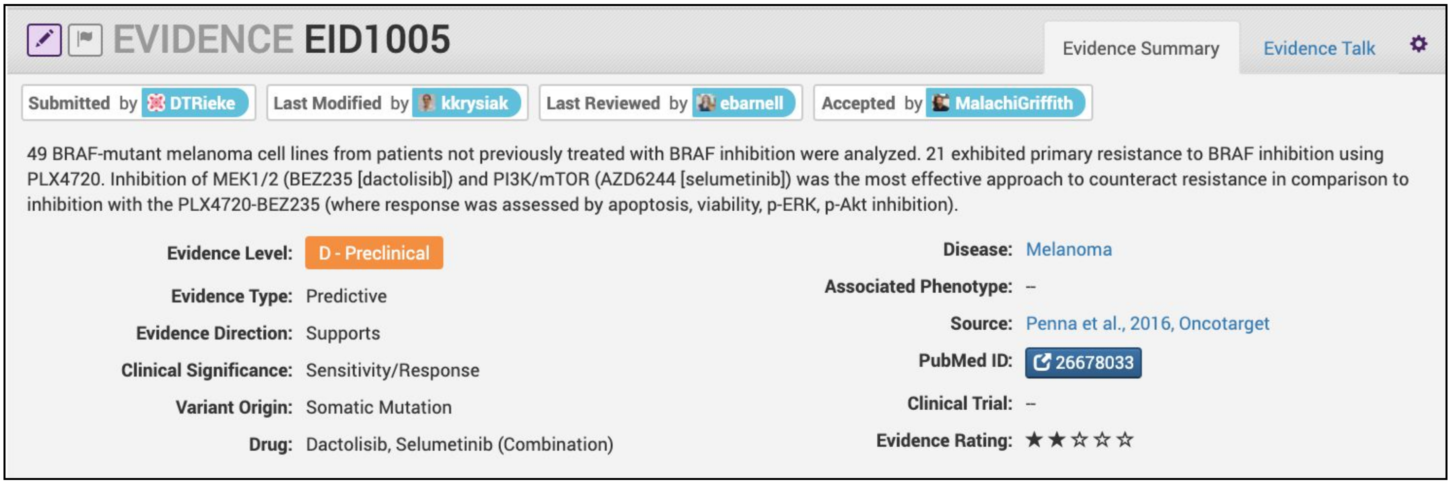


**Curation Practices:**

- D-level Evidence Items typically describe animal models or cell line studies. The sample size for these studies can influence the Trust Rating, whereby increased numbers of mice or independent biological replicates used should increase the Trust Rating.
- A concise description of the experiments performed should be prepared by the curator, supporting the Evidence Item Clinical Significance, and describing the controls that were used, and the significant findings that were observed.
- Evidence Statements should include the gene/variant being evaluated, the study population, disease state, study size, statistical significance (e.g., p-value, confidence interval), duration of the study, and other relevant information that is required to assess the evidence for variant interpretation.

**Supplementary Figure 17. Example of an E-level (Inferential) Evidence Item**

Below is an example of an E-level (Inferential) Evidence Item for the *BRAF* - V600 Amplification variant. In this example, the Evidence Item is describing how *BRAF* - V600E Amplification could be a mechanism of selumetinib resistance in patients with colorectal cancer.

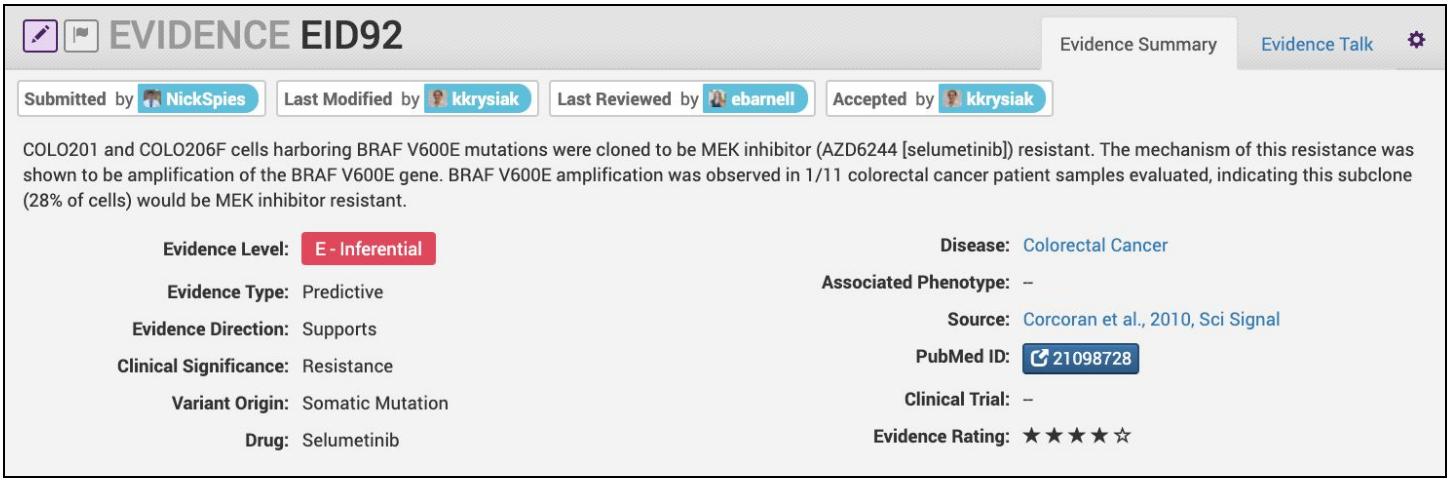


**Curation Practices:**

- E-level Evidence Items provide inferential support for the described variant. This could mean that the variant was not ever actually measured, or that the results from the study do not directly evaluate the claims made by the Evidence Item.
- E-level Evidence Items can be derived from *in silico* predictions, cell lines, animal models, or human studies.
- Evidence Statements should include the gene/variant being evaluated, the study population, disease state, study size, statistical significance (e.g., p-value, confidence interval), duration of the study, and other relevant information that is required to assess the evidence for variant interpretation. Often these data are not available for E-level Evidence Items.

### Examples of Evidence Types

**Supplementary Figure 18. Example of a Predictive Evidence Type**

Below is an example of an Evidence Item that illustrates the Predictive Evidence Type. This example describes the CLEOPATRA trial (NCT00567190), which evaluated 808 patients with HER2-positive metastatic breast cancer. These patients demonstrated significant sensitivity/response when treated with combination therapy of docetaxel, pertuzumab and trastuzumab.

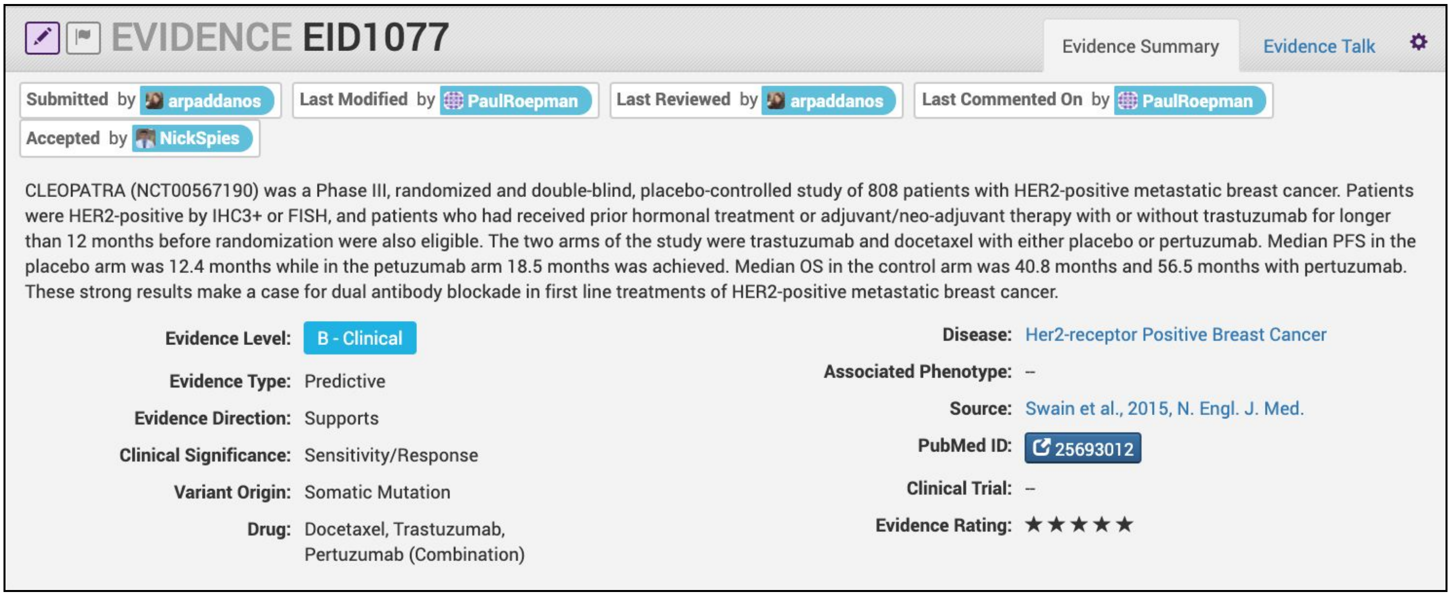

EVIDENCE EID1077

Evidence Summary | Evidence Talk

Submitted by arpaddanos | Last Modified by PaulRoepman | Last Reviewed by arpaddanos | Last Commented On by PaulRoepman | Accepted by NickSpies

CLEOPATRA (NCT00567190) was a Phase III, randomized and double-blind, placebo-controlled study of 808 patients with HER2-positive metastatic breast cancer. Patients were HER2-positive by IHC3+ or FISH, and patients who had received prior hormonal treatment or adjuvant/neo-adjuvant therapy with or without trastuzumab for longer than 12 months before randomization were also eligible. The two arms of the study were trastuzumab and docetaxel with either placebo or pertuzumab. Median PFS in the placebo arm was 12.4 months while in the petuzumab arm 18.5 months was achieved. Median OS in the control arm was 40.8 months and 56.5 months with pertuzumab. These strong results make a case for dual antibody blockade in first line treatments of HER2-positive metastatic breast cancer.

| | | | |
|---|---|---|---|
| Evidence Level: | B - Clinical | Disease: | Her2-receptor Positive Breast Cancer |
| Evidence Type: | Predictive | Associated Phenotype: | -- |
| Evidence Direction: | Supports | Source: | Swain et al., 2015, N. Engl. J. Med. |
| Clinical Significance: | Sensitivity/Response | PubMed ID: | 25693012 |
| Variant Origin: | Somatic Mutation | Clinical Trial: | -- |
| Drug: | Docetaxel, Trastuzumab, Pertuzumab (Combination) | Evidence Rating: | ★★★★★ |

**Curation Practices:**

- Predictive Evidence Items should include the Drug Name(s) and Drug Interaction Type (for multiple drugs).
- The most current name of the Drug (excluding trade names) should be used in the Drug field to reduce duplication. The Evidence Statement should contain the drug name used in the study with the current name in brackets, when applicable (see **Supplementary Figure 15** for an example).
- Drug Interaction Types are required anytime more than one drug is mentioned for a given study. If multiple drug interaction types are at play (e.g., combinations and substitutes), considering separating these concepts into more than one Evidence Item.
- If applicable, the Clinical Trial name and ID should be included in the Evidence Statement. Any clinical trial IDs available in PubMed for the Source linked to this Evidence Item will be automatically imported and linked to this Evidence Item when the PubMed Source is imported into CIViC.
- The duration of exposure to the drug and confounding interactions (e.g., wash-out periods, previous treatment, cancer stage) should be listed.
- Assigning a Clinical Significance of Sensitivity/Response can depend on factors such as response rate, which will vary significantly with disease and treatment. In some cases a response rate of 15% may represent a significant improvement, and merit a valuation of Sensitivity/Response. A general guideline for CIViC curation is to follow the author's published (and peer-reviewed) interpretations and conclusions of the results.
- Extensive guidelines, use cases, and examples for curation of predictive evidence are given in **Supplementary Figure 22** and **Supplementary Table 4**.

**Supplementary Figure 19. Example of a Diagnostic Evidence Type**

Below is an example of an Evidence Item that illustrates the Diagnostic Evidence Type. This example describes the World Health Organization guidelines for classifying chronic myelomonocytic leukemia (CMML). Specifically, if a patient has a PCM1-JAK2 fusion or a rearrangement involving PDGFRA, PDGFRB, or FGFR1, especially in the setting of eosinophilia, the patient does not have CMML.

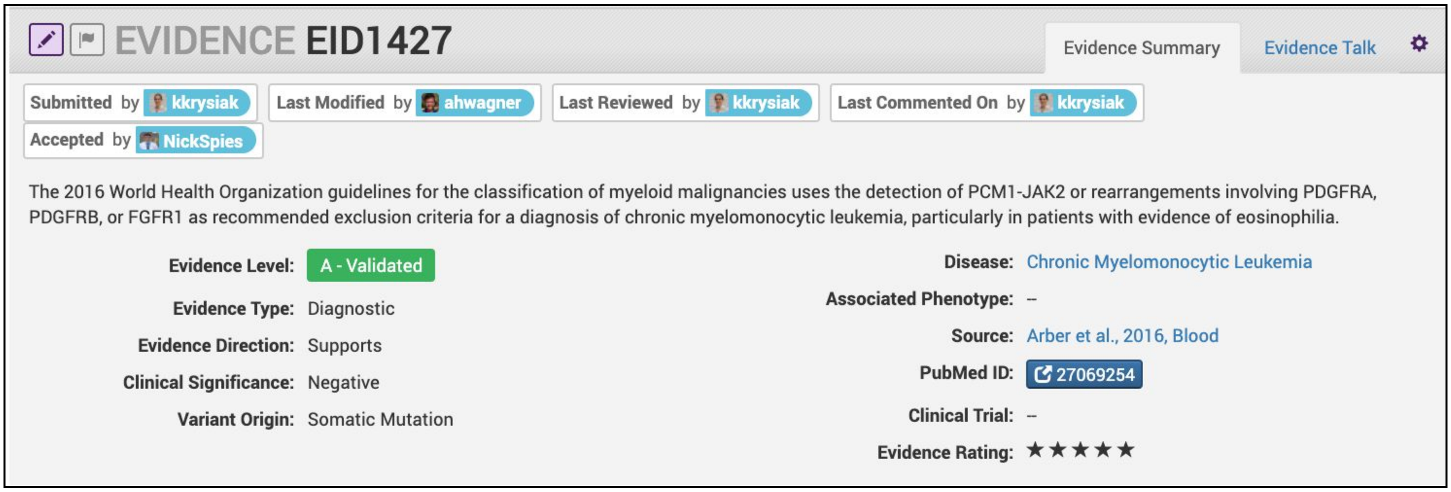


**Curation Practices:**

- Diagnostic Evidence Items should only be used if the variant assists in labeling the patient with a specific disease or disease subtype and should not be used to denote that the particular variant is prevalent in a specific disease.
- Generally, Diagnostic Evidence Items describe variants that can help accurately diagnose a cancer type or subtype with high sensitivity and specificity, for which diagnoses may otherwise be challenging.
- Diagnostic Evidence Items are very closely tied to the terms of the Disease Ontology (DO) in CIViC. The Disease Ontology works to actively generate mappings to other highly used ontologies, but the terms in the DO are generally accepted diseases which are part of medical practice. Therefore, literature proposing a novel disease type - for instance studies suggesting a novel cancer subtype defined by the presence of a specific oncogenic variant - are not generally admitted as part of the CIViC data model. Alternatively, if a curator with expertise in the field feels that the novel subtype has met with a sufficient level of acceptance, they may submit this type of Evidence Item using a non-DO term, and suggest that DO admit this term into the ontology.
- Literature describing diagnostic practice guidelines (such as those of the World Health Organization) may be used in curation and submitted as A-level Evidence Items.
- Literature describing small numbers of observations in patient samples of a certain variant, where the authors state that the variant may have diagnostic value, may be admitted as lower star Case Study (C-level) data. Similar literature employing larger numbers could be labeled as Clinical (B-level).
- Guidelines and use cases for curation of diagnostic evidence are given in **Supplementary Table 4**.

**Supplementary Figure 20. Example of a Prognostic Evidence Type**

Below is an example of an Evidence Item that describes a Prognostic Evidence Type. This example describes a 406-patient trial whereby observation of any somatic *TP53* mutation in chronic lymphoblastic leukemia conferred poor prognosis relative to wildtype *TP53*.

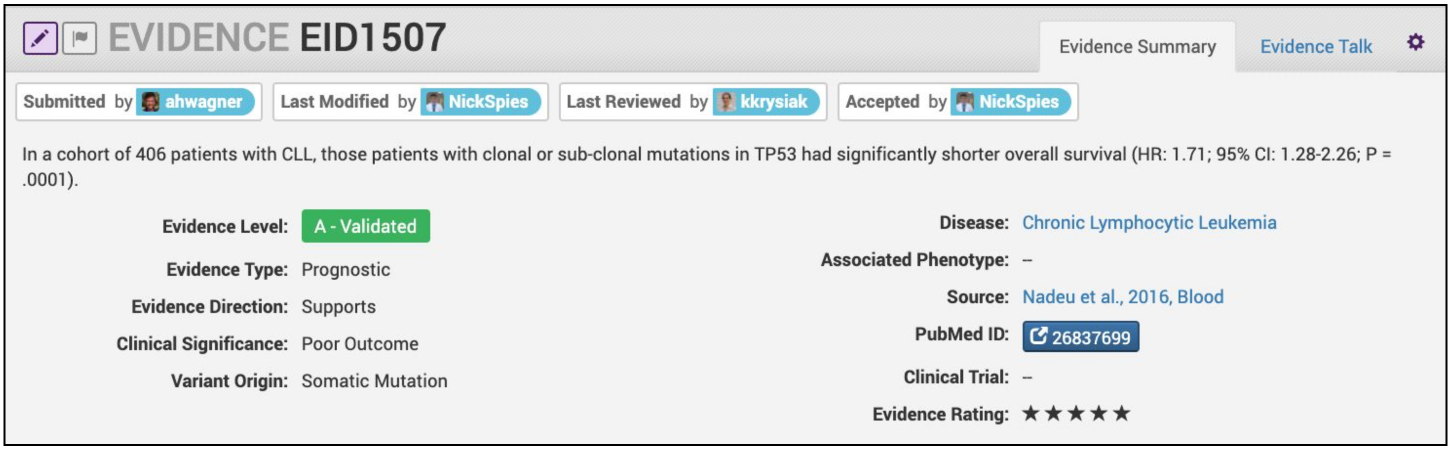


**Curation Practices:**

- Prognostic Evidence Items should include the measured outcome (e.g., overall survival, complete response, partial response), number of subjects and applicable statistics.
- If described in the literature, a definition of the measured outcome should be given.
- Prognostic evidence is characterized by either better outcomes for patient subpopulations with the given variant, which are not specific to any particular treatment context, or worse outcomes which are not indicative of variant resistance to a specific treatment. Instead, the change in outcome should be largely correlated to the presence of the variant.
- In some cases, a variant subpopulation with worse outcome may benefit from subsequent therapy targeted to that variant (e.g., HER2 amplification in breast cancer).
- Guidelines, use cases, and examples for curation of prognostic evidence are given in **Supplementary Figure 22** and **Supplementary Table 4**.

**Supplementary Figure 21. Example of a Predisposing Evidence Type**

Below is an example of an Evidence Item that describes a Predisposing Evidence Type. This example describes a case study of two families (9 total patients) with *VHL* - Q195* (c.583C>T) variants. All patients in this cohort were diagnosed with Von Hippel-Lindau Disease. Associated phenotypes for these patients included cerebellar hemangioblastomas, pheochromocytomas, renal cell carcinoma, and retinal capillary hemangioma. In this example, the ACMG code was PVS1 and Clinical Significance was Uncertain.

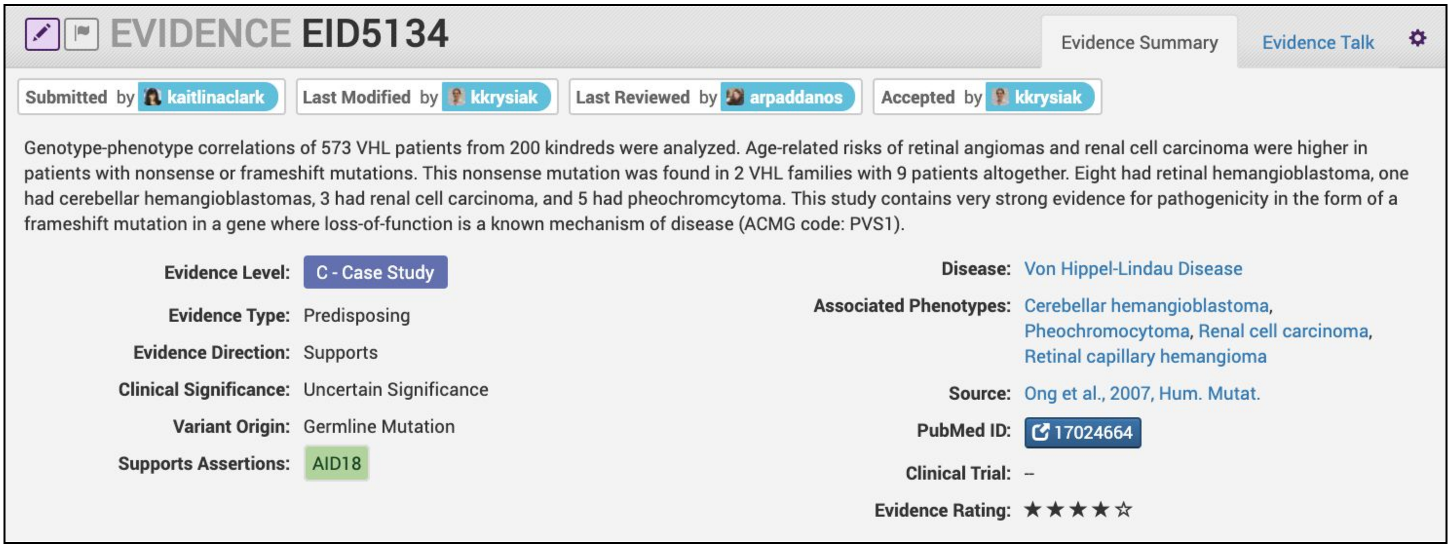


**Curation Practices:**

- Typically, but not always, Predisposing Evidence Items are Germline Mutations or Germline Polymorphisms. In rare circumstances, the patient can have a predisposing variant that develops as a result of a somatic mutation or mosaicism during embryogenesis that is widespread but not necessarily heritable.
- ACMG evidence codes (ACMG criteria) are derived from the evidence presented in the specific Source and are listed at the end of the Evidence Statement with a brief justification for each code's use.
- ACMG evidence codes not directly derived from Source associated with the Evidence Item (e.g. population databases for PM2) are captured at the Variant Summary (**Supplementary Figure 6**) or Assertion level (**Figure 4**).
- A pathogenicity valuation is made for each Predisposing Evidence Item (EID) on a per-Evidence Item basis, using only the ACMG evidence codes listed in the EID and derived from the EID evidence source. The above example lists PVS1 as derived from the literature source resulting in the Clinical Significance of Uncertain Significance (using ACMG classification criteria), reflecting that isolated literature source. At the Assertion level, ACMG codes are combined from multiple Evidence Items and other sources to classify the variant based on the field-wide state of knowledge for the variant and disease (**Figure 5B**).

**Supplementary Figure 22. Example of a Functional Evidence Type**

Below is an example of an Evidence Item that describes a Functional Evidence Type. This example summarizes the impact of a novel *KIAA1549-BRAF* fusion event on the function of the *BRAF* protein. Specifically, the fusion product showed gain of function activity in cell lines relative to wildtype kinase. This activity was also demonstrated to be comparable to a known gain of function variant, *BRAF* V600E.

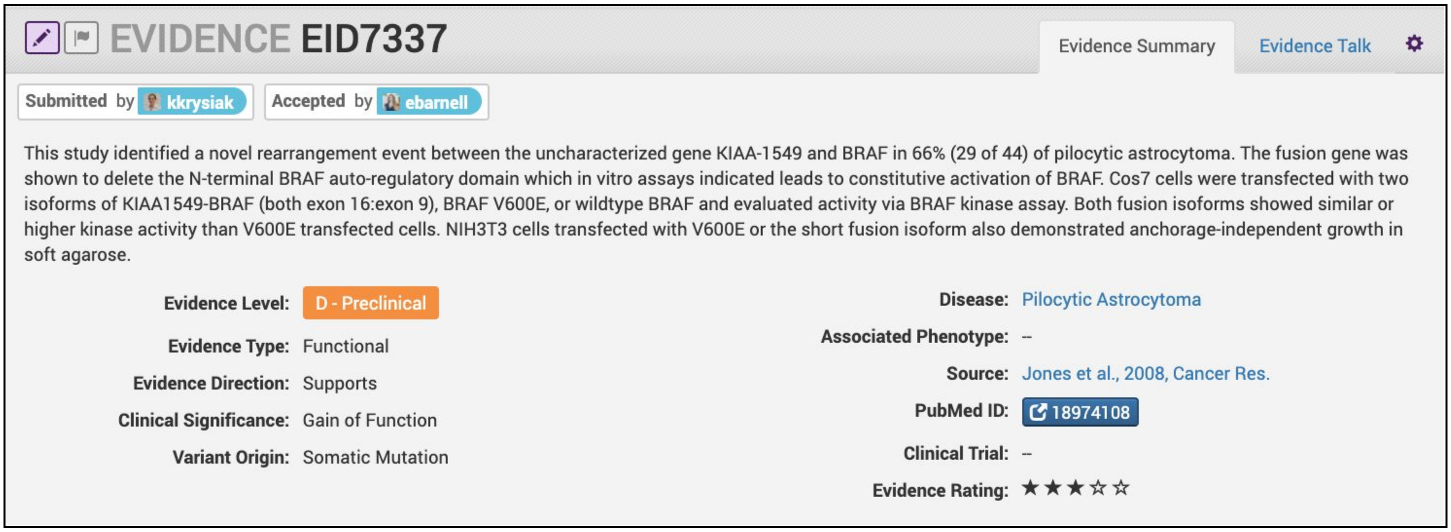


**Curation Practices:**

- Functional Evidence Items should describe how the variant alters biological function from the reference state. This can include a change in function or lack of change in function.
- Clinical Significance for Functional Evidence Types adhere to the following rules:
  - Gain of Function = A variant whereby new/enhanced function is conferred on the gene product;
  - Loss of Function = A variant whereby the gene product has diminished or abolished function;
  - Unaltered Function = A variant whereby the function of the gene product is unchanged;
  - Neomorphic = A variant whereby the function of the gene product is a new function relative to the wildtype function;
  - Unknown = A variant that cannot be precisely defined by gain-of-function, loss-of-function, or unaltered function.
- Functional Evidence Items may be used to support certain ACMG codes (e.g. PM1). In these cases, the ACMG code should be listed in the Evidence Statement along with a brief justification for its inclusion.
- In some cases, Functional Evidence Items may appear as supporting evidence for a Predisposing Assertion, for instance in support of a PM1 evidence code.

**Supplementary Figure 23. Interpreting clinical trial data to curate Predictive/Prognostic Evidence Items**

This figure provides hypothetical examples for interpreting clinical trial data to curate Predictive and Prognostic Evidence Items. Each color represents a different hypothetical 2-armed clinical trial (Treatment vs. Control) where each arm contains wildtype (WT) and mutant (MT) patient populations. Each of the seven studies contains different values for progression free survival (PFS), which are assumed to represent significant differences between populations. The yellow boxes indicate the selections that should be made for the Evidence Type, Evidence Direction, and Evidence Significance. The text below these selections describes the study being evaluated.

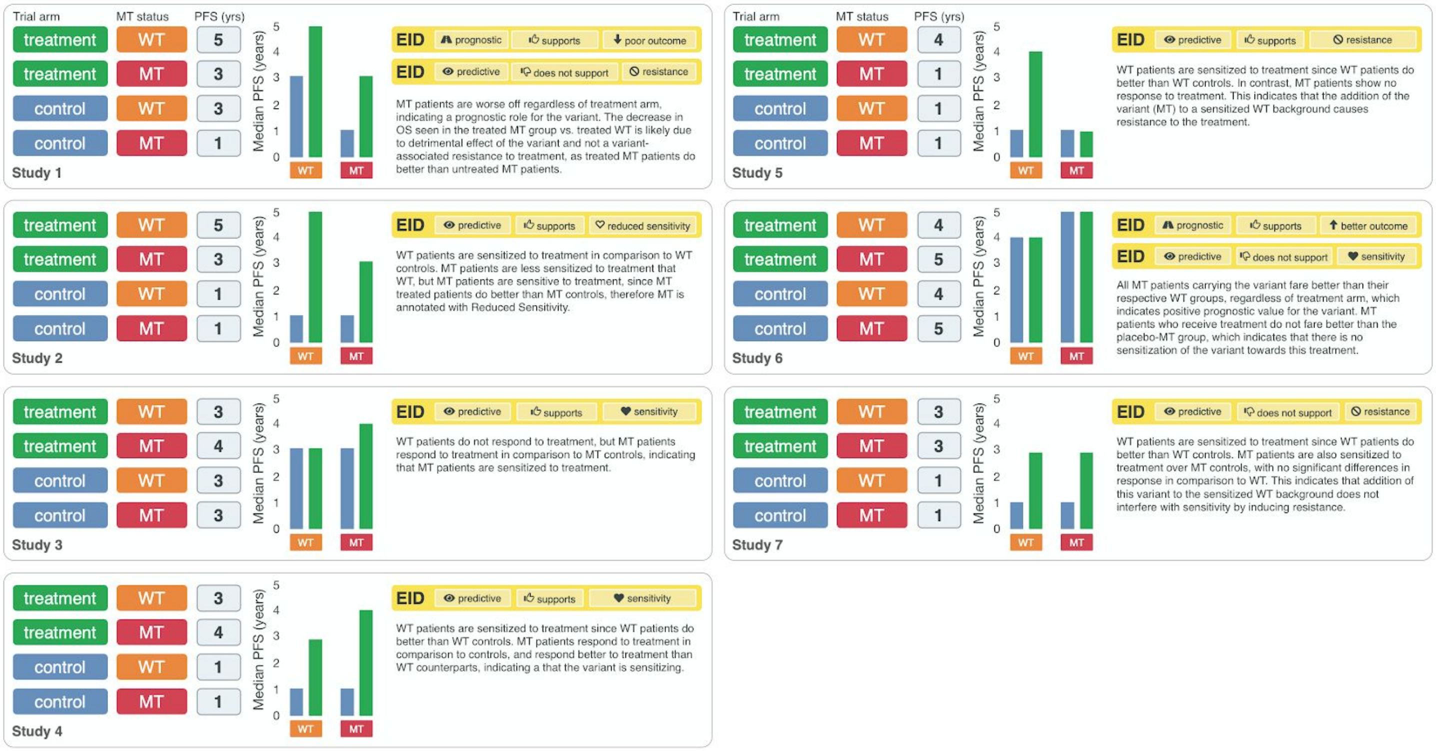


#### Curation Practices:

- These examples do not show all possible ways to obtain the listed Clinical Significances.
- It is recommended to always follow the authors' interpretations of the trial results when writing EIDs since other parameters (e.g. response rate) impact clinical interpretations, often in a disease specific manner.

### Examples of Trust Ratings

**Supplementary Figure 24. Evidence Item with 5-star Trust Rating**

The example Evidence Item below describes a Phase III clinical trial (PROFILE 1014) that evaluated the impact of the *ALK* - Fusions variant on therapeutic response with crizotinib for patients with lung non-small cell carcinoma. The clinical trial was a randomized, double-blinded, placebo-control study of 343 patients that evaluated progression free survival, objective response rate, and quality of life. The results were published in the New England Journal of Medicine.

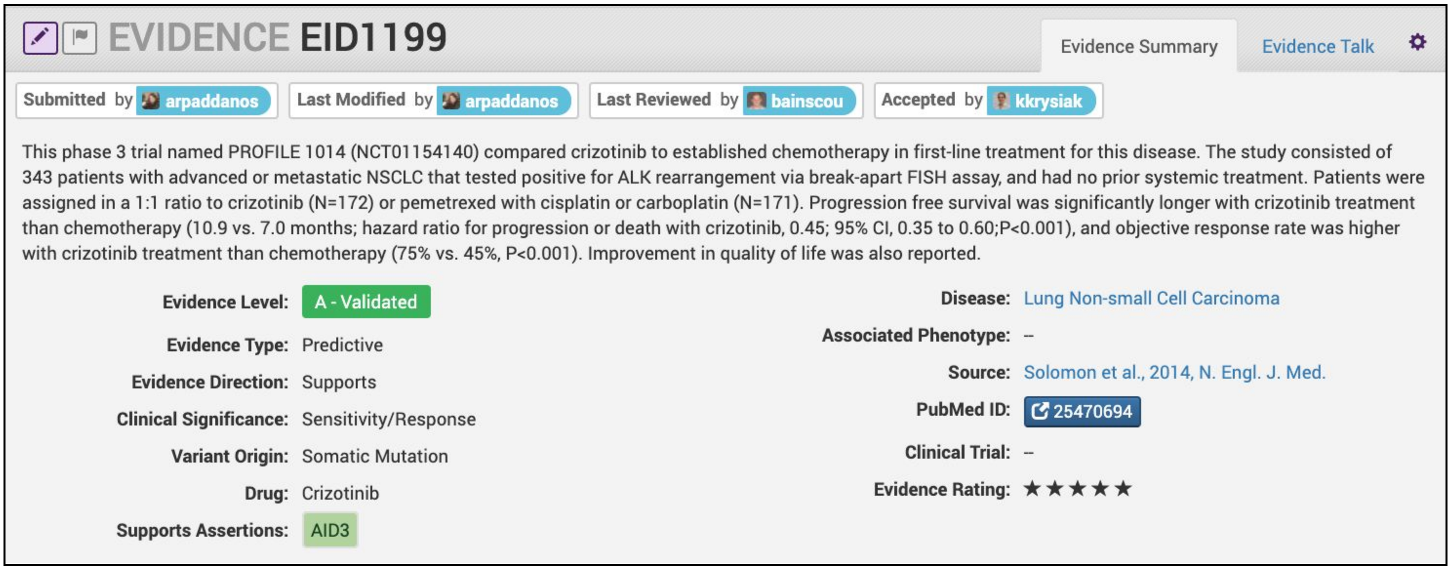


**Curation Practices:**

- Evidence Items with a 5-star rating should be strong, well-supported evidence from a lab or journal with respected academic standing. Experiments should be well controlled, and results should be clean and reproducible across multiple replicates. Evidence should be confirmed using independent methods and the study should be statistically powered.
- In general, Trust Rating is a rating of the unit of evidence extracted from a data source (publication or ASCO abstract) and is not a rating of the publication or abstract itself. Thus, an isolated supplementary figure in a high quality and well-researched publication may yield a relevant piece of clinical information on a variant of interest, and an Evidence Item (EID) could be prepared from this figure. Due to the limited nature of the data supporting this type of Evidence Item, it would receive a lower Trust Rating. This is because this rating would applies only to the evidence used to create this single EID, not to the publication as a whole.

**Supplementary Figure 25. Evidence Item with 4-star Trust Rating**

The example Evidence Item below describes a Phase 2A clinical trial with multiple arms that included evaluation of the therapeutic effect of pertuzumab and trastuzumab on 37 patients with *HER2* - Amplified colorectal cancer. The study was sufficiently powered to demonstrate an increase in patient response and remission provided their advanced, refractory state and relatively rare molecular alteration for that tumor type. The study was part of a clinical trial that was registered through the NIH and was published in the Journal of Clinical Oncology.

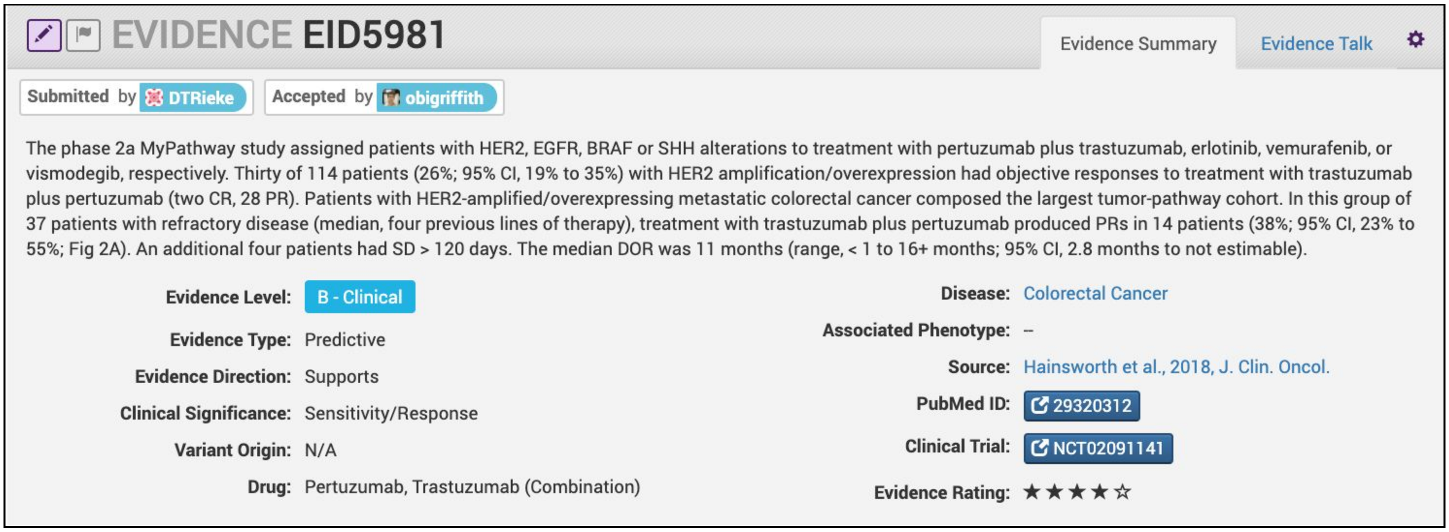


**Curation Practices:**

- Evidence items with a 4-star rating should be strong, have well-supported evidence, well-controlled experiments, and convincing results. Any discrepancies from expected results are well-explained and not concerning.
- This example was similar in design as the 5-start example, however, the reduced sample size contributed to the reduction in the start rating.

**Supplementary Figure 26. Evidence Item with 3-star Trust Rating**

The example Evidence Item below describes the same Phase 2A clinical trial from **Supplementary Figure 23**, but differs in the advanced solid tumor type being evaluated. In the subset of patients with bladder cancer, three of nine patients showed response to combination therapy with trastuzumab and pertuzumab, which supports sensitivity/response. Although this evidence item is derived from the same clinical trial, the reduction in Trust Rating is representative of the smaller number of patients and large 95% confidence interval.

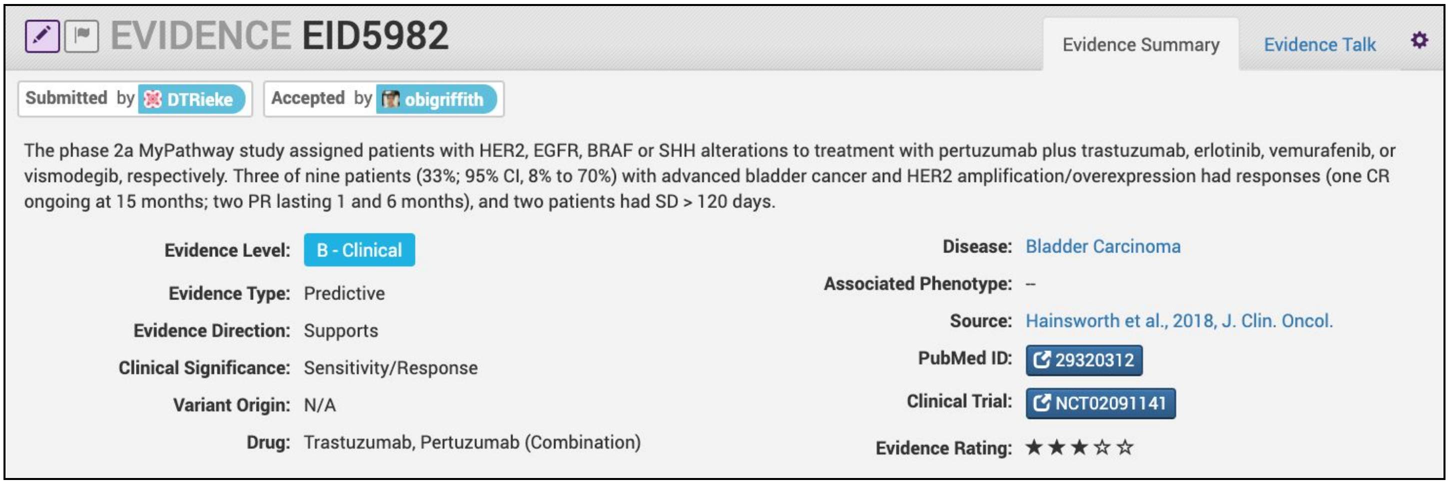


**Curation Practices:**

- Evidence Items with a 3-star rating are convincing but not supported by a breadth of experiments. These Evidence Items be smaller scale projects, or novel results without many follow-up experiments.
- Even though these Evidence Items might contain reduced amount of data discrepancies from expected results should still be explained and not concerning.

**Supplementary Figure 27. Evidence Item with 2-star Trust Rating**

The example Evidence Item below describes a Phase II clinical trial that evaluated 29 patients with breast cancer who were being treated with either afatinib, lapatinib, or trastuzumab. In this study, 18 patients showed response to one of the therapeutics. The Trust Rating for this evidence item was only 2 stars due to lack of evidence supporting the clinical claim (supports sensitivity/response). Specifically, the sample size was low for each of the three arms, there was no reported statistical significance. Additionally, the clinical endpoint for the study was objective response rate, which is not as strong of an endpoint as other metrics such as overall survival.

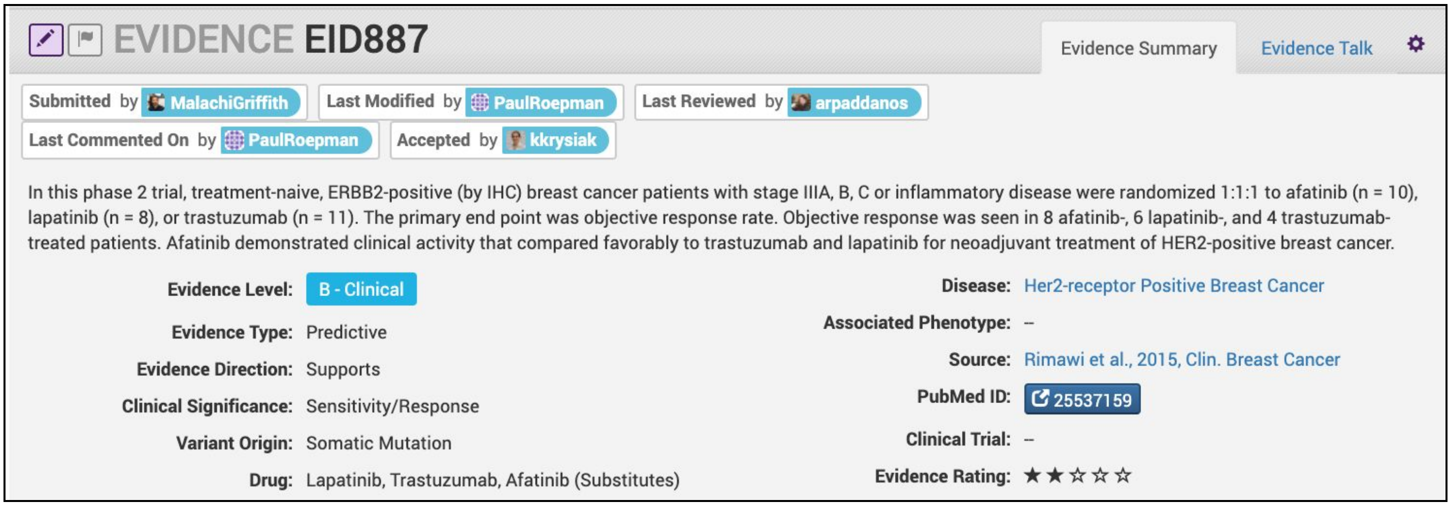


**Curation Practices:**

- Evidence items with a 2-star rating are not well supported by experimental data, and little follow-up data is available. These Evidence Items might be derived from a journal with low academic impact.
- Typically, Evidence Items received a 2-star rating if the experiments lack proper controls, have small sample size, or are not statistically convincing.

**Supplementary Figure 28. Evidence Item with 1-star Trust Rating**

The example Evidence Item below describes a B-level clinical study that evaluated 6 patients with *ERBB2* - Amplification for response to capecitabine, oxaliplatin, and chemoradiotherapy, with or without cetuximab. There was no difference in outcome between the 6 patients with the variant when compared to the 135 patients with no visible *ERBB2* - Amplification on FISH / IHC. The Evidence Item a heterogenous combination of variant detection methods, a low number of patients in the experimental arm (n=6) and overall low statistical power. Therefore, despite being a B-level Evidence Item, the curator assigned the EID a 1-star Trust Rating.

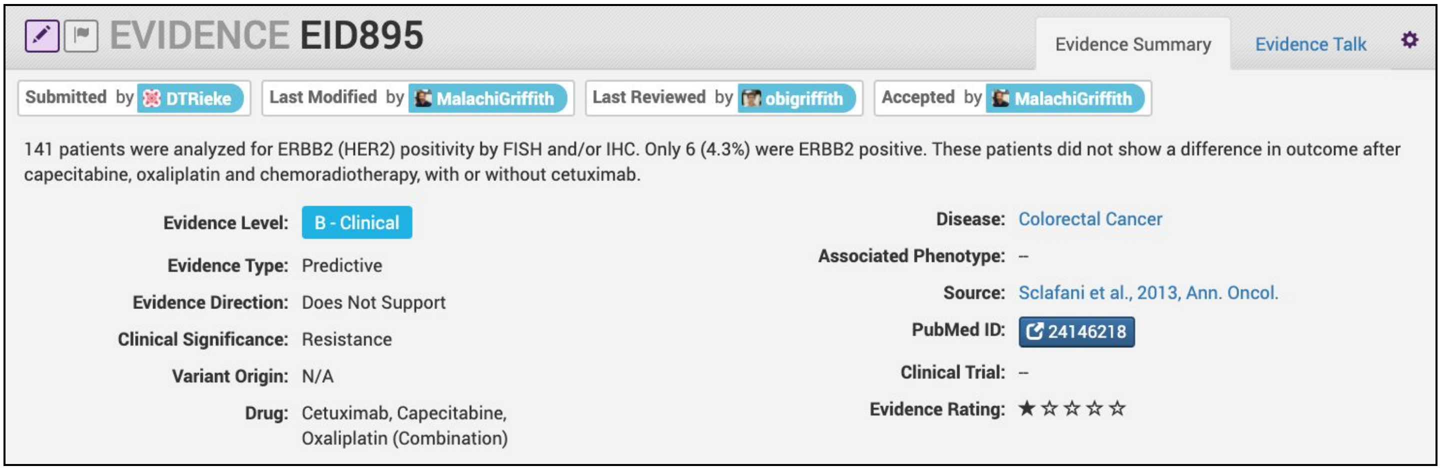


**Curation Practices:**

- Evidence items with a 1-star rating contain claims that are not well-supported by experimental evidence. Typically, the results are not reproducible and/or have very small sample size. No follow-up is done to validate novel claims.
- Typically, Evidence Items received a 1-star rating if the experiments lack proper controls, have small sample size, or are not statistically convincing.

### The Assertion data model

#### Examples of Assertion curation

**Supplementary Figure 29. Overview of CIViC Assertion entry form**

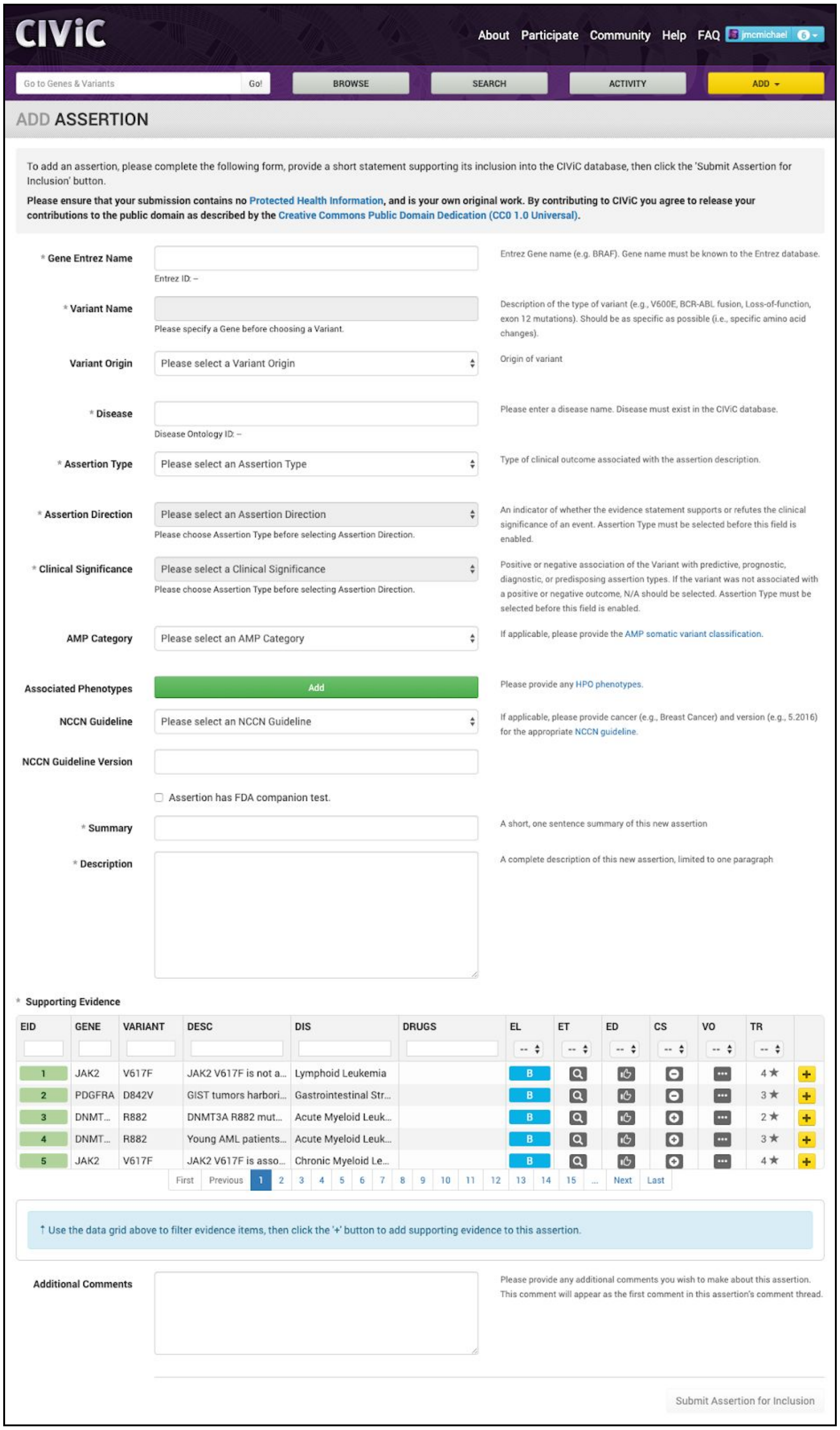


**Curation Practices:**

- The Supporting Evidence grid allows users to associate Evidence Items with Assertions. This can be accomplished by filtering on the provided fields (e.g., EID, Gene, Variant) and selecting the yellow plus sign on the right.
- A sufficient amount of evidence should be added to an Assertion so that the collection of Evidence Items represents that state of the field for the disease, variant, and type of Evidence Statement.
- As new evidence emerges for a variant, additional Evidence Items can be curated and added to the supporting evidence for an Assertion, potentially raising the ACMG classification or AMP Tier and Level.
- Practice guidelines, which are standard for the field from which the Assertion is derived, should be thoroughly consulted. Approved disease stage, and approved treatment lines should be outlined in the Assertion Description, if applicable.
- Approved companion diagnostics (e.g. Vysis Break-Apart Fish diagnostic for ALK-fusions) should be listed in the Assertion Description.

### Examples of Variant Origin

##### Supplementary Figure 30. Example of curation of Variant Origin

This is an example of an Assertion whereby the Variant Origin reflects the Evidence Items supporting the Assertion. In this example, the *VHL* - Q195* (c.583C>T) variant is implicated as Pathogenic based on ACMG Codes PVS1, PM2, and PP4. All Evidence Items supporting this Assertion are Case Studies from families that show how this specific germline variant caused the resulting disease.

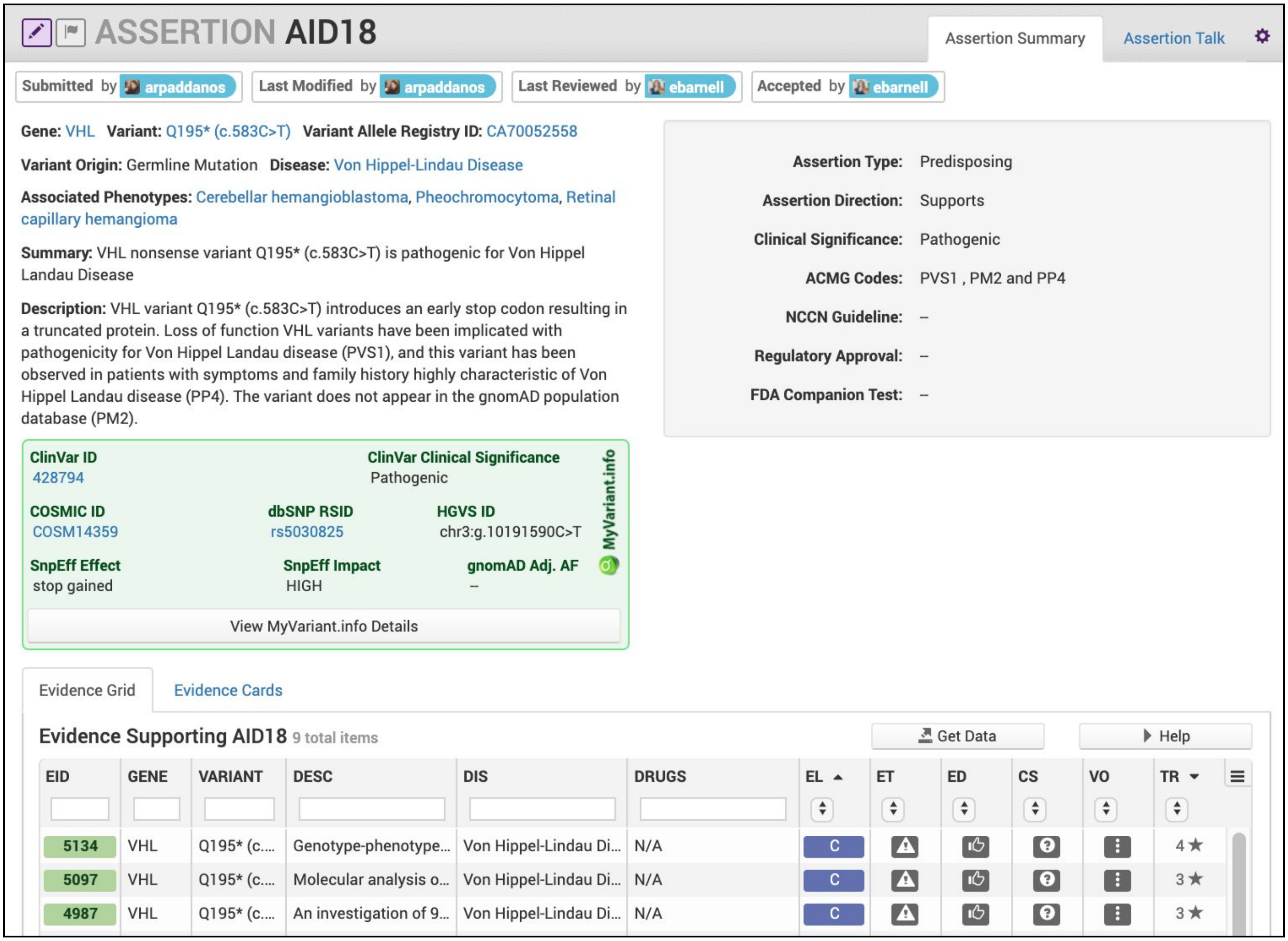


**Curation Practices:**

- Typically, Germline Mutations or Germline Polymorphisms are associated with Predisposing Assertions, however, this is not always the case.
- Clicking on the Evidence Cards tab of the Evidence Grid will display all underlying Evidence Items in their entirety.

### Examples of Disease

**Supplementary Figure 31. Selection of Disease Type for Assertions**

Only one Disease is permitted for each Assertion. It is recommended that the Disease be as specific as possible while still holding true for all Evidence Items associated with the Assertion. The example Assertion below is composed of Evidence Items whose Disease is either "Melanoma" or "Skin Melanoma". At the Assertion level, the less specific Disease type (Melanoma) is selected to represent all supporting Evidence Items.

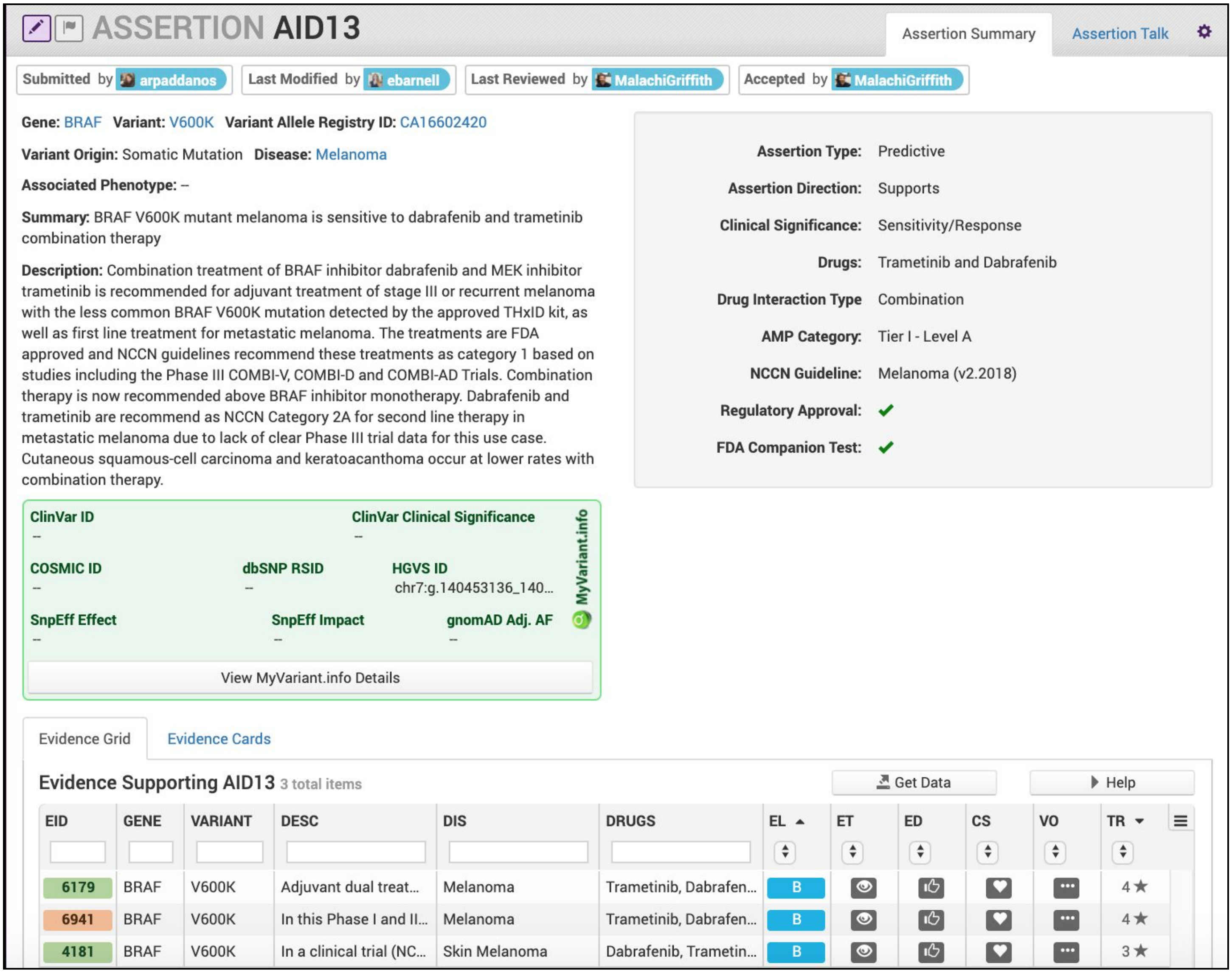


**Curation Practices:**

- Disease stage to which the assertion applies, as well as the line of treatment (e.g. first line, salvage, etc) should be made explicit in the Assertion Description. While the body of supporting Evidence Items may be derived from studies with differing patient populations with regard to stage and line of treatment, as well as preclinical studies in disease models, practice guidelines (e.g. NCCN etc) should be consulted for approved use cases.

### Examples of Assertions

##### Supplementary Figure 32. Exemplary Predictive Assertions

The figure below shows a Predictive Assertion with an exemplary Assertion Summary and Assertion Description. In this example, the Assertion describes that *BRAF* V600E confers sensitivity to combination therapy of dabrafenib and trametinib for patients with melanoma. The AMP Category is Tier I - Level A for this variant, disease and treatment, and is an association listed in the Melanoma NCCN Guidelines (v2.2018).

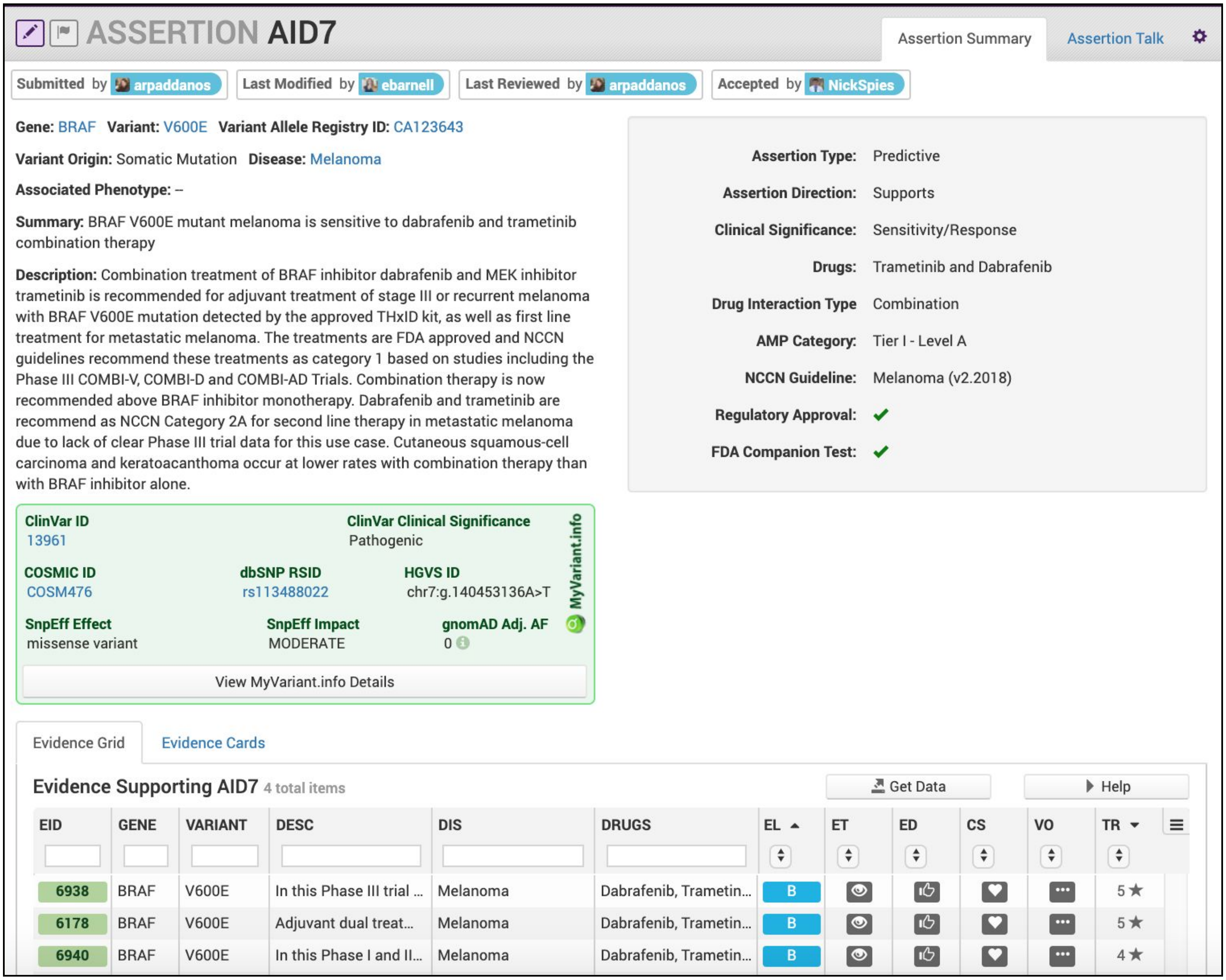


**Curation Practices:**

- All Evidence Items relevant to the Assertion should associated, even if they disagree with the Assertion Summary. Disagreements can be discussed in the Description section and rationale for discounting discrepant evidence should be recounted.
- AMP Level and Tier should be associated with each Predictive, Diagnostic and Prognostic Assertions [7]. For methods on assigning AMP Tier / Level, See **Figure 5A**.

**Supplementary Figure 33. Exemplary Prognostic Assertions**

The figure below shows a Prognostic Assertion with an exemplary Assertion Summary and Assertion Description. In this example, the Assertion describes that the *BRAF* V600E variant confers poor outcome for patients with colorectal cancer. This variant has an associated FDA companion diagnostic test, is listed in the NCCN Guidelines for colorectal cancer (v2.2017), and falls under the Tier I - Level A AMP category.

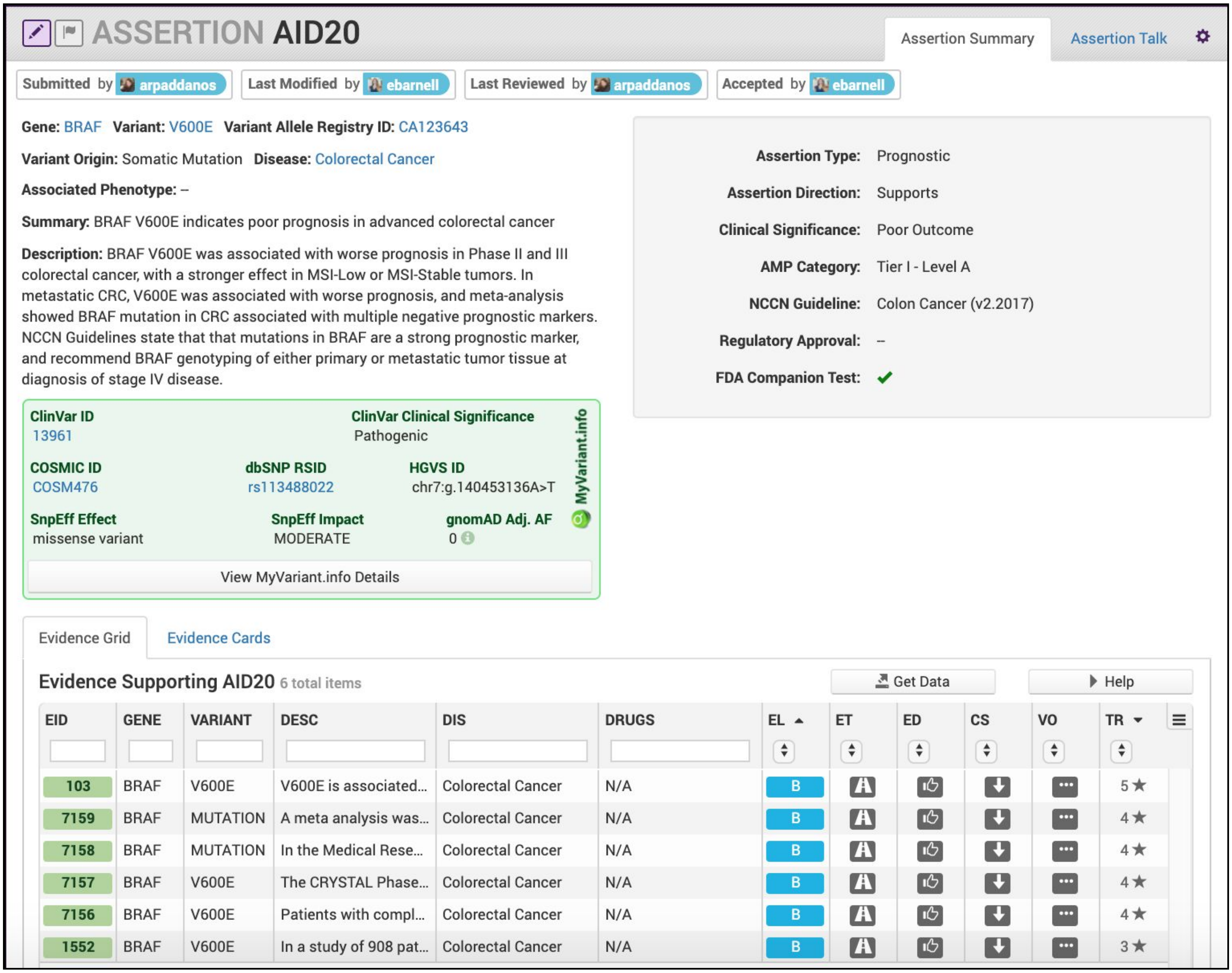


**Curation Practices:**

- All Evidence Items relevant to the Assertion should associated, even if they disagree with the Assertion Summary. Disagreements can be discussed in the Description section and rationale for discounting discrepant evidence should be recounted.
- Prognostic evidence in CIViC demonstrates variant association with better or worse patient outcome in a general manner, that is independent of any specific treatment context. Therefore, a larger collection of evidence showing similar prognostic outcomes under a range of different treatment or untreated regimes is ideal.
- Application of AMP Tier and Level [7] is dependant on practice guidelines (e.g. NCCN) ascribing prognostic value to the variant for the given disease, or failing this, the quality and level of evidence supporting the Assertion (**Figure 5A**).

**Supplementary Figure 34. Exemplary Diagnostic Assertions**

Below is an example of a Diagnostic Assertion with an exemplary Assertion Summary and Assertion Description. In this example, the Assertion describes how an in-frame fusion between *DNAJB1* and *PRKACA* can be used to diagnose a specific subtype of hepatocellular carcinoma (HCC). Presence of this fusion can be used to clarify that the patient has fibrolamellar HCC.

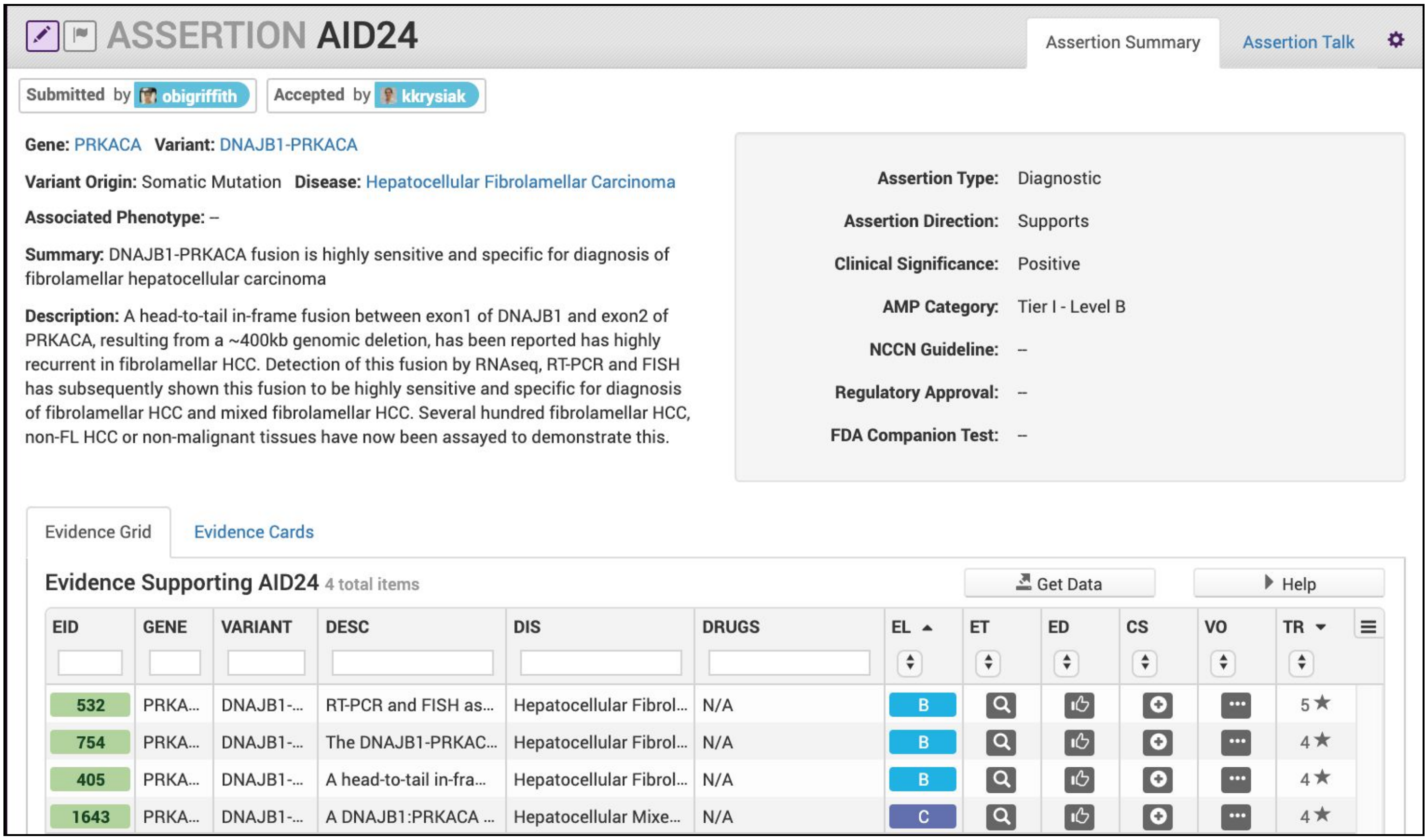


**Curation Practices:**

- All Evidence Items relevant to the Assertion should associated, even if they disagree with the Assertion Summary. Disagreements can be discussed in the Description section and rationale for discounting discrepant evidence should be recounted.
- The evidence supporting the Assertion should sufficiently cover what is known regarding the diagnostic power for the variant in the specific disease context.
- For Tier I Level A Diagnostic Assertions, details from relevant practice guidelines should be given, along with any additional specific information which is applicable (e.g., disease stage).
- Lower Tier and Evidence Level Assertions may be created for diagnostic variants not currently in practice guidelines. Variants backed by stronger clinical data may be Tier I Level B as above. Variants with smaller amounts of evidence for diagnostic potential will receive lower Tiers and Evidence Levels (**Figure 5A**).

**Supplementary Figure 35. Exemplary Predisposing Assertion**

Below is an example of a Predisposing Assertion. In this example, an inframe deletion repeatedly observed in the literature is considered pathogenic for Von Hippel-Lindau Disease. Utilizing the ACMG guidelines [8], evidence codes were assembled from the literature (PP1, PS2) and variant-level information (PM4, PM2) to be categorized as Pathogenic. Specific evidence is associated with codes in the Description and all evidence evaluated when producing the Assertion is associated with the Assertion.

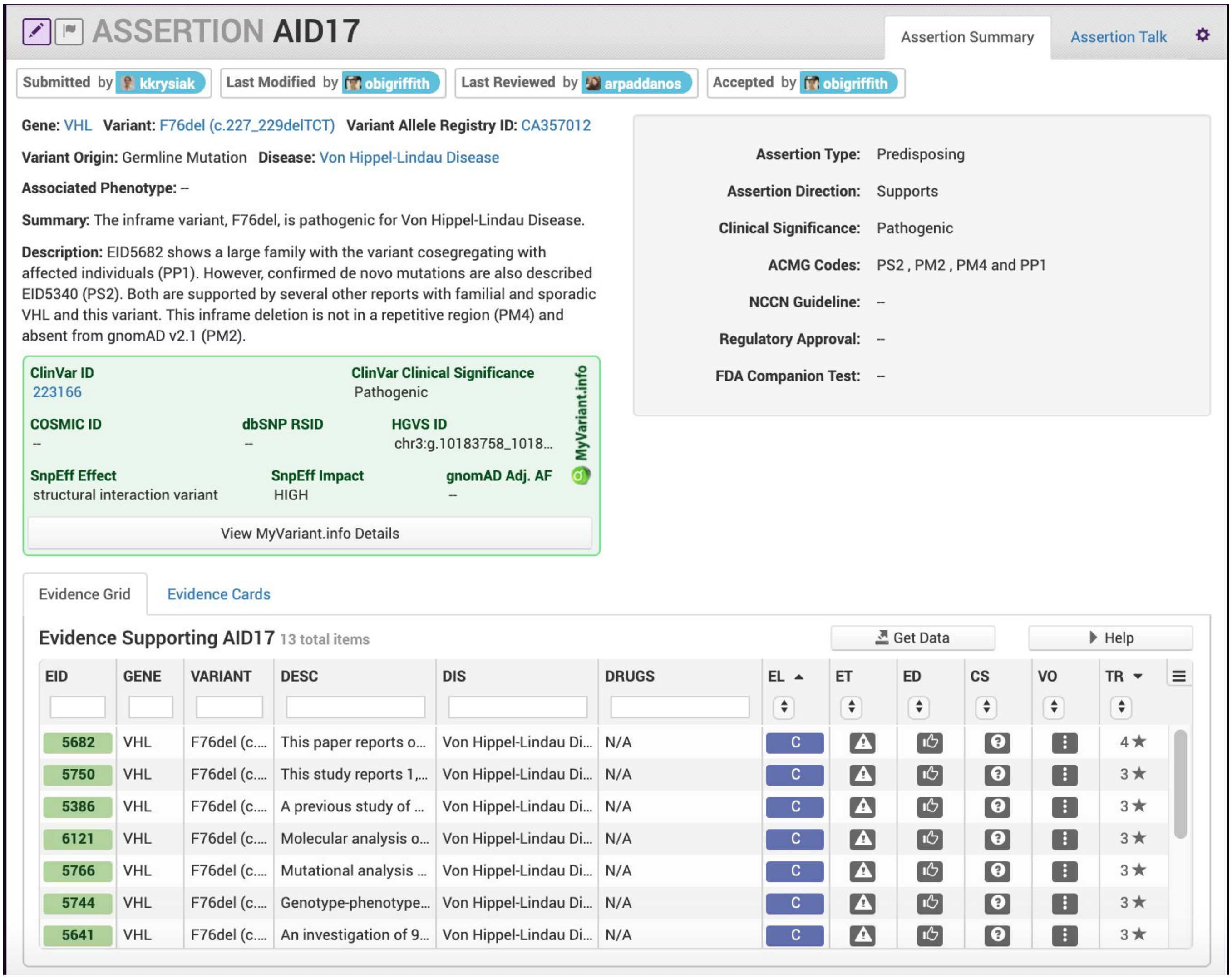


**Curation Practices:**

- ACMG codes [8] supporting the Predisposing Assertion are derived from supporting Evidence Items, and other sources such as population databases (See **Figure 5B**). Any evidence codes applied should be explained in the Description section, allowing others to rapidly re-evaluate the evidence used.
- All Evidence Items relevant to the Assertion should associated, even if they disagree with the Assertion Summary. Disagreements can be discussed in the Description section and rationale for discounting discrepant evidence should be recounted.
- Thoroughly evaluated Assertions can have a clinical significance of Variant of Unknown Significance using ACMG criteria. This permits other users to quickly re-evaluate this variant in the context of new evidence, potentially leading to reclassification, but reducing future curation burden if the variant is observed again.

### Examples of Assertion Supporting Evidence

**Supplementary Figure 36. Requirements for an Assertion to be accepted**

A complete list of Assertions can be found on the Assertions tab of the Browse page.

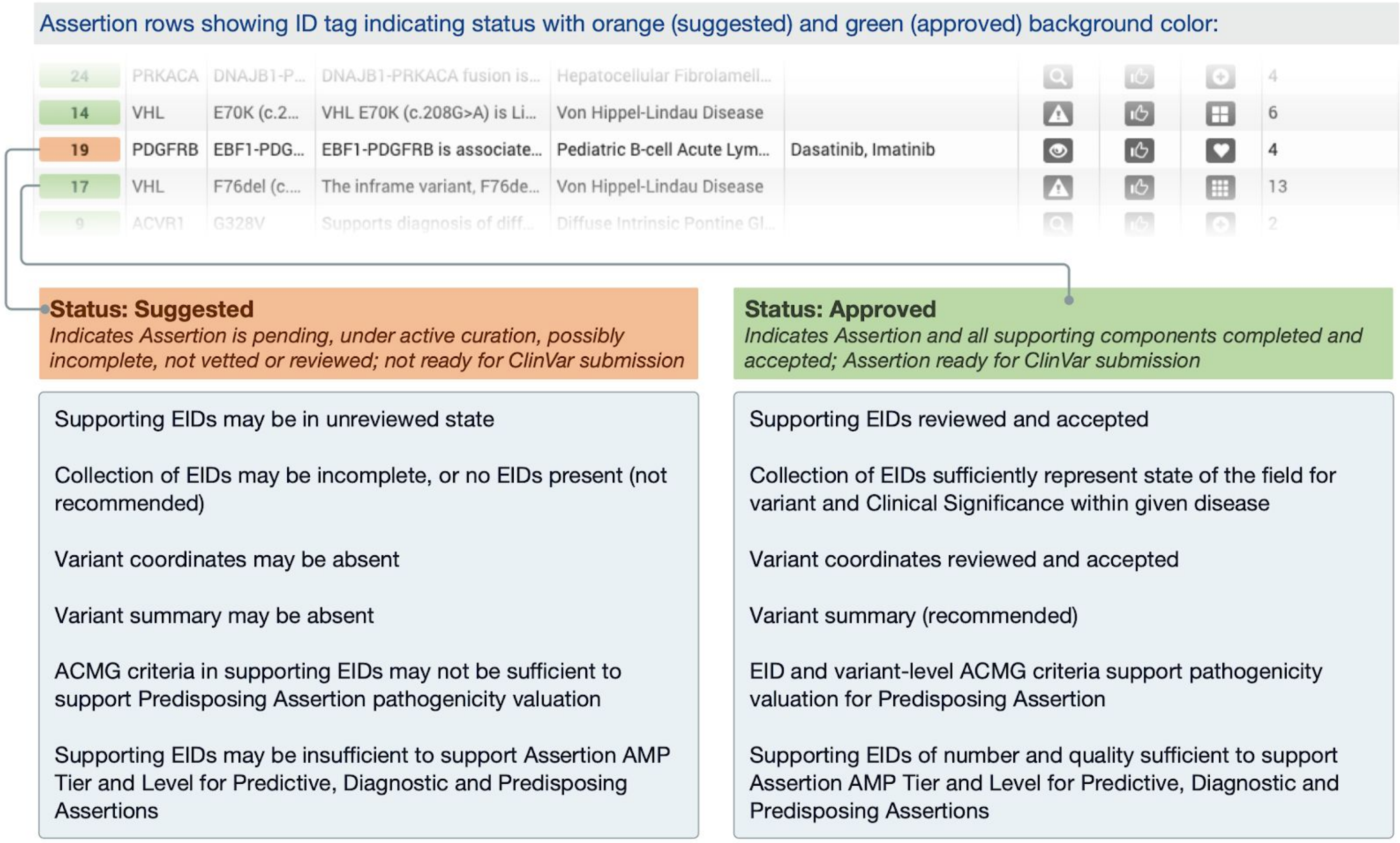


**Curation Practices:**

- Assertions can be curated without being associated with any Evidence Items; however, they cannot be accepted until at least one Evidence Item is linked to the Assertion.
- A new Assertion can only be created for a variant/gene that already exists in CIViC.
- Evidence Items that are associated with Assertions must be accepted prior to the Assertion being accepted; however, these Evidence Items may still be revised and edited after the Assertion has been accepted.

### Supplementary Tables

**Supplementary Table 1. General curation notation for all items within CIViC**

| Button/Icon | Name | Description |
|---|---|---|
| 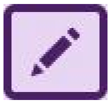 | Edit Button | Selection of the purple pencil in the upper left corner allows curators to edit an entity. |
| 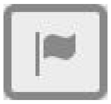 | Flag Button | Clicking the flag in the upper left corner allows a curator to flag an entity for additional review. |
| 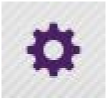 | Preferences Button | Users can subscribe to curation updates for a specific entity by clicking the gear icon in the top right of the screen. |
| 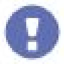 | Pending Revisions Icon | Hovering over the exclamation mark icon will show pending revisions for the entity. |

#### Supplementary Table 2. Examples of Variants supported by the CIViC interface

| Variant name | Variant type (Sequence Ontology) | Demonstrates support of… |
|---|---|---|
| DPYD*2A Homozygosity | Splice Donor Variant | pharmacogenomic nomenclature |
| ALK Fusions | Transcript Fusion | fusions with an unknown partner common in FISH techniques |
| EML4-ALK | Transcript Fusion | specific gene fusions |
| EML4-ALK E6;A20 | Transcript Fusion | fusions with known specific exon boundaries / specific fusion isoforms |
| BCR-ABL T315I | Missense Variant | specific variants in the context of other variants |
| CDKN2A Promoter Hypermethylation | N/A | epigenetic modifications |
| p16 Expression | N/A | expression changes |
| rs3814960 | 5 Prime UTR Exon Variant | polymorphisms |
| FLT3-ITD | Inframe Insertion | imprecise insertions with shared consequences |
| KIT Exon 11 Mutation | Coding Sequence Variant | categorical variants covering specific transcriptional boundaries |
| TP53 DNA Binding Domain Mutation | DNA Binding Site | categorical variants covering specific functional boundaries |
| MET Exon 14 Skipping Mutation | Exon Loss Variant | categorical variants covering specific transcriptional consequences |
| BRCA Loss-of-function | Loss of Function Variant | categorical variants covering specific functional consequences |
| BRAF V600E | Protein Altering Variant | a specific amino acid change which includes all possible DNA changes which result in this missense change |
| BRAF V600 | Protein Altering Variant | categorical variants involving a single amino acid |
| BRAF non-V600 | Protein Altering Variant | categorical variants excluding a common hotspot |
| VHL S65W (c.194C>G) | Missense Variant | precise base pair resolution variants |
| ERBB2 SERUM LEVELS | N/A | sample-specific protein levels measured by assays such as ELISA |
| IGF1R NUCLEAR EXPRESSION | N/A | cellular compartment-specific expression measured by assays such as immunohistochemistry (IHC) |
| STAT3 SH2 Domain Mutation | Coding Sequence Variant | variants within a specific domain |

#### Supplementary Table 3. Examples of classification Sequence Ontology Term classifications

| Sequence Ontology Term | Sequence Ontology Definition | Examples |
|---|---|---|
| missense_variant | A sequence variant, that changes one or more bases, resulting in a different amino acid sequence but where the length is preserved. | G12D |
| stop_gained | A sequence variant whereby at least one base of a codon is changed, resulting in a premature stop codon, leading to a shortened polypeptide. | R130* |
| protein_altering_variant | A sequence_variant which is predicted to change the protein encoded in the coding sequence. | G12<br>KINASE DOMAIN MUTATION |
| frameshift_truncation | A frameshift variant that causes the translational reading frame to be shortened relative to the reference feature. | v2288fs*1 |
| inframe_deletion | An inframe non synonymous variant that deletes bases from the coding sequence. | DEL I843<br>V560DEL<br>DEL755-759 |
| inframe_insertion | An inframe non synonymous variant that inserts bases into in the coding sequence. | P780INS<br>M77INSAYVM<br>ITD |
| [gene_variant<br><br>OR transcript_variant]<br><br>AND<br><br>[loss_of_function_variant<br><br>OR<br><br>gain_of_function_variant] | [ gene_variant: A sequence variant where the structure of the gene is changed.<br>OR<br>transcript_variant: A sequence variant that changes the structure of the transcript ]<br><br>AND<br><br>[ loss_of_function_variant: A sequence variant whereby the gene product has diminished or abolished function.<br><br>OR<br><br>gain_of_function_variant: A sequence variant whereby new or enhanced function is conferred on the gene product. ] | MUTATION |
| exon_variant | A sequence variant that changes exon sequence. | EXON 10 MUTATION |
| transcript_fusion<br><br>OR RARELY<br><br>gene_fusion | transcript_fusion: A feature fusion where the deletion brings together transcript regions.<br><br>OR RARELY<br><br>gene_fusion: A sequence variant whereby a two genes have become joined. | EML4-ALK<br>ALK FUSIONS |
| transcript_fusion<br><br>AND<br><br>missense_variant | transcript_fusion: A feature fusion where the deletion brings together transcript regions.<br><br>AND<br><br>missense_variant: A sequence variant that changes one or more bases, resulting in a different amino acid sequence but where the length is preserved. | ELM4-ALK G1269A |

| | | |
|---|---|---|
| transcript_translocation<br><br>OR<br><br>feature_translocation<br><br>OR<br><br>transcript_fusion | transcript_translocation: A feature translocation where the region contains a transcript.<br><br>OR<br><br>feature_translocation: A sequence variant, caused by an alteration of the genomic sequence, where the structural change, a translocation, is greater than the extent of the underlying genomic features.<br><br>OR<br><br>transcript_fusion: A feature fusion where the deletion brings together transcript regions. | REARRANGEMENT |
| wild_type | An attribute describing sequence with the genotype found in nature and/or standard laboratory stock. | WILD TYPE |
| loss_of_heterozygosity | A functional variant whereby the sequence alteration causes a loss of function of one allele of a gene. | LOH |
| transcript_amplification | A feature amplification of a region containing a transcript. | AMPLIFICATION |
| transcript_ablation | A feature ablation whereby the deleted region includes a transcript feature. | DELETION |
| copy_number_change | A sequence variant where copies of a feature (CNV) are either increased or decreased. | COPY NUMBER VARIATION |
| loss_of_function_variant | A sequence variant whereby the gene product has diminished or abolished function. | LOSS-OF-FUNCTION |
| Loss_of_function_variant OR transcript_ablation | loss_of_fuction_variant: A sequence variant whereby the gene product has diminished or abolished function.<br><br>transcript_ablation: A feature ablation whereby the deleted region includes a transcript feature. | LOSS |
| exon_loss_variant | A sequence variant whereby an exon is lost from the transcript. | EXON 14 SKIPPING MUTATION |
| 5_prime_UTR_variant | A UTR variant of the 5' UTR. | 5' UTR MUTATION |
| 3_prime_UTR_variant | A UTR variant of the 3' UTR. | 3' UTR MUTATION |
| N/A | | EXPRESSION<br>NUCLEAR EXPRESSION<br>CYTOPLASMIC EXPRESSION<br>OVEREXPRESSION<br>UNDEREXPRESSION<br>METHYLATION<br>PROMOTER METHYLATION<br>PROMOTER HYPERMETHYLATION |

**Supplementary Table 4. Use cases for all combinations of Evidence Types, Direction, and Clinical Significance for Predictive Evidence**

| Predictive Evidence Items | | | | | |
|---|---|---|---|---|---|
| **Context** | **Direction** | **Preclinical (Level D)** | **Case Study (Level C)** | **Clinical (Level B)** | **Comment** |
| Clinical or preclinical studies testing for the ability of a variant to induce sensitivity to a given treatment in a specific disease context, where patient populations or preclinical samples without the variant may be used as controls. | Predictive: Supports Sensitivity or Response | Controlled experiments in preclinical models demonstrating sensitization of variant in comparison to wildtype. | Clinical observation of a patient suggesting variant association with a treatment response, where treatment response is not observed / expected for wildtype | Phase I, II, III or other clinical study demonstrating statistical variant association with sensitivity, possibly compared to wildtype control population. | Associates a given variant and treatment combination with sensitivity. If work described in the evidence item has been directly cited to support regulatory approval or practice guideline, then evidence item is labeled as Validated Level A. |
| | Predictive: Does Not Support Sensitivity or Response | Controlled experiments in preclinical models demonstrating no sensitization of variant in comparison to wildtype. | Clinical observation of a patient with variant responding to treatment as would a wildtype patient. | Phase I, II, III or other clinical study with failure to show statistical benefit associated with the variant, possibly compared to wildtype control population. | Variant behaves the same with treatment as non-sensitized wildtype. |
| Clinical or preclinical studies testing for the ability of a variant to induce resistance to a particular treatment in a specific disease context, where the background (without variant) is sensitive to the treatment. | Predictive: Supports Resistance | Controlled experiments for particular treatment in preclinical models demonstrating abolished sensitization with variant in comparison to sensitization in the absence of variant. | Clinical observation of patient suggesting variant association with a treatment resistance, where response would otherwise have been expected. Evidence receives higher star rating if patient history shows previous sensitivity to treatment, with variant appearing in testing after, but not in testing performed before the resistance occurred. | Phase I or higher phase study demonstrating statistical variant association with resistance to a given treatment. | Introduction of variant abolishes sensitivity observed in the background state without variant. If work described in the evidence item has been directly cited to support regulatory approval or practice guideline, then evidence item is labeled as Validated Level A. |
| | Predictive: Does Not Support Resistance | Controlled experiments for particular treatment in preclinical models demonstrating no abolished sensitization with variant in comparison to sensitization in the absence of the variant. | Observation of a patient with a given variant responding to treatment in the same way as a sensitized patient would be expected to. | Phase I or higher phase study demonstrating no statistical association with the variant and resistance to a given treatment. | This annotation is not equivalent to “Supports Sensitivity”, since the variant is not sensitizing, but instead is neutral, and does not affect the baseline sensitivity. The annotation is best used when guidelines suggest variants of the given type may induce resistance, such as *KRAS* mutations in colorectal cancer with respect to EGFR inhibitor treatment. |

| | | | | | |
|---|---|---|---|---|---|
| Comparison of a variant to a different variant known to be sensitive to a given treatment for a specific disease. | Predictive: Supports Reduced Sensitivity | Controlled experiments in preclinical models demonstrating lesser degree of sensitization of the variant in comparison to an established sensitizing variant, but increased sensitization over wildtype. | Supports Sensitivity/Response best used in this context | Clinical study showing statistically significant intermediate response for given variant between established sensitized variant and baseline wildtype gene. | In case studies (Level C Evidence) this Clinical Significance is not recommended due to lack of adequate numbers of controls for comparison. Note that Reduced Sensitivity is used to compare two variants, but not two treatment regimes. |
| | Predictive: Does Not Support Reduced Sensitivity | Supports Sensitivity/Response recommended in this context | Supports Sensitivity/Response recommended in this context | Supports Sensitivity/Response recommended in this context | Does Not Support Reduced Sensitivity is not currently a recommended annotation, as no clinically relevant use case for the annotation is apparent. |
| Comparison of two different treatment types against the same variant, within the same disease context. | Predictive: Supports Sensitivity | (Noninferiority) Controlled experiments in preclinical models demonstrating a variant responds equally well to a new treatment in comparison to established treatment for the variant. | NA | (Noninferiority) Clinical studies showing patients do not fare worse with a new variant-targeted treatment in comparison to established treatment for the given variant. This evidence supports sensitivity for the variant to the new treatment. | Case studies will not support this type of evidence since a single or small group of patients cannot generate the necessary statistical power for comparison. If work described in the evidence item has been directly cited in support of regulatory approval or practice guidelines, then the evidence item is labeled Validated Level A. |
| | Predictive: Does Not Support Sensitivity | Controlled experiments in preclinical models demonstrate lesser degree of sensitization of variant to a new treatment in comparison to established treatment for the variant. | NA | Clinical trials demonstrating lesser degree of sensitization of variant to a new treatment in comparison to established treatment for the variant. In this case, the evidence does not support sensitivity for the variant to the new treatment. | When curating Evidence Items from results which show a newer treatment is less effective than an existing treatment for a given variant, then the Predictive, Does Not Support Sensitivity annotation may be used for the variant and disease with respect to the new treatment. The comparison to existing treatment can be described in the evidence summary. |
| Observation of patient samples or preclinical systems with association to a specific disease type, which are positive for the given variant. Comparison to various control samples, may be performed. | Diagnostic: Supports Positive | Preclinical work suggesting association between variant and disease or disease subtype. | Observations of variant being present in a small number of patients with the given disease, and potential comparison to patients without variant or disease | Clinical observations in a patient population of significant variant association with a positive diagnosis of a given disease. | Evidence from publications describing practice guidelines or regulatory approval may be used for curation and labeled as Validated level A. Work suggesting novel disease classification based on or related to the given variant may be curated by experts in the field, but is not recommended for general curation. Submission of |

| | | | | | |
|---|---|---|---|---|---|
| Publications citing guidelines also can be used. | | | | | new terms to Disease Ontology should accompany such curation. |
| | Diagnostic: Does Not Support Positive | Preclinical work suggesting no association between variant and disease or disease subtype. | Observations demonstrating lack of variant in small number of patients with specified disease or variant presence in patients without the specified disease. | Clinical studies with statistical results showing variant cannot support positive diagnosis. | This annotation will generally be used in cases where the variant could be expected to have diagnostic significance (e.g. previous findings), so that reports to the contrary could hold clinical interest. |
| Observation of patient samples or preclinical systems with contraindication to a specific disease type, which are positive for the given variant. Comparison to various control samples may be performed. | Diagnostic: Supports Negative | Preclinical work suggesting variant associated with negative diagnosis. | Smaller numbers of patient observations suggest variant may be a contraindication for a specific disease. | Studies with statistically significant findings which suggest that variant may be added to exclusion criteria for a given disease. | See comments for Diagnostic: Supports Positive |
| | Diagnostic: Does Not Support Negative | Preclinical work suggesting lack of association between variant and negative diagnosis. | Smaller numbers of patient observations suggest variant is not a contraindication for a specific disease. | Studies with statistically significant findings suggesting that the variant has no diagnostic power for the given disease. | See comments for Diagnostic: Does Not Support Positive |
| Variant present in patient populations or preclinical samples directly associated with better outcome, or associated with known markers of better outcome. | Prognostic: Supports Better Outcome | Experiments in preclinical systems suggesting variant is associated with better outcome, for instance through demonstration of association with cellular markers of less aggressive disease. | Case study reports or smaller trial subgroups, where patients with the given variant show good outcomes with respect to a given clinical measure, or show other indications associated with better outcome (e.g. patient samples show markers of better outcome). | Clinical observation that patient subgroups with the variant have better outcomes than patients without the variant, and that this result is not specific to a particular treatment type. | If work described in the evidence has been directly cited to support regulatory approval or practice guidelines, then the Evidence Item is labeled Validated Level A. Prognostic evidence refers to better or worse outcome associated with a specific variant and disease, which is shown to occur regardless of specific treatment context. |
| | Prognostic: Does Not Support Better Outcome | Controlled preclinical experiments showing variant lacks association with better outcome. | Case study reports or trial subgroups with small numbers of patients with the given variant, that do not suggest a better outcome. | Clinical patient data which show no significant association of variant with better outcome in comparison to patients without variant. | This annotation will generally be used in cases where the variant could be expected to have prognostic significance (e.g. previous findings), so that reports to the contrary could hold clinical interest. |
| Variant present in patient populations or preclinical samples directly associated with poor outcome, or associated with known markers of poor outcome. | Prognostic: Supports Poor Outcome | Preclinical work suggesting association between variant and indicators of poor prognosis such as proliferative biomarkers, (e.g. Ki-67). | Case study reports or trial subgroups with small numbers of patients with the given variant, which suggest a poor outcome. Patient samples may show markers associated with a worse outcome. | Clinical observation that patient subgroups with the variant have significantly worse outcomes by some clinical measure, not specific to a particular treatment context. | See comments for Prognostic: Supports Better Outcome |

| | | | | | |
|---|---|---|---|---|---|
| | Prognostic Does Not Support Poor Outcome | Preclinical work that suggests lack of association with variant and indicators of poor prognosis such as proliferative biomarkers. | Case study reports or trial subgroups with smaller numbers of patients with the given variant that do not suggest a poor outcome. | Clinical patient data which show no significant association of variant with poor outcome, in comparison to patients without variant. | See comments for Prognostic Does Not Support Better Outcome |

**Supplementary Table 5. Definitions of Clinical Significance for all Evidence Types**

| Evidence Type | Clinical Significance | Definition / Curation Tips |
|---|---|---|
| Predictive *(Impact on therapeutic response )* | Sensitivity/Response | Associated with a clinical or preclinical response to treatment |
| | Resistance | Associated with clinical or preclinical resistance to treatment |
| | Adverse Response | Associated with an adverse response to drug treatment |
| | Reduced Sensitivity | Response to treatment is lower than seen in other treatment contexts |
| | N/A | Variant does not inform clinical action |
| Diagnostic *(Impact on diagnosis or disease subtype)* | Positive | Associated with diagnosis of disease or disease subtype |
| | Negative | Associated with lack of disease or disease subtype |
| Prognostic *(Impact on disease progression or patient survival )* | Better Outcome | Demonstrates better than expected clinical outcome |
| | Poor Outcome | Demonstrates worse than expected clinical outcome |
| | N/A | Variant does not inform clinical action |
| Predisposing *(Impact on disease susceptibility )* | Pathogenic | Very strong evidence the variant is pathogenic |
| | Likely Pathogenic | Strong evidence (>90% certainty) the variant is pathogenic |
| | Benign | Very strong evidence the variant is benign |
| | Likely Benign | Not expected (>90% certainty) to have a major contribution to the disease |
| | Uncertain Significance | Does not fulfill the ACMG criteria for pathogenic, likely pathogenic, likely benign or benign |
| Functional *(Impact on biological alterations)* | Gain of Function | A sequence variant whereby enhanced function is conferred on the gene product |
| | Loss of Function | A sequence variant whereby the gene product has diminished or abolished function |
| | Unaltered Function | A sequence variant whereby the function of the gene product is unchanged |
| | Neomorphic | A sequence variant whereby the gene product creates a novel function |
| | Unknown | A functional variant that cannot be precisely defined by gain-of-function, loss-of-function, neomorphic or unaltered function |

**Supplementary Table 6. General guidelines and examples for trust ratings**

To quickly discern how much trust curators and users have in a single Evidence Statement, a five-star trust rating system is used. Each Evidence Item is given a rating, from 1 to 5 stars, based on the quality of the evidence the statement summarizes. This rating depends on a number of factors, including journal impact, study size, quality control, orthogonal validation, and reproducibility. It should be noted that this rating is largely subjective and may be debated (ideally within the CIViC interface). The rating is specific to the data and conclusions within the Evidence Statement. The overall publication/study might be high quality in a high impact publication, but the Evidence Statement may refer to a single conclusion in the study, and that part of the study might not be well supported. For example, the assertion may relate to patients with a particular mutation, and the study might involve an impressive 500 patients, but if only 2 patients have the mutation in question, the quality rating may be low for this Evidence Statement.

| | |
|---|---|
| ★★★★★ | Strong, well supported evidence from a lab or journal with respected academic standing. Experiments are well controlled, and results are clean and reproducible across multiple replicates. Evidence confirmed using independent methods. The study is statistically well powered. |
| ★★★★ | Strong, well supported evidence. Experiments are well controlled, and results are convincing. Any discrepancies from expected results are well-explained and not concerning. |
| ★★★ | Evidence is convincing, but not supported by a breadth of experiments. May be smaller scale projects, or novel results without many follow-up experiments. Discrepancies from expected results are explained and not concerning. |
| ★★ | Evidence is not well supported by experimental data, and little follow-up data is available. Publication is from a journal with low academic impact. Experiments may lack proper controls, have small sample size, or are not statistically convincing. |
| ★ | Claim is not supported well by experimental evidence. Results are not reproducible, or have very small sample size. No follow-up is done to validate novel claims. |

**Supplementary Table 7. Markdown and Macros.**

Markdown is used to add emphasis, styling, images, and links to comments. Macros add links to specific CIViC users and entities. Below show the methods for employing Markdown and Macros to fields within the CIViC interface. These are particularly intended for comments on CIViC entities.

| **Markdown** | |
|---|---|
| **Headers** | # This is an <h1> tag |
| | ## This is an <h2> tag |
| | ###### This is an <h6> tag |
| **Emphasis** | *This text will be italic*<br>_This will also be italic_ |
| | **This text will be bold**<br>__This will also be bold__ |
| | *Italics and boldface **can be** combined* |
| **Unordered List** | * Item 1<br>* Item 2<br>* Item 2a<br>* Item 2b<br>* Item 3 |
| **Ordered Lists** | 1. Item 1<br>2. Item 2<br>3. Item 3<br>* Item 3a<br>* Item 3b |
| **Images** | `![this cool diagram](http://site.com/images/cool.png)`<br>Format: `![Alt Text](url)` |
| **Links** | http://civic.genome.wustl.edu - automatic!<br>[CIViC](http://civic.genome.wustl.edu) |
| **Blockquotes** | As Charles Darwin said:<br><br>>It is not the strongest<br>>of the species that survive,<br>>nor the most intelligent,<br>>but the one most responsive to change. |
| **Inline Code** | I think you should use an<br>`<addr>` element here instead. |

<table>
<tr><th colspan="3">Macros</th></tr>
<tr><td rowspan="4">@ Mention suggestions</td><td colspan="2">Type '@', and the first few letters of a user's name, and CIViC will show you a dropdown menu of users with matching display names. Hit enter to insert the user mention link, which will display in the rendered comment as a link to the user's profile page, and generate a notification to the mentioned user.</td></tr>
<tr><td>@username</td><td>Select any user in the CIViC community</td></tr>
<tr><td>@editors</td><td>Generate notification for all users that have editor roles</td></tr>
<tr><td>@admins</td><td>Generate notification for all users that have admin roles</td></tr>
<tr><td rowspan="7">#ENTITY link macro</td><td colspan="2">'#' followed by an entity type abbreviation, and an entity ID will be displayed as a link to that entity's summary view.</td></tr>
<tr><td>#V123</td><td>Variant link</td></tr>
<tr><td>#G123</td><td>Gene link</td></tr>
<tr><td>#E123</td><td>Evidence Item link</td></tr>
<tr><td>#VG123</td><td>VariantGroup link</td></tr>
<tr><td>#R123</td><td>Revision link</td></tr>
<tr><td>#A123</td><td>Assertion link</td></tr>
<tr><td rowspan="7">Macro suggestion prompts</td><td colspan="2">Type '#', followed by an entity type abbreviation, a colon and search string, and CIViC will show you a list of entities with matching names (or summaries, in the case of evidence items). For example, entering '#V:v600' will display a list of variants with 'V600' in the names; '#G:BR' will display a list of Genes with the string 'BR' in the name. Select an entity to insert an #ENTITY string for that entity.</td></tr>
<tr><td>#G:[string]</td><td>Search for a gene by name</td></tr>
<tr><td>#V:[string]</td><td>Search for a variant by name</td></tr>
<tr><td>#VG:[string]</td><td>Search for a variant group by name</td></tr>
<tr><td>#E:[string]</td><td>Search for an evidence item by summary</td></tr>
<tr><td>#R:[string]</td><td>Search for a revision by change contents</td></tr>
<tr><td>#A:[string]</td><td>Search for an assertion by summary</td></tr>
</table>

### References

1. Griffith M, Griffith OL, Coffman AC, Weible JV, McMichael JF, Spies NC, et al. DGIdb: mining the druggable genome. Nat Methods. 2013;10:1209–10.

2. Wagner AH, Coffman AC, Ainscough BJ, Spies NC, Skidmore ZL, Campbell KM, et al. DGIdb 2.0: mining clinically relevant drug–gene interactions. Nucleic Acids Res. Narnia; 2016;44:D1036–44.

3. Cotto KC, Wagner AH, Feng Y-Y, Kiwala S, Coffman AC, Spies G, et al. DGIdb 3.0: a redesign and expansion of the drug–gene interaction database. Nucleic Acids Res. Narnia; 2018;46:D1068–73.

4. Xin J, Mark A, Afrasiabi C, Tsueng G, Juchler M, Gopal N, et al. MyGene.info and MyVariant.info: Gene and Variant Annotation Query Services [Internet]. bioRxiv. 2015 [cited 2019 Apr 30]. p. 035667. Available from: https://www.biorxiv.org/content/10.1101/035667v1.abstract

5. Robarge JD, Li L, Desta Z, Nguyen A, Flockhart DA. The star-allele nomenclature: retooling for translational genomics. Clin Pharmacol Ther. 2007;82:244–8.

6. Tate JG, Bamford S, Jubb HC, Sondka Z, Beare DM, Bindal N, et al. COSMIC: the Catalogue Of Somatic Mutations In Cancer. Nucleic Acids Res. Narnia; 2019;47:D941–7.

7. Li MM, Datto M, Duncavage EJ, Kulkarni S, Lindeman NI, Roy S, et al. Standards and Guidelines for the Interpretation and Reporting of Sequence Variants in Cancer: A Joint Consensus Recommendation of the Association for Molecular Pathology, American Society of Clinical Oncology, and College of American Pathologists. J Mol Diagn. 2017;19:4–23.

8. Richards S, Aziz N, Bale S, Bick D, Das S, Gastier-Foster J, et al. Standards and guidelines for the interpretation of sequence variants: a joint consensus recommendation of the American College of Medical Genetics and Genomics and the Association for Molecular Pathology. Genet Med. 2015;17:405–24.
